# Chemical clues to infection: Metabolome differentiation underlies host colonization of potential biocontrol agents from the entomopathogenic genus *Cordyceps*

**DOI:** 10.1101/2025.07.06.659992

**Authors:** Esteban Charria-Girón, Rita Toshe, Artit Khonsanit, Noppol Kobmoo, Papitchaya Kwanthong, Tatiana E. Gorelik, Janet Jennifer Luangsa-ard, Sherif S. Ebada, Marc Stadler

## Abstract

*Cordyceps* species are widely recognized as entomopathogens, with some developed as biocontrol agents. These fungi produce bioactive metabolites contributing to their ecology and pathogenicity, yet their specific role during host infection remains poorly understood. To gain insights into how these fungi control their insect hosts, we investigated the metabolome and virulence traits of two potential biocontrol agents from the genus *Cordyceps*. Virulence assays on beet armyworms (*Spodoptera exigua*, Lepidoptera) revealed varying levels of pathogenicity, with *C. javanica* BCC 82944 exhibiting a higher virulence than *C. blackwelliae* BCC 37653, which revealed intermediate pathogenicity. Using state-of-the-art metabolomics, combined with 3D electron diffraction (3D ED) crystallography and comprehensive 1D/2D NMR spectroscopy, we identified diverse metabolites, including the cyclodepsipeptides beauverolides. *Cordyceps javanica* exhibited remarkable beauverolide diversity, featuring various amino acid rearrangements and fatty acid chain lengths, including three previously undescribed derivatives (**1**–**3**). While the main products of *C. blackwelliae* were diketopiperazines, feature-based molecular networking (FBMN) analysis uncovered the production of unprecedented beauverolides. To explore the functional relevance of these unique natural products, we analyzed the original insect cadavers from which each fungus was isolated. Our results revealed the presence of beauverolides and beauvericins in the host tissue, providing for the first time direct evidence of their involvement in fungal colonization during infection. Notably, not all beauverolides induced insect mortality *in vitro*, suggesting differentiated biological functions dependent on their amino acid organization. These findings indicate that distinct secondary metabolites may contribute to specific steps of the infection process. Moreover, the detection of species-specific metabolite profiles in the insect corpses suggests that *Cordyceps* species have developed chemically divergent infection strategies, possibly shaped by host specificity and ecological niche adaptation.

## Introduction

The genus *Cordyceps* (Cordycipitaceae, Hypocreales) encompasses fungi renowned for their ability to parasitize insects, a unique trait that defines their role as entomopathogens. Historically, fungi associated with insects and arachnids featuring apparent stromata were classified in the genus *Cordyceps*.^1^ However, successive taxonomic revisions based on molecular phylogenetics have led to the reclassification of these taxa into various families within Hypocreales, and several genera within the Cordycipitaceae, with up to 23 different genera currently accepted.^2–5^ These taxonomic advances have significantly broadened our understanding of the diversity within this family providing a robust foundation for the study of their ecological roles and derived applications. Based on the acquired information, a phylogenetic correlation could be found between certain genera and their host/substrate specificity, suggesting their ability to colonize and effectively target specific insect hosts.^6,7^ This unique parasitic behaviour has been leveraged to develop specific biocontrol agents useful for agricultural fields. Moreover, species from the genera *Cordyceps* and *Ophiocordyceps*, along with other medicinal fungi, have significant applications in traditional Chinese medicine, being used to treat various conditions such as respiratory, liver, and cardiovascular diseases, as well as low libido, impotence, hyperlipidemia, and chronic kidney disease.^8^ The global market for medicinal fungi in traditional Chinese medicine is estimated to be worth up to USD 1.5 billion.^9^

*Cordyceps* species are known as prolific producers of bioactive secondary metabolites, with many playing a role in their virulence and facilitate the colonization of host insects. These compounds might be involved in suppressing host defences or manipulating host physiology, yet their exact role in the infection process remains poorly understood.^10–12^ Despite the potential involvement of these metabolites in virulence, this aspect of *Cordyceps* chemical biology has been largely understudied. In contrast, these compounds might prove useful in the development of therapeutic treatments due to their multifaceted roles and diverse biological properties.^13,14^ In this study, we explored the metabolome and virulence traits of *Cordyceps javanica* and *C. blackwelliae*, two insect-associated fungi from Thailand selected for their distinct ecological roles and potential as biocontrol agents. By combining state-of-the-art metabolomics, classical chemical screening, and virulence assays, we aimed to identify key metabolites involved in the infection process and evaluate their suitability for pest control applications.

## Results and Discussion

### Virulence and Chemical Traits from Studied *Cordyceps* Strains

The virulence of three *Cordyceps* strains was evaluated against the beet armyworm (*Spodoptera exigua*, Lepidoptera). *Cordyceps blackwelliae* was originally isolated from a Lepidoptera pupa, while both *C. javanica* strains BCC 82944 and BCC 79245 were derived from Lepidoptera larvae. To assess the virulence, two types of mortality were considered: (1) mycelium-associated mortality (MM) representing the dead insects exhibiting fungal mycelia/spores on their surfaces, and (2) total mortality (TM) encompassing all deceased insects, regardless of fungal material presence. This distinction was made to provide deeper insights into the mechanisms of virulence. It is well known that hypocrealean entomopathogenic fungi grow and produce various secondary metabolites, particularly mycotoxins,^15,16^ inside the insect body to kill hosts. However, fungal growth and toxins production must be carefully balanced; premature hosts kill might not allow sufficient accumulation of necessary nutrients, hindering the fungi from full development to sporulation.^17^ Once the host insects are dead and colonized, these fungi sporulate and disperse, which should require appropriate conditions and lytic enzymes to break through the cuticles from within.^18^ Therefore, the presence of mycelial coverage might be linked to a moderate production of secondary metabolites and lytic enzymes, allowing an outward colonization by partially degrading insect tissues. Conversely, the absence of visible fungal growth on the surface of insect cadavers (as observed in TM) suggests the potential involvement of secondary metabolites that might have killed the insects early and acted systemically at a cellular level.

From the two *C. javanica* strains, BCC 82944 was more virulent, achieving 100% TM by the third day of experiment, whereas BCC 79245 caused an average TM of 80%, with both causing MM only around 20 to 40% (Figure 1A). *Cordyceps blackwelliae* BCC 37653 displayed an intermediate virulence compared to both other strains with TM up to around 90% the third day post-infection but failed to produce an observable fungal material even after seven days (Figure 1B). In summary, all strains demonstrated a high total mortality but with a limited development of observable mycelia covering insect corpses. However, *C. javanica* seemingly can produce more mycelia than *C. blackwelliae*. The early mortality suggests that the evaluated strains produce entomopathogenic secondary metabolites within the insect’s body, playing a specialized role in killing the beet armyworm while the differential mycelial development might be due to differences in the sets of metabolites produced specifically between both species.

**Figure 1.**
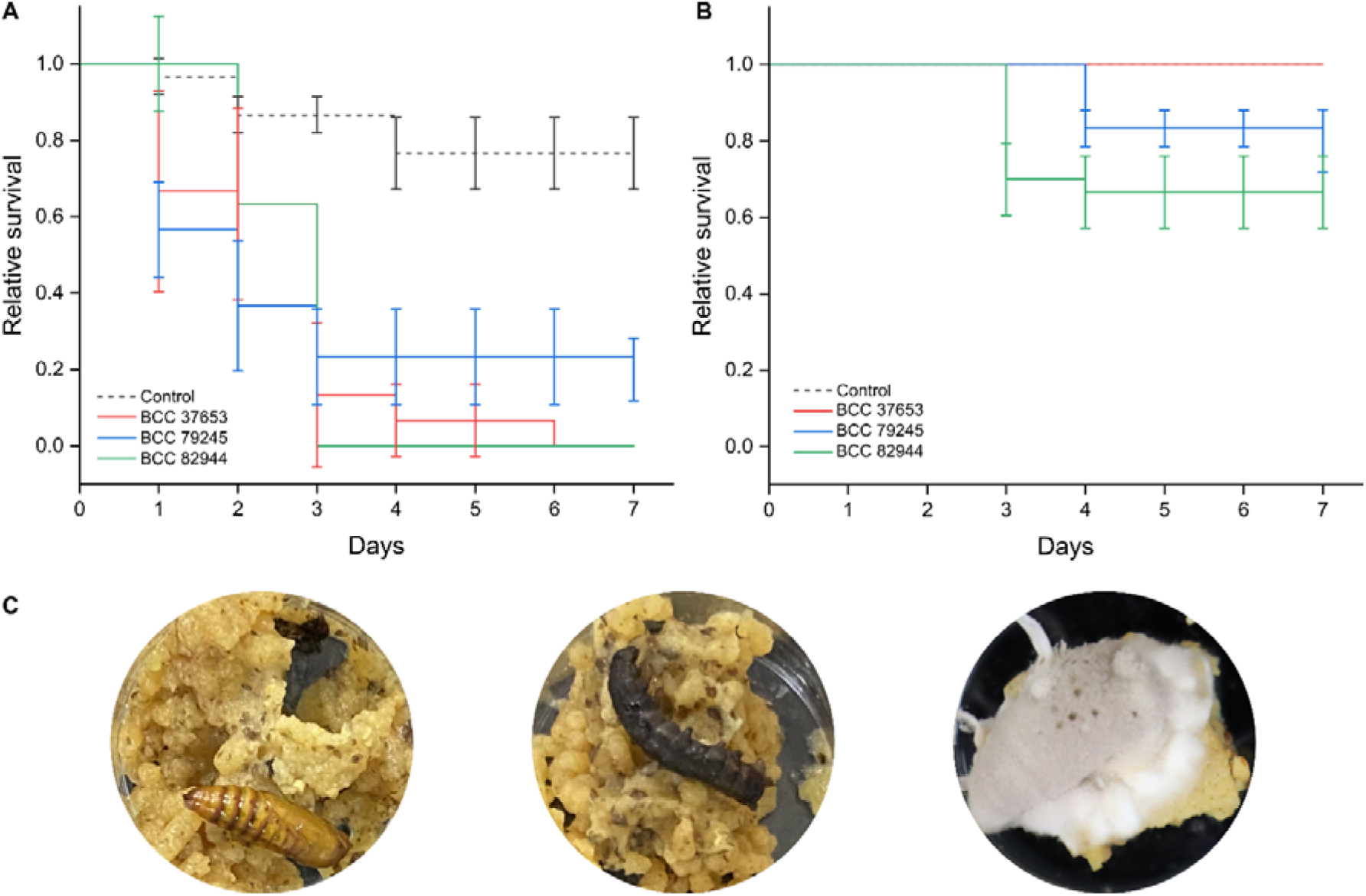
Virulence of *Cordyceps* spp., *C. blackwelliae* BCC 37653, *C. javanica* BCC 79245, and BCC 82944 against the beet armyworm (*Spodoptera exigua*, Lepidoptera) observed over a 7-day period. (**A**) Relative survival considering total mortality (TM). (**B**) Relative survival considering mycelium-associated mortality (MM). Error bars denote the standard error across three biological replicates. (**C**) From left to right: larva turned into pupa, deceased insect without visible mycelia, and deceased insect with visible mycelia.

Building on the hypothesis that secondary metabolites produced by these fungi play a role in the infection process, we decided to investigate their metabolome. However, it is important to note that the metabolites produced during infection may differ from those observed under axenic *in vitro* cultivation. To establish a baseline for comparison, each fungus was cultured in yeast malt (YM) liquid medium under axenic conditions until the available glucose was totally consumed. Three days after glucose depletion, secondary metabolites were extracted as described below in the Experimental section. The obtained crude extracts were analyzed using ultrahigh-performance liquid chromatography coupled with diode array detection and ion mobility tandem mass spectrometry (UHPLC-DAD-IM-MS/MS), and the resulting raw data files were pre-processed with MetaboScape software.^19^ Afterwards, features detected in the cultivation blank were excluded, and the resulting feature table was dereplicated against our in-house spectral library of fungal natural products. While *C. javanica* was expected to produce beauverolide-type cyclodepsipeptides and asperfurans,^13^ the secondary metabolites of *C. blackwelliae*, to the best of our knowledge, was not explored prior to this study.

To expand our understanding of the chemical space produced by both *Cordyceps* species, the natural product classes of the detected features were *de novo* predicted from their MS/MS spectra using CANOPUS.^20,21^ Hierarchical clustering analysis (HCA), combined with CANOPUS predictions, revealed clear differences in the secondary metabolite profiles of *C. blackwelliae* and *C. javanica* (Figure 2). While certain alkaloids and fatty acids were the most abundant chemical classes in extracts from *C. blackwelliae*, compounds classified as amino acids and peptides, including beauverolides, were the major metabolites in both *C. javanica* strains.

**Figure 2.**
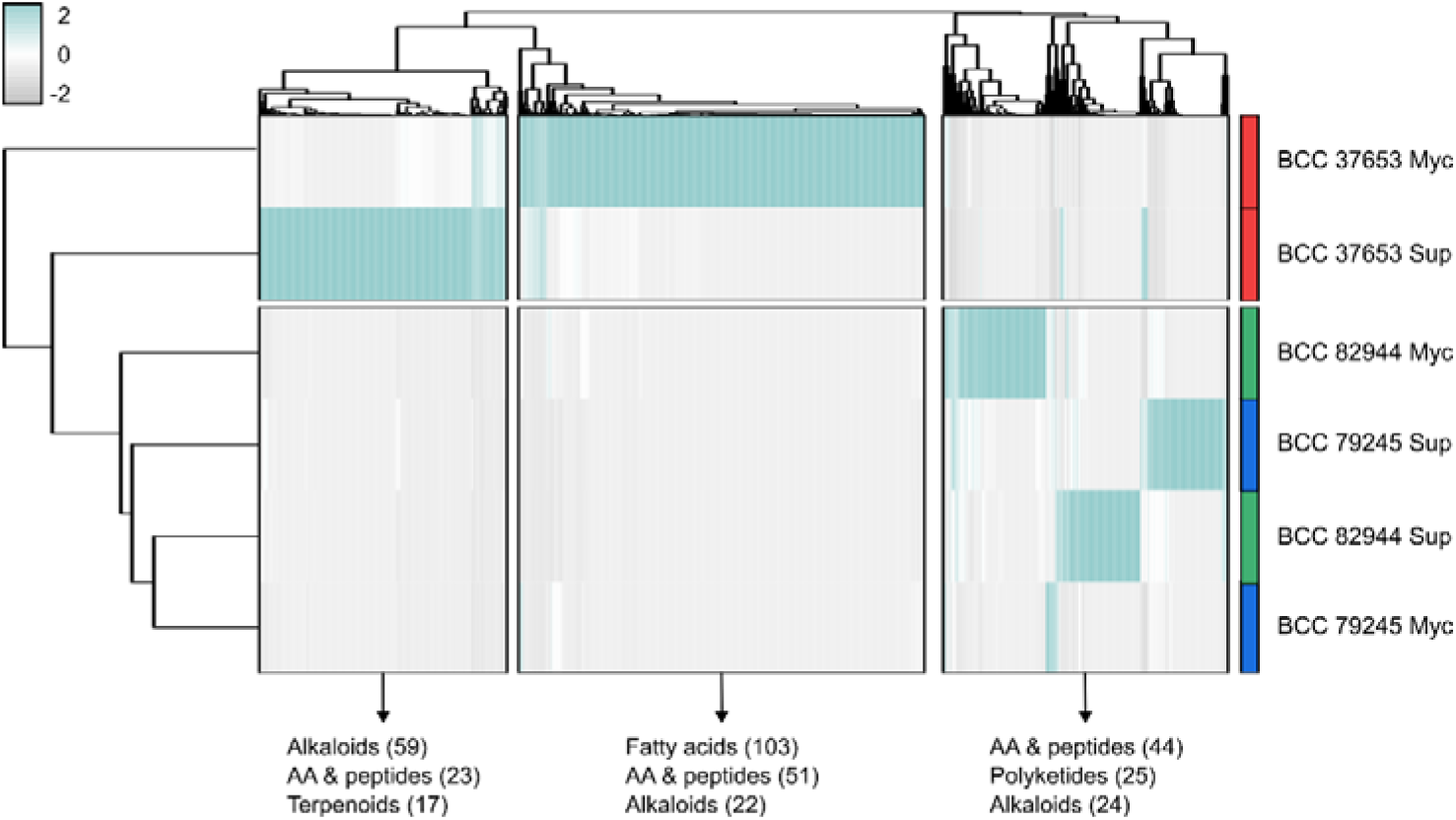
Heatmap displaying the hierarchical clustering of features detected from the cultivation of *Cordyceps blackwelliae* BCC 37653 and *C. javanica* (BCC 79245 and BCC 82944) in YM liquid medium. The relative abundance of features within the crude extracts is visualized, and the three most abundant natural product classes, as predicted by CANOPUS, are highlighted for the major clusters. The heatmap, along with dendrograms, was generated using the R package pheatmap.

The beauverolides I and J_b_, which were specifically identified in the extracts of the *C. javanica* strains through comparison with authentic standards, were not detected in *C. blackwelliae* whose main metabolites were classified as alkaloids, our feature-based molecular networking (FBMN) analysis revealed a broader diversity of beauverolides, beyond the annotated metabolites. Both *Cordyceps* species were found to produce beauverolide-like cyclodepsipeptides that were more prominent in *C. javanica*. However, *C. blackwelliae* produced distinct and unique analogs that were clustered together within the same molecular family (MF) as beauverolides I and J_b_. This observation suggests that, although the metabolome of *C. blackwelliae* is significantly more diversified than that of *C. javanica*, the production of beauverolides seems to be conserved between the two species. The beauverolides family was mainly characterized in *Beauveria bassiana* and known to play a role in the early pathogenesis.^22,23^ The conservation of this chemical family among the two *Cordyceps* species studied here support their common role in the early pathogenesis while the differences at the overall metabolome may, in fact, be related to the distinct virulence mechanisms of these two entomopathogenic fungi.

### Targeted Isolation and Structure Elucidation of Compounds

We hypothesize that beauverolide-type cyclic depsipeptides are the authentic metabolites associated with these fungi, as diketopiperazines are broadly produced across various fungal lineages and are not exclusive to entomopathogenic genera, making their role in infection less likely.^24^ Since both *C. javanica* strains produced beauverolides in higher titers than *C. blackwelliae*, we attentively evaluated beauverolide production in several fermentation media for BCC 82944 and BCC 79245 (see Experimental Section). Notably, *C. javanica* BCC 82944 exhibited a broader diversity of beauverolides, including some with previously unreported molecular formulas proposing novel structures. The obtained results revealed that GG1 medium supported the highest production. Despite that *C. blackwelliae* BCC 37653 predominantly produced diketopiperazines, we also scaled up its cultivation in YM medium to comprehensively profile its secondary metabolome. Chemical investigation of the crude extracts afforded seven beauverolide congeners including three previously undescribed. The crude extract of *C. blackwelliae* BCC 37653 afforded three diketopiperazine and two β-carboline derivatives (Figure 3).

**Figure 3.**
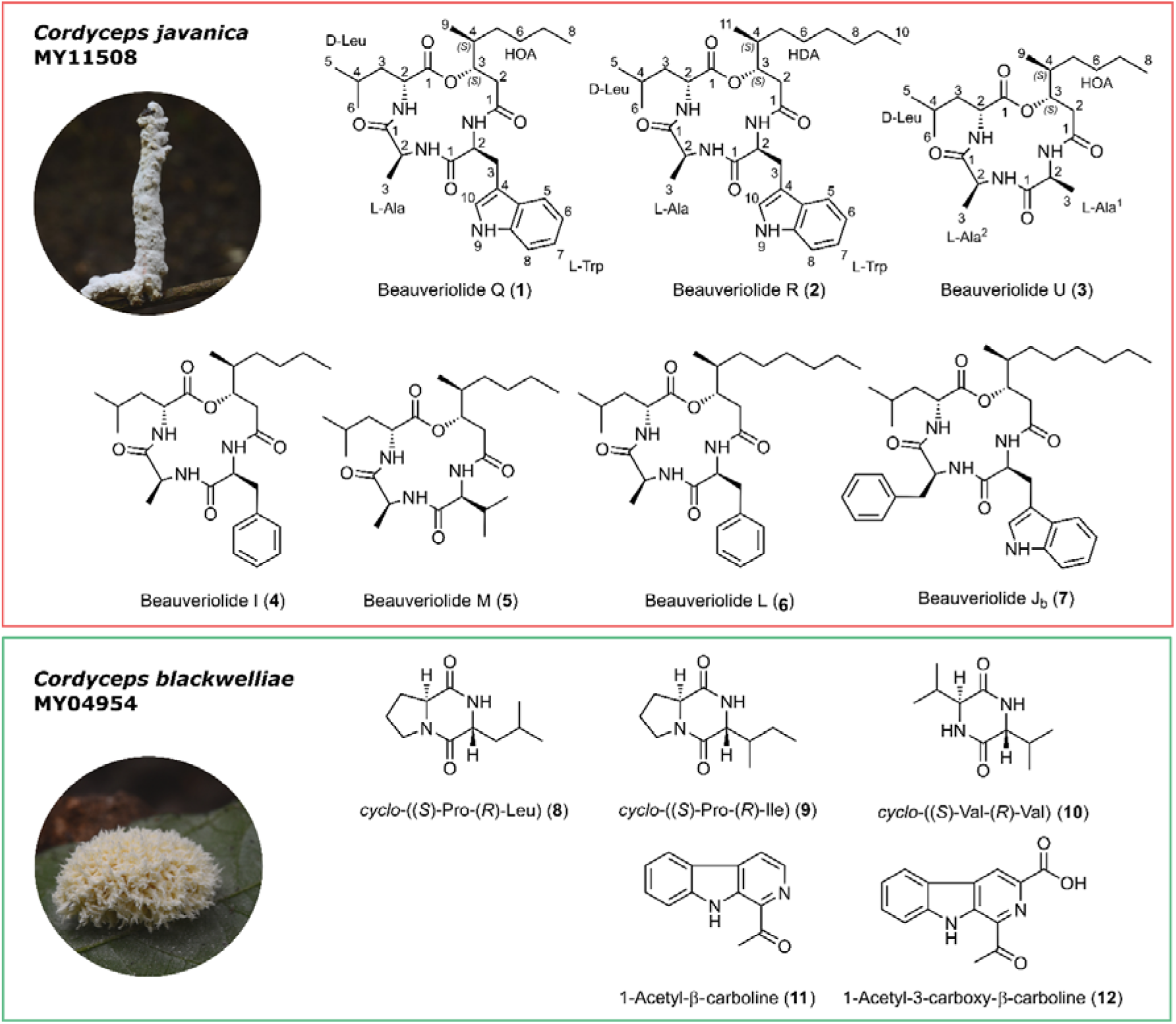
Chemical structures of the secondary metabolites isolated from *C. javanica* BCC 82944 (herbarium code MY11508) and *C. blackwelliae* BCC 37653 (herbarium code MY04954).

Compound **1** was obtained as a white powder, and its molecular formula was determined as C_29_H_42_N_4_O_5_ based on HR-ESI-MS data (Figure S2), showing a protonated molecular ion peak at *m/z* 527.3226 [M + H]^+^ and a sodium adduct at *m/z* 549.3046 [M + Na]^+^, indicating eleven degrees of unsaturation. Due to the poor solubility of **1** in deuterated solvents, NMR spectroscopy and attempts to grow suitable single crystals for X-ray diffraction were unsuccessful.

Therefore, 3D electron diffraction (3D ED) was employed as an alternative.^25^ Crystals of **1** had lateral dimensions in the micron range and were sufficiently thin to allow the collection of high-quality 3D ED data. Relevant crystallographic information can be found in the supplementary data. Once the crystal structure was determined, we targeted dynamical refinement with the aim of determining the absolute stereochemistry.^26^ The 3D ED data workflow is complex, involving multiple steps of data conversion. During this process, pattern flipping can occur, potentially leading to an incorrect determination of the absolute structure. To address this, we pre-calibrated the entire data workflow using GRGDS—a 5-amino-acid peptide with a known stereochemistry (Figure S3). Indeed, the workflow contained a pattern-flipping step, but once identified, this issue was resolved, enabling the reliable determination of the absolute stereochemistry of **1** (Figure 4).

**Figure 4.**
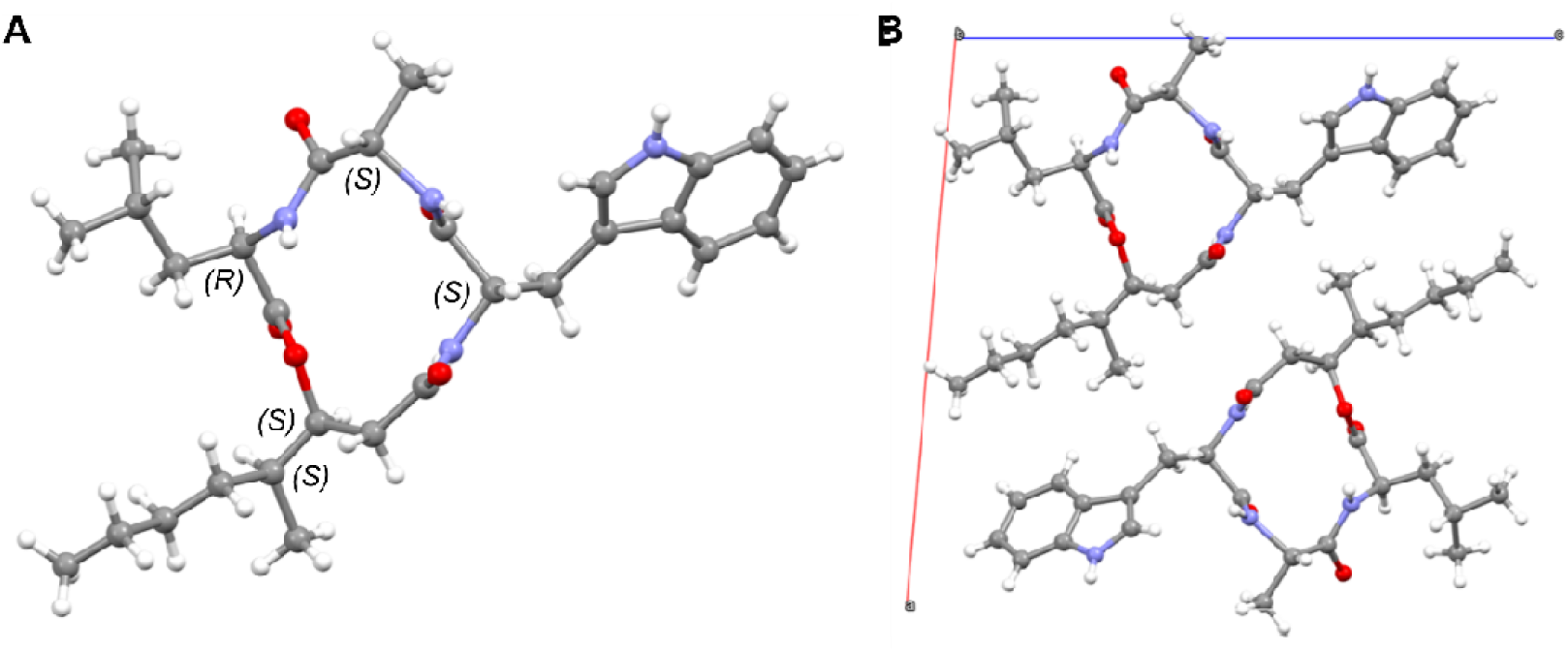
(**A**) Molecular conformation and absolute stereochemistry of beauverolide Q (**1**) as determined from 3D ED. (**B**) Molecular packing within the crystal structure viewed along the b axis.

To our surprise, during the preparation of our manuscript, a crystal structure and the absolute stereochemistry of a similar compound, beauverolide I (**4**), was published,^27^ also determined from 3D ED data. Interestingly, despite the different aromatic substituents and, consequently, distinct crystal structures, the macrocycle and aliphatic parts of the two molecules adopt nearly identical conformations. This suggests a relative rigidity of the macrocycle. An overlay of the two molecular conformations—**1** and beauverolide I (**4**)—is shown in Figure S4.

Based on the obtained results and by searching the reported literature, compound **1** was unambiguously determined to a previously undescribed beauverolide derivative featuring *cyclo*-(–C_9_–*L*-Trp–*L*-Ala–*D*-Leu) named beauverolide Q, whose molecular formula was tentatively reported in a chemotaxonomic study of the genera *Beauveria* and *Paecilomyces*.^28^ It was mentioned as a synthetic chemical among many others in a Japanese patency as acyl-CoA:cholesterol acyltransferase 2 (ACAT-2) inhibitors with no spectral data included.^29^ Compound **2** was obtained as a white powder whose HR-ESI-MS (Figure S7) established its molecular formula as C_31_H_46_N_4_O_5_ indicating eleven degrees of unsaturation similar to **1**. Comparison of the molecular formulas of **1** and **2**, revealed that **2** has an additional C_2_H_4_ moiety than **1**, as indicated by a mass difference of 28 Da. Unlike **1**, compound **2** was soluble in deuterated DMSO and therefore, its NMR spectral data (Table 1) could be obtained. The ^1^H NMR spectral data of **2** revealed the pattern of a peptide containing three N*H* proton signals (δ_H_ 7.19 ∼ 8.42 ppm) and four α-protons at δ_H_ 3.90 ∼ 4.39 ppm indicating the presence of three amino acid residues. In addition, the ^1^H NMR spectral data of **2** revealed an oxygenated multiplet proton at δ_H_4.84 (dt, *J* = 10.2, 5.3 Hz) that is directly correlated to an oxygenated sp^3^ carbon (δ_C_ 75.8).

**Table 1.**
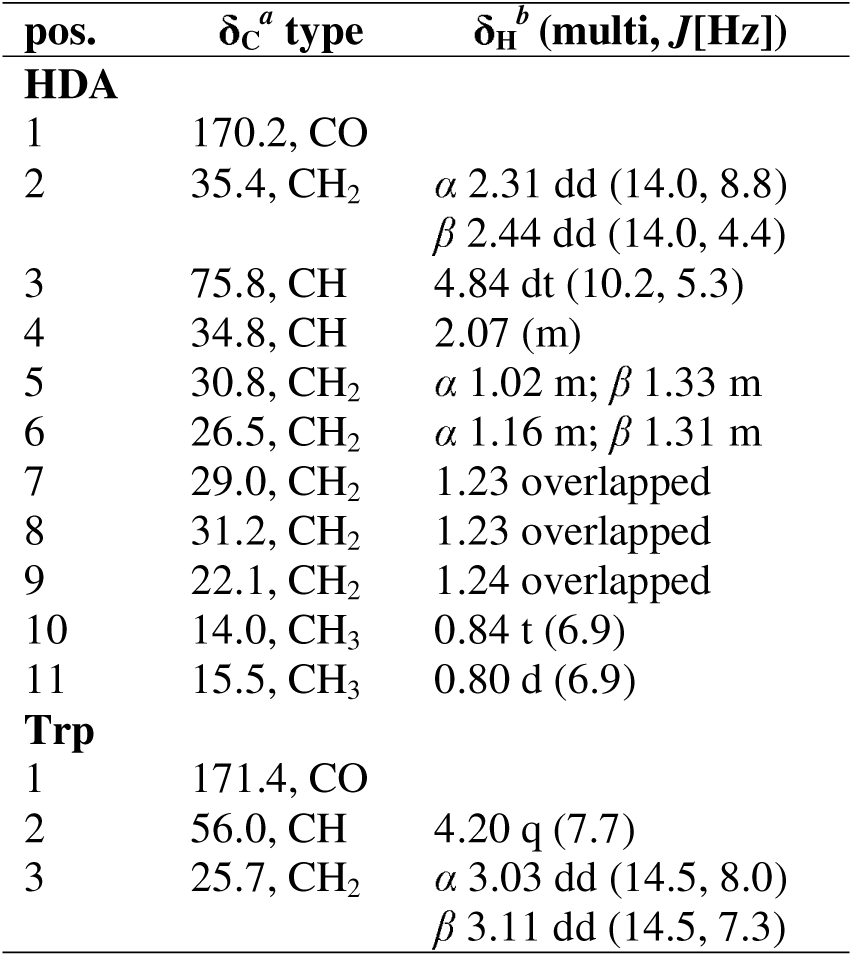
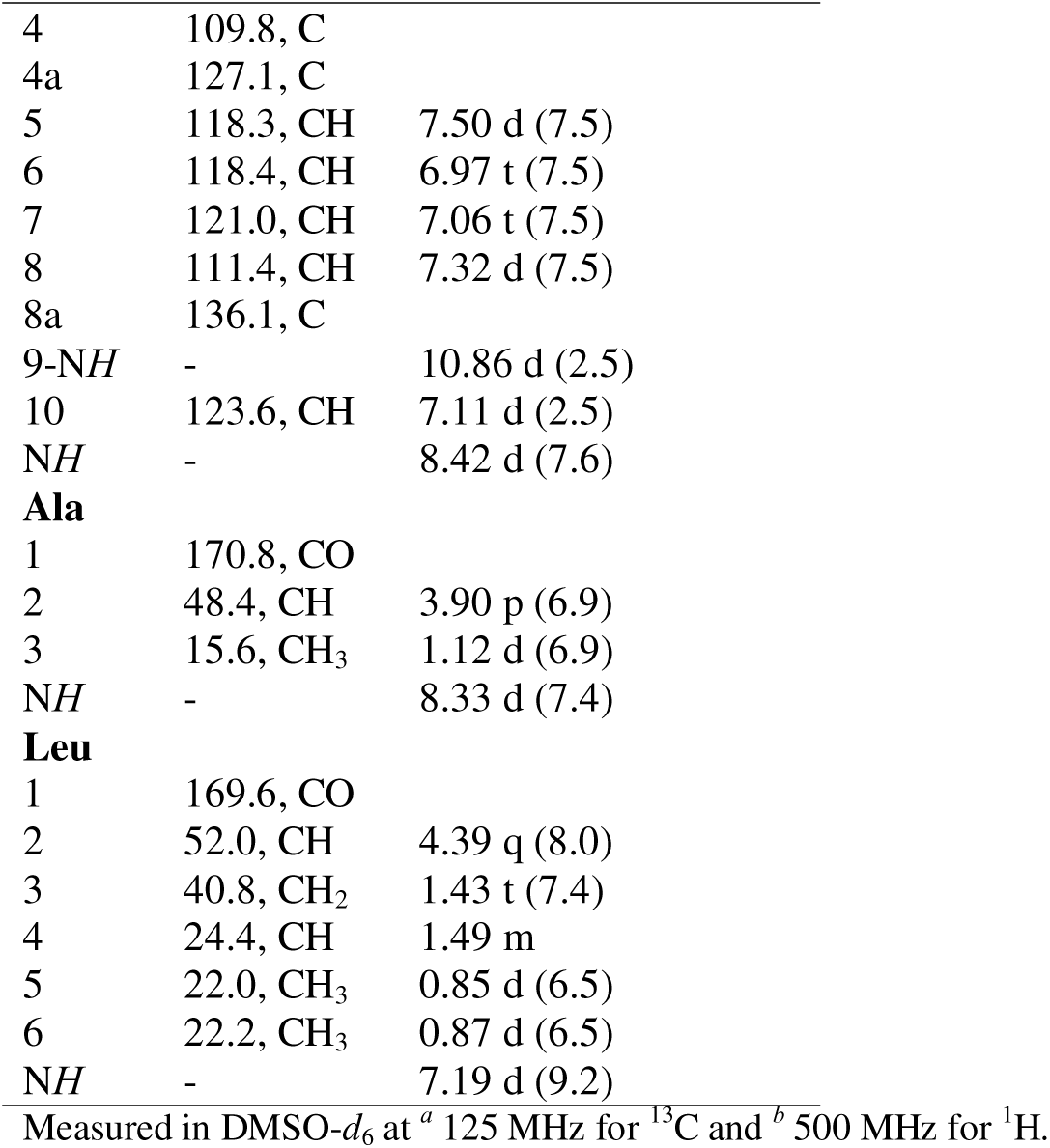
1D (^1^H and ^13^C) NMR data of **2**.

| pos. | $\delta_C^a$ type | $\delta_H^b$ (multi, $J$[Hz]) |
| --- | --- | --- |
| **HDA** |  |  |
| 1 | 170.2, CO |  |
| 2 | 35.4, $CH_2$ | $\alpha$ 2.31 dd (14.0, 8.8)<br>$\beta$ 2.44 dd (14.0, 4.4) |
| 3 | 75.8, CH | 4.84 dt (10.2, 5.3) |
| 4 | 34.8, CH | 2.07 (m) |
| 5 | 30.8, $CH_2$ | $\alpha$ 1.02 m; $\beta$ 1.33 m |
| 6 | 26.5, $CH_2$ | $\alpha$ 1.16 m; $\beta$ 1.31 m |
| 7 | 29.0, $CH_2$ | 1.23 overlapped |
| 8 | 31.2, $CH_2$ | 1.23 overlapped |
| 9 | 22.1, $CH_2$ | 1.24 overlapped |
| 10 | 14.0, $CH_3$ | 0.84 t (6.9) |
| 11 | 15.5, $CH_3$ | 0.80 d (6.9) |
| **Trp** |  |  |
| 1 | 171.4, CO |  |
| 2 | 56.0, CH | 4.20 q (7.7) |
| 3 | 25.7, $CH_2$ | $\alpha$ 3.03 dd (14.5, 8.0)<br>$\beta$ 3.11 dd (14.5, 7.3) |
| 4 | 109.8, C |  |
| 4a | 127.1, C |  |
| 5 | 118.3, CH | 7.50 d (7.5) |
| 6 | 118.4, CH | 6.97 t (7.5) |
| 7 | 121.0, CH | 7.06 t (7.5) |
| 8 | 111.4, CH | 7.32 d (7.5) |
| 8a | 136.1, C |  |
| 9-NH | - | 10.86 d (2.5) |
| 10 | 123.6, CH | 7.11 d (2.5) |
| NH | - | 8.42 d (7.6) |
| **Ala** |  |  |
| 1 | 170.8, CO |  |
| 2 | 48.4, CH | 3.90 p (6.9) |
| 3 | 15.6, CH<sub>3</sub> | 1.12 d (6.9) |
| NH | - | 8.33 d (7.4) |
| **Leu** |  |  |
| 1 | 169.6, CO |  |
| 2 | 52.0, CH | 4.39 q (8.0) |
| 3 | 40.8, CH<sub>2</sub> | 1.43 t (7.4) |
| 4 | 24.4, CH | 1.49 m |
| 5 | 22.0, CH<sub>3</sub> | 0.85 d (6.5) |
| 6 | 22.2, CH<sub>3</sub> | 0.87 d (6.5) |
| NH | - | 7.19 d (9.2) |

Measured in DMSO-*d*<sub>6</sub> at <sup>a</sup> 125 MHz for <sup>13</sup>C and <sup>b</sup> 500 MHz for <sup>1</sup>H.

The ^1^H NMR and ^1^H–^1^H COSY spectral data of **2** (Figures 5, S8 and S10) revealed the presence of a tryptophan residue (Trp) supported by the spin system extending among the characteristic four adjacent aromatic protons (δ_H_ 6.97 ∼ 7.50 ppm) and a doublet aromatic proton at δ_H_ 7.11 (d, *J* = 2.5 Hz) correlated via ^1^H–^1^H COSY cross peak to an exchangeable N*H* proton at δ_H_ 10.86 (d, *J* = 2.5 Hz) of the indole ring.

**Figure 5.**
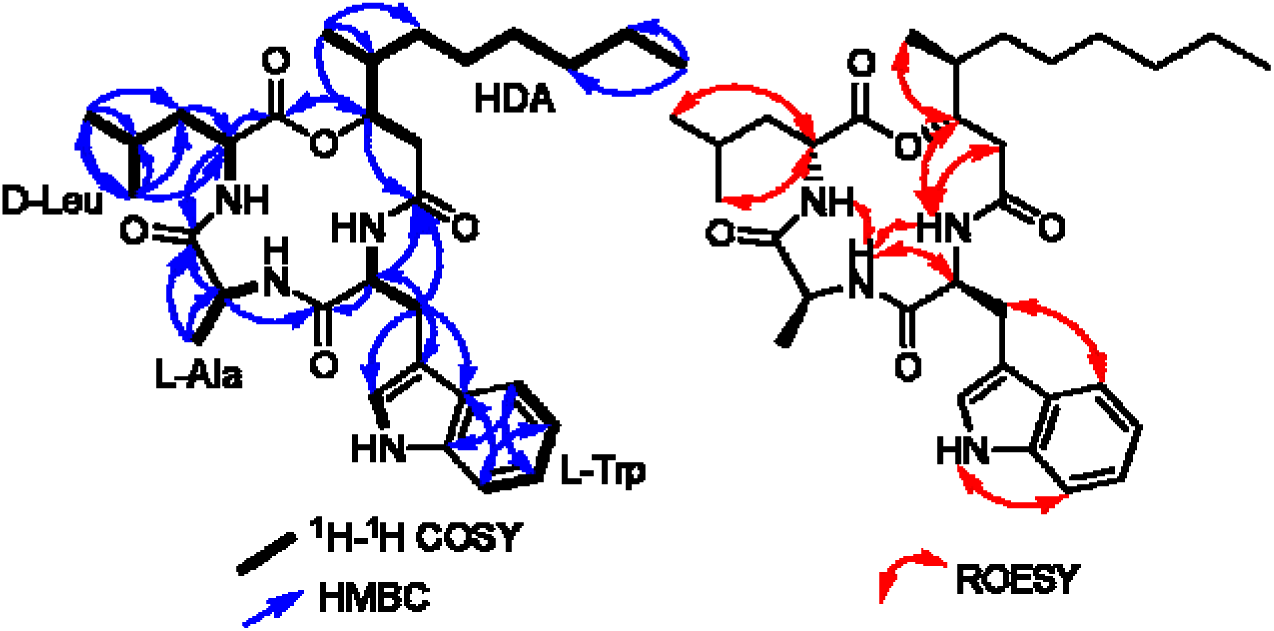
Key ^1^H-^1^H COSY, HMBC and ROESY correlations of **2**.

A careful investigation of ^1^H NMR and ^1^H–^1^H COSY spectra (Table 1, Figure 5) suggested the presence of two amino acid residues namely; alanine (Ala) and leucine (Leu) supported by two spin systems as follows: 1) from an exchangeable N*H* proton at δ_H_ 8.33 (d, *J* = 7.4 Hz) to an α-proton at δ_H_ 3.90 (p, *J* = 6.9 Hz) and a doublet methyl group at δ_H_ 1.12 (d, *J* = 6.9 Hz); 2) from an exchangeable N*H* proton at δ_H_ 7.19 (d, *J* = 9.2 Hz) to an α-proton at δ_H_ 4.39 (q, *J* = 8.0 Hz), a triplet methylene group at δ_H_ 1.43 (t, *J* = 7.4 Hz) and a multiplet methine proton at δ_H_ 1.49 ending by two doublet methyl group at δ_H_ 0.85/0.87 (d, *J* = 6.5 Hz). A literature search based on these results confirmed the identity of **2** as a beauverolide derivative, specifically beauverolide R, previously reported in a chemotaxonomic study of the related genera. However, no detailed spectral data other than tandem mass spectrometry had been reported previously.^28^ In addition to the three amino acid residues, Trp–Ala–Leu, and according to the molecular formula, the fourth moiety was deduced to be 3-hydroxy-4-methyldecanoic acid (HDA). The amino acid sequence was confirmed by HMBC and ROESY spectra of **2** (Figure 5) indicating its amino acid sequence as *cyclo*-(–C_11_–Trp–Ala–Leu) evidenced by the revealed key correlations. Attempts to assign the absolute configurations of the amino acids sequence using Marefy’s method were unsuccessful despite being repeated twice. However, based on the shared biosynthetic origin, supported by previous literature, and the available crystal structure of **1** from this study, the absolute configurations were assigned as *cyclo*-(–C_11_–*L*-Trp–*L*-Ala–*D*-Leu), consistent with other beauverolides. Based on the obtained NMR spectral data, compound **2** was elucidated as a previously undescribed cyclodepsipeptide, beauverolide R.

Compound **3** was obtained as a colorless oil. The HR-ESI-MS analysis (Figure S15) determined its molecular formula as C_21_H_37_N_3_O_5_, revealing a protonated molecular ion peak and a sodium adduct at *m/z* 412.2809 [M + H]^+^ and 434.2625 [M + Na]^+^, respectively, and indicating five degrees of unsaturation. Similar to **1**, this metabolite was insoluble in deuterated solvents and due to its oily nature, all crystallographic approaches, including 3D electron diffraction, were unsuccessful for structure determination. However, these compounds, comprising one 3-hydroxy-4-methyl fatty acid, two *L*-amino acids, and one *D*-amino acid, exhibit distinct MS fragmentation patterns. Consequently, we conducted an in-depth MS/MS analysis of compound **3**. In our MS/MS experiments, beauverolide I (**4**), also isolated during this study, clustered together with compound **3** in our FBMN analysis. Compound **4** has a molecular mass of 487.3043 Da and comprises phenylalanine, alanine, leucine, and a 3-hydroxy-4-methyl octanoyl residue. The 76 Da mass difference between the two compounds corresponds to the loss of a C_6_H_4_ group, which indicates the substitution of *L*-Phe with *L*-Ala in compound **3** (Figure S5). Similarly, compound **3** clustered as well with beauverolide M (**5**), which has a molecular mass of 439.3048 Da and features *L*-Val instead of *L*-Phe. Both compounds generated common fragment ions at *m/z* 341.24, 281.19, 228.16, 210.15, 156.14, 139.12, and 132.11. However, the 28 Da difference between the two metabolites can be attributed to a C_2_H_4_ group, indicating the substitution of *L*-Val with *L*-Ala in compound **3** (Figure S5). Altogether, this evidence supports the identification of **3** as a previously undescribed derivative that was given a trivial name beauverolide S.

Three additional beauverolides, M (**5**), L (**6**), and J_b_ (**7**), were identified by comparison of their HR-ESI-MS and NMR data with previously reported literature.^13,28,30,31^ In addition, chemical investigation of the crude extract derived from *C. blackwelliae* BCC 37653 in YM medium afforded five known compounds. These were identified as diketopiperazines, consistent with our MS/MS-based analysis, and were elucidated as *cyclo*-((*S*)-Pro-(*R*)-Leu) (**8**), *cyclo*-((*S*)-Pro-(*R*)-Ile) (**9**) and *cyclo*-((*S*)-Val-(*R*)-Val) (**10**), 1-acetyl-β-carboline (**11**) and 1-acetyl-3-carboxy-β-carboline (**12**) by comparisons of their HR-ESI-MS and NMR spectral data with the reported literature.^32–36^

### Biological Properties of Secondary Metabolites and Ecological Relevance

To investigate the biological properties of the isolated compounds, they were subjected to antimicrobial assay against several bacterial and fungal pathogens, including Gram-positive and Gram-negative bacteria, filamentous fungi, and yeasts (see Experimental section). The obtained results (Table S3) revealed that beauverolides S (**3**) and I (**4**) exhibited weak activity against *Staphylococcus aureus* (MIC = 66.6 µg/mL), with **4** also inhibited *Bacillus subtilis* at the same concentration. Beauverolide M (**5**) exhibited weak activity against *Mucor hiemalis*. Interestingly, none of the metabolites isolated from *C. blackwelliae* exhibited antimicrobial activity against the tested microorganisms.

All compounds were evaluated for their cytotoxic effects against two mammalian cell lines, with those showing strong activity subjected to further testing against five additional cell lines. Among the tested compounds, beauverolides I (**4**) and J_b_ (**7**) (Table S4) displayed cytotoxic effects against human endocervical adenocarcinoma (KB3.1), with IC_50_ values of 13.3 and 36.5 µM, respectively. Compound **4**, however, was further evaluated against other cell lines, displaying potent cytotoxicity against human lung carcinoma (A549), human epidermoid carcinoma (A431), and human prostate carcinoma (PC-3) at IC_50_ values of 0.9, 1.2 and 3.5 µM, respectively. While in the case of the compounds from *C. blackwelliae*, no cytotoxic effect was observed against the tested cell lines. The obtained results of the antimicrobial and cytotoxic assays for compounds **1**–**12** are summarized in supplementary material (Tables S3 and S4).

Our findings suggest that beauverolides, while not broadly antimicrobial, may play a more specialized role in the entomopathogenic lifestyle of *Cordyceps*. Their activity against specific bacterial and fungal strains might reflect a dual role in modulating microbial competitors within the insect host and weakening host defense mechanisms by targeting its microbiome, as it has been demonstrated in the case of helvolic acid in *Metarhizium* spp.^11^ Furthermore, the cytotoxic effects of beauverolides, particularly beauverolide I (**4**), might point towards toxicity to insects, thereby contributing to virulence. Although metabolites from *C. blackwelliae* showed no antimicrobial or cytotoxic activities in our assays, it is worth noting that FBMN analysis underscored the presence of beauverolide-like compounds in the extracts from this fungus. Unfortunately, these compounds could not be isolated in sufficient purity or quantity for structural and biological characterization, leaving this as an interesting avenue for future studies.

Given the observed cytotoxic and antimicrobial activities during in vitro assays, we proceeded to evaluate the virulence of beauverolide I (**4**) in a more ecologically relevant context. For this purpose, we established a virulence assay to evaluate individual compounds against the beet armyworm, following a similar protocol to the spore suspension assays (see Experimental section). For this assay, only the TM was considered, since mycelial growth was not expected. As a positive control, the well-characterized mycotoxin cytochalasin D was selected for its ability to disrupt the eukaryotic actin cytoskeleton, as well as the fact that it is widely produced by entomopathogenic fungi of the genus *Metarhizium*.^12^ For proof-of-concept testing, beauverolides I (**4**) and M (**5**) were selected due to their availability in sufficient amounts for bioassays. Beauverolide I (**4**) displayed significant virulence, causing an average mortality of 33% during the first five days, increasing to 67% TM in the last two days of the experiment (Figure 6). In contrast, beauverolide M (**5**), differs from **4** in having *L*-Val replacing *L*-Phe, showed no insecticidal activity in beet armyworms, even after 7 days, and demonstrated lower TM than the two negative controls. Meanwhile, the treatment with cytochalasin D resulted in 83% TM from the second day (Figure 6A). These findings highlight the potential of beauverolide I (**4**) as an effector molecule in the entomopathogenicity of *Cordyceps* spp. The observed mortality in beet armyworms aligns with its cytotoxic effects against mammalian carcinoma cell lines, reinforcing its role as a bioactive compound with significant insecticidal potential. Interestingly, beauverolide M (**5**), despite being structurally similar to **4**, showed no virulence against the beet armyworm, suggesting that even small structural changes can dramatically alter the biological activity of these compounds, and their production might play a different role during infection.

**Figure 6.**
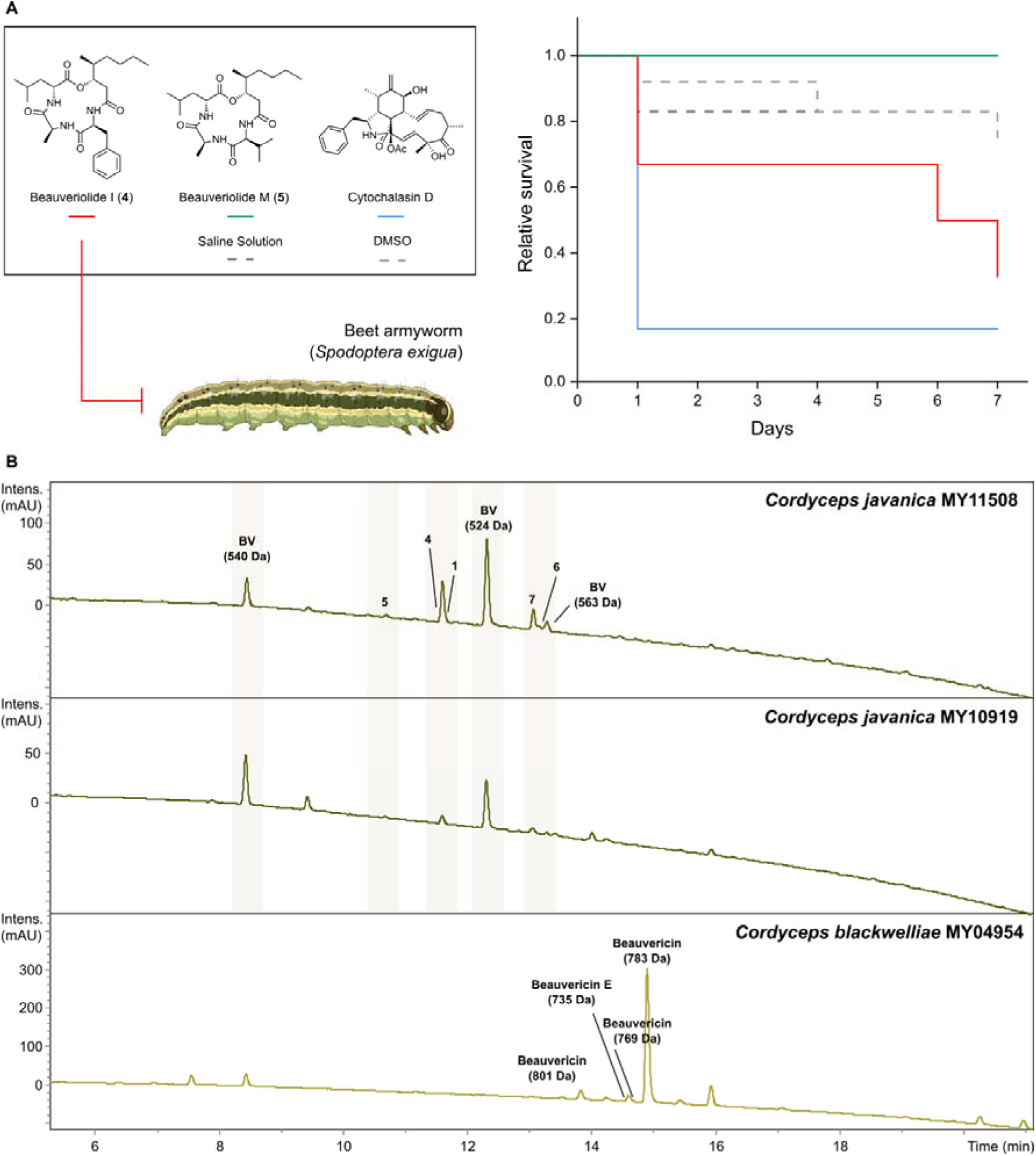
**(A)** Evaluation of the virulence of beauverolides I (**4**) and M (**5**) produced by *C. javanica* against the beet armyworm (*Spodoptera exigua*, Lepidoptera) over a 7-day period, considering the relative survival rate based on total mortality (TM). Cytochalasin D served as the positive control, while saline solution (8.5% NaCl) and DMSO were used as negative controls. **(B)** HPLC-UV/Vis chromatograms (210 nm) of the crude extracts obtained from the insect host corpses from *C. javanica* strains MY11508 and MY10919, and *C. blackwelliae* strain MY04954 herbarium material. Isolated metabolites are indicated by bold compound numbers. Putative beauverolide derivatives are labeled as BV followed by their corresponding molecular weight. Similarly, detected beauvericins are labeled with their name and molecular weight.

Our study also underscores the relevance of specific amino acid substitutions in modulating the biological properties of beauverolides. Modifying the *L*-Trp residue in beauverolide Q (**1**) with *L*-Phe, *L*-Ala, or *L*-Val results in distinct biological activities, including altered cytotoxicity and virulence profiles. This indicates that the amino acid at this certain position is crucial for both antimicrobial and insecticidal properties. The lack of antimicrobial activity in some beauverolides, such as those containing HDA, further supports the hypothesis that these molecules may have different functions, such as facilitating insect pathogenicity rather than acting broadly against microbial competitors. To the best of our knowledge, beauverolides exhibit selective biological activities that depend on their amino acid configuration, as demonstrated by beauverolide I (**4**), which showed insecticidal effects and strong cytotoxicity against carcinoma cell lines, with no apparent toxicity toward mouse fibroblast cells. In fact, beauveriolides I and III, of which the first shares the same chemical structure as compound **4**, have previously been reported to exhibit potent antiatherogenic activity in macrophages, while showing no adverse effects in animal studies.^37^ Additionally, beauverolides have been detected during the late stages of silkworm larval infection by *Beauveria bassiana*, as revealed by untargeted metabolomics analyses.^38^ Although the precise mechanisms underlying their selective *in vitro* activity remain unresolved and likely differ from those involved in insect pathogenesis, our findings support the non-toxic profile of metabolites from both *C. javanica* and *C. blackwelliae*. This represents a valuable step toward evaluating their potential use as biocontrol agents.

Finally, to determine whether the natural products identified *in vitro* are also produced during host infection, we examined the original insect cadavers from which both *Cordyceps* strains were isolated. Organic extraction of the authentic herbarium material revealed the presence of beauverolides in *C. javanica* associated insect tissue and beauvericins in that of *C. blackwelliae* (Figure 6B). This provides, for the first time, direct evidence that these secondary metabolites are produced *in vivo* in their natural hosts and confirms their likely involvement during colonization. These findings reinforce the ecological relevance of our *in vitro* observations and support the hypothesis that secondary metabolites contribute to virulence, either through direct insecticidal activity or by modulating microbial competitors within the host.

The distinct metabolomes observed, *C. javanica* producing beauverolides both *in vitro* and in infected tissues, while *C. blackwelliae* produces the widespread diketopiperazines *in vitro* but beauvericins in insect corpses, may reflect divergent ecological strategies or host-specific adaptations, suggesting species-specific metabolic and regulatory programs shaped by evolutionary pressures. Notably, both *Cordyceps* species denote significant taxonomic distance among them, which results more evident in the underlying chemical differentiation observed, however, it is an open question whether this difference is a unique scenario or there is a broader pattern that can be observed at the genus or family level and this will be the aim of future studies. Despite the clear functional implications of the detected compounds, we believe that future investigation must embark into the spatial and temporal dynamics of these molecules during infection, as well as their molecular targets and regulatory mechanisms. Altogether, our work highlights the potential of *Cordyceps* species as rich sources of structurally and functionally diverse secondary metabolites, advancing our understanding of fungal entomopathogenicity and informing the development of more effective and ecologically sound biocontrol agents.

## EXPERIMENTAL SECTION

### General Experimental Procedures

Optical rotations were measured at 20 °C using an MCP-150 polarimeter (Anton-Paar Opto Tec GmbH, Seelze, Germany). UV/Vis spectra were obtained with a UV-2450 spectrophotometer (Shimadzu, Kyoto, Japan). NMR spectra were acquired using an Avance III 500 spectrometer (Bruker, Billerica, MA, USA; ^1^H NMR at 500 MHz, ^13^C NMR at 125 MHz), with compounds dissolved in DMSO-*d*_6_. High-resolution electrospray ionization mass spectra (HR-ESI-MS) were obtained using an Agilent 1200 Infinity Series HPLC-UV system (Agilent Technologies, Santa Clara, CA, USA) with a C18 Acquity UPLC BEH column (50 × 2.1 mm, 1.7 μm; Waters, Milford, MA, USA). The mobile phases consisted of solvent A (deionized water with 0.1% formic acid) and solvent B (acetonitrile with 0.1% formic acid). The gradient elution was set from 5% B for 0.5 min, increasing to 100% B over 19.5 min, holding at 100% B for 5 min, at a flow rate of 0.6 mL/min, with UV/Vis detection at 190-600 nm. The system was connected to a time-of-flight mass spectrometer (ESI-TOF-MS, maXis, Bruker, Billerica, MA, USA; scan range 100-2500 *m/z*, rate 2 Hz, capillary voltage 4500 V, dry temperature 200 °C).

### Fungal Material

*Cordyceps blackwelliae* (BCC37653) was isolated from lepidopteran pupae found on the underside of leaves on 19 July 2019 from Khao Yai National Park, Nakhon Ratchasima Province, Thailand (14°26’20.72’N, 101°22’20.02’E). The pure culture was deposited in the BIOTEC culture collection (BCC). Fungal specimens were dried in a food dehydrator and deposited in the BIOTEC Bangkok Herbarium (BBH), Thailand Science Park, Pathum Thani Province, Thailand. Combined-loci phylogenetic analyses based on ITS and EF1 sequences (GenBank accession numbers: *ITS* = PP709053, *EF1* = PP735442) confirmed that BCC 37653 was nested with the type species of *C. blackwelliae* TBRC 7257.^39^ *Cordyceps javanica* strains (BCC 82954 and BCC 79245) were acquired from the BIOTEC Culture Collection (BCC) and reported previously.^13^

### Virulence Assays

The virulence of two strains of *Cordyceps javanica* (BCC 82944 and BCC 79245) and one of *C. blackwelliae* (BCC 37653) was evaluated against the beet armyworm (*Spodoptera exigua*, Lepidoptera). First, the strains were grown on PDA at 25 °C for varying durations (ranging from one week to one month) until sporulation occurred. Spores from each strain were collected into 1 mL of sterilized water, counted, and then diluted to a concentration of 10^8^ spores/mL. For the virulence assays, a 3-µL spore suspension aliquot was injected into each beet armyworm, with thirty insects divided into three replicates of ten individuals each per strain. Mortality was monitored daily for one week, categorized into two types of mortality: (1) unconditional mortality, where any dead insect was counted as non-survival, and (2) mortality with mycelia, where only insects that died and were covered with fungal mycelia were considered non-survival. This categorization distinguished general virulence from reproductive success, with the mortality rate calculated as the number of dead insects (either with or without fungal mycelia) divided by the total number of ten insects in each replicate.

To evaluate the virulence of pure compounds on beet armyworms, beauverolides I (**4**) and M (**5**) were selected due to their sufficient isolated quantities at the time of testing, with cytochalasin D serving as a positive control. The compounds were first dissolved in DMSO and subjected to serial dilution to achieve a concentration range from 40 µg/µL to 0.625 µg/µL. Beet armyworms were injected with 0.5 µL of each dilution, with six worms tested per concentration to assess toxicity over 7 days. DMSO was used as a solvent due to its efficacy in dissolving the compounds, whereas saline and DMSO alone were injected as negative controls. Following injection, the worms were incubated in medium at 25 °C for 7 days, with their viability was monitored daily.

### Metabolomics Studies

To evaluate the production of secondary metabolites by strains of *C. blackwelliae* and *C. javanica*, each fungus cultured in yeast malt liquid medium (YM 6.3; 10 g/L malt extract, 4 g/L D-glucose, 4 g/L yeast extract, pH 6.3 before autoclaving). Each fungal strain was grown on YM agar and after sufficient growth, five plugs were excised using a cork borer (7 mm diameter) and transferred to flasks containing 100 mL of SMYA medium, which were incubated at 25 °C with shaking at 220 rpm for 7 days. For metabolite screening, 3 mL of the respective seed cultures were added to 500 mL flasks containing 200 mL of the liquid medium and incubated at 23 °C and 140 rpm. Glucose levels were monitored daily, and cultures were harvested three days post-glucose depletion. The supernatant and mycelium were separated via vacuum filtration and processed separately. The supernatant was extracted twice with ethyl acetate, and the obtained organic phases were combined and evaporated under reduced pressure. The mycelia were soaked in acetone, ultrasonicated, and filtered. The acetone solution was evaporated, dispersed in water, and processed in a similar manner as the supernatant.

After detecting the production of diverse secondary metabolites by the evaluated strains, we focused on exploring the diversity of beauverolides. Since *C. javanica* exhibited both the highest yield and diversity of these compounds, we cultured both strains in five additional liquid media (ZM ½, Q6 ½, GG1, Supermalt, and MMK) and two solid media (BRFT and V+YES). The cultivation and extraction procedures for the solid media were conducted similarly to those used for YM and liquid cultures, with solid cultures treated in the same manner as mycelia from liquid media.

To investigate the presence of secondary metabolites in insect host corpses, a segment was excised from the interface between the insect body and the emerging fruiting body or visible mycelia. The tissue was extracted twice with 1 mL of a 1:1 acetone:methanol solution by vigorous shaking followed by ultrasonication. The combined extracts were filtered, dried under reduced pressure, and resuspended in 50=µL of DMSO for analysis.

All crude extracts were analyzed using ultrahigh-performance liquid chromatography coupled with diode array detection and ion mobility tandem mass spectrometry (UHPLC-DAD-IM-MS/MS) under previously established instrumental settings and conditions.^40^ The raw data were processed using MetaboScape 2022 (Bruker Daltonics, Bremen, Germany) within a retention time range of 1.0–20 min, and the resulting feature table was dereplicated against our in-house spectral library of fungal secondary metabolites. Feature-based molecular networks were constructed on the GNPS platform following our established protocols,^20^ with spectral data subsequently searched against GNPS spectral libraries. The resulting molecular networks were visualized using Cytoscape software.

### Scaled-up Cultivation of *C. javanica* and Purification of Beauverolides

Cultivation in GG1 yielded the highest titers of secondary metabolites when compared to other media. Thus, the scale-up cultivation of *C. javanica* BCC 82944 in this medium was conducted (4 L in total). Both the mycelial and supernatant extracts showed similar secondary metabolite profiles according to our LCMS analyses, and therefore were combined (20 g). An initial liquid-liquid pre-separation was performed using *n*-heptane, methanol, and water. A 9:1 methanol-to-water solution was mixed with an equal volume of *n*-heptane in a separatory funnel, resulting in two immiscible phases and a precipitate in the methanol-water phase. The precipitate was filtered, resulting in Fraction I (6.0 g). The methanol-water phase corresponded then to Fraction II (12.6 g), and the *n*-heptane phase to Fraction III (1.4 g), which were collected, evaporated, and stored for further preparative work.

The purification of Fraction I (6.0 g) was performed using preparative HPLC Büchi Pure C-850 FlashPrep system (Büchi Labortechnik GmbH; Essen, Germany) with the following conditions: Gemini C_18_ column (250 × 50 mm, 10 μm; Phenomenex^®^, Torrance, CA, USA) as a reversed-phase stationary phase. The mobile phase consisted of deionized water with 0.1% formic acid (solvent A) and acetonitrile with 0.1% formic acid (solvent B) at pH 2.5. The flow rate was set at 30 mL/min, and detection was performed using UV at wavelengths of 210, 254, 300, and 350 nm. During each run, a total of 300 mg was injected and 10 runs were performed using a gradient from 30% B to 55 % in 12 minutes, 55% B to 85% in 30 minutes, from 85% B to 100% in 10 minutes and ended with an isocratic elution at 100% for 10 min. A total of six compounds were obtained, **3** (1.2 mg, *t*_R_= 35 min), **5** (5.3 mg, *t*_R_= 42 min), **1** (3.2 mg, *t*_R_= 45 min), **4** (14.1 mg, *t*_R_= 47 min), **6** (6.3 mg, *t*_R_= 56 min) and **2** ( 1.7 mg, *t*_R_= 54 min). The isolation of compound **7** (6.9 mg, *t*_R_= 62 min), in addition to compounds **4** (712.7 mg, *t*_R_= 42 min) and **6** (19.9 mg, *t*_R_= 56 min) resulted from the purification of Fraction II (12.6 g) using the same method described before but with the following gradient: 40 % to 100 % in 60 minutes and ended with isocratic elution at 100% for 10 min.

### Scaled-up Cultivation of *C. blackwelliae* and Purification of Diketopiperazines

To further investigate the secondary metabolites produced by the less virulent fungus *C. blackwelliae* BCC 37653, a scale-up cultivation was conducted on YM medium in Thailand. For the latter reason, the seed culture strategy was different, hence 5 mycelial plugs of fully-grown YMA plates were transferred to 500-mL flasks containing 200 mL of YM medium. A total of fifty flasks (10 L medium) were incubated at 23 °C and 140 rpm. The glucose levels were also daily monitored, and cultures were harvested three days after glucose depletion. The extraction of both mycelia and the supernatant followed the same protocol as for the scale-up cultivation of *C. javanica* to yield 200 and 300 mg of crude extract for the mycelia and supernatant, respectively.

The purification of the crude extracts was done in Germany using a PLC 2250 preparative HPLC system (Gilson, Middleton, WI, USA) equipped with Gemini C_18_ column (250 × 50 mm, 10 μm; Phenomenex^®^, Torrance, CA, USA) with the same conditions as mentioned before despite using the following gradient: from 35% B to 100% in 75 minutes and ended with isocratic elution at 100% B for 10 minutes, which resulted in the purification of five compounds. The purification of the mycelial extract did not result in enough material of the compounds for further analysis. From the supernatant, the following compounds were isolated: **10** (3.2 mg, *t*_R_= 26 min), **9** (2.8 mg, *t*_R_= 27 min), **8** (5.9 mg, *t*_R_= 28 min), **12** (0.48 mg, *t*_R_= 47 min) and **11** (0.57 mg, *t*_R_= 52 min).

*Beauverolide Q* (**1**): White powder; UV-Vis (MeOH): λ_max_ 225; HR-ESI-MS: *m/z* 527.3226 [M+H]^+^ (calcd. 527.3228 for C_29_H_43_N_4_O_5_^+^), 549.3046 [M+Na]^+^ (calcd. 549.6574 for C_29_H_42_N_4_NaO_5_^+^); *t*_R_= 11.64 min (LR-ESI-MS); C_29_H_42_N_4_O_5_ (526.32 g mol^−1^).

*Beauverolide R* (**2**): White powder; UV-Vis (MeOH): λ_max_ 226; NMR data (^1^H: 500 MHz, ^13^C: 125 MHz, DMSO-*d*_6_) see Table **1**; HR-ESI-MS: *m/z* 555.3540 [M+H]^+^ (calcd. 555.3541 for C_31_H_47_N_4_O_5_^+^), 577.3359 [M+Na]^+^ (calcd. 577.7105 for C_31_H_46_N_4_NaO_5_^+^); *t*_R_= 13.19 min (LR-ESI-MS); C_31_H_46_N_4_O_5_ (554.35 g mol^−1^).

*Beauverolide S* (**3**): Colorless oil; UV-Vis (MeOH): λ_max_ 225; HR-ESI-MS: *m/z* 412.2809 [M+H]^+^ (calcd. 412.2806 for C_21_H_38_N_3_O_5_^+^), 434.2625 [M+Na]^+^ (calcd. 434.5254 for C_21_H_37_N_3_NaO_5_^+^); *t*_R_= 9.78 min (LR-ESI-MS); C_21_H_37_N_3_O_5_ (411.27 g mol^−1^).

### Single Crystal Structure via 3D Electron Diffraction of Beauverolide Q (1)

The dry material was suspended in *n*-hexane, and a drop of the suspension was placed onto a holey-carbon-coated copper TEM grid. The grid was air-dried, with the excess solvent soaked up using filter paper. The grids were transferred into a TEM at room temperature and cooled to liquid nitrogen temperature directly in the TEM vacuum. Electron diffraction data were collected using a ThermoFisher GLACIOS TEM operating at 200 kV at liquid nitrogen temperature, employing the EPU-D (ThermoFisher) module. Diffraction patterns were recorded in nanodiffraction mode with the effective probe beam size on the sample of 1 micron.

The data were recorded in MRC format, the MRC stacks were converted to 16-bit TIFFs using the MRC2TIFF converter (10.5281/zenodo.7936068). The TIFF file sequences were then processed in PETS2.^41^ The best-performing dataset was selected for structure analysis. The unit cell parameters were: a = 16.5864 Å, b = 5.1703 Å, c = 17.3397 Å, α = 90.000°, β = 94.011°, γ = 90.000°, with a volume of 1483 Å, corresponding to a monoclinic unit cell. Extinctions corresponding to a *2*[ screw axis along the b-axis were detected, with no further extinctions observed, indicating the *P2*[ space group (No. 4). The expected molecular volume was 719.74 Å³,^42^ corresponding to two molecules in the unit cell. The structure was solved using SHELXD and refined kinematically with SHELXL.^43^ The absolute structure was determined during the dynamical refinement in JANA.^44, 45^ The resulting stereochemistry is shown in Figure 4. More information regarding the crystallographic analysis can be found in supporting information file.

### Antimicrobial and Cytotoxic Assays

Serial dilution assays were performed in 96-well microtiter plates to determine the minimum inhibitory concentrations (MICs) against a panel of yeasts, filamentous fungi, and bacteria, as well as the half-maximal inhibitory concentrations (IC_50_) against human cell lines, following our previously described protocol.^19^

## Data Availability

All data supporting the findings of this study are included in the article and its supplementary materials.

## Acknowledgments

The authors wish to thank Dr. Kirsten Harmrolfs, Esther Surges and Aileen Golasch for recording the NMR spectra as well as HR-MS data, Ulrike Beutling for the metabolomics analyses, Wera Collisi for conducting the bioassays, Silke Reinecke for its advisory assistance, Wasana Noisripoom for the sequencing data of the studied strains, Nopporn Chutiwitoonchai for conducting the antiviral screening, and Natalia Andrea Llanos-López for carrying out the nematode assay. This work benefited from the European Union’s Horizon 2020 Research and Innovation Staff Exchange program (MSCA-RISE grant no. 101008129, acronym: MYCOBIOMICS), the Southeast Asia–Europe Joint Funding Scheme (SEA– Europe Grant number JFS20ST-127, Acronym: Antiviralfun), and the Grant no. P21-50844, from the National Science and Technology Development Agency (NSTDA). Esteban Charria-Girón was supported by the HZI POF IV Cooperativity and Creativity Project Call and the DAAD stipend 57694196 Forschungsstipendien für Doktorandinnen und Doktoranden (2024). Rita Toshe received a grant from the Bischöfliche Stiftung Cusanuswerk. The Alexander von Humboldt (AvH) Foundation is gratefully acknowledged for granting S.S.E. the Georg-Forster Fellowship for Experienced Researchers Stipend (Ref 3.4-1222288-EGY-GF-E).

## Notes

### Competing Interest Statement

The authors have declared no competing interest.

