## Supporting Information for "Chemical clues to infection: Metabolome differentiation underlies host colonization of potential biocontrol agents from the entomopathogenic genus *Cordyceps*"

#### Contents of Supporting Information


| # | Contents | Page |
|---|---|---|
| **1** | Figure S1. LR-ESI-MS spectrum of **1**. | **S4** |
| **2** | Figure S2. HR-ESI-MS spectrum of **1**. | **S5** |
| **3** | Table S1. Crystallographic data for **GRGDS TFA** and **Beauveriolide Q** | **S6** |
| **4** | Table S2. Dynamical refinement of **GRGDS TFA** and **Beauveriolide Q** | **S7** |
| **5** | Figure S3. Absolute structure determination of **GRGDS TFA** – calibration of the data workflow. Left: the model as obtained in the structure determination procedure, right: the inverted model, bottom: the expected stereochemistry of **GRGDS**. | **S8** |
| **6** | Figure S4. Overlay of the molecular conformations of compound **1** (stick representation) and beauveriolide I (**4**) (CSD code 2332378, wire representation) as determined from electron diffraction data. | **S8** |
| **7** | Figure S5. Molecular family encompassing beauveriolides produced by *Cordyceps javanica* BCC 82944. Hexagon-shaped nodes represent isolated congeners, while figure-eight–shaped nodes indicate dereplicated beauveriolides with putative annotations based on previously reported structures. Suggested structural additions or losses between nodes are highlighted for the isolated metabolites. | **S9** |
| **8** | Figure S6. LR-ESI-MS spectrum of **2**. | **S10** |
| **9** | Figure S7. HR-ESI-MS spectrum of **2**. | **S11** |
| **10** | Figure S8. $^1$H NMR spectrum of **2** in DMSO-$d_6$ at 500 MHz. | **S12** |
| **11** | Figure S9 $^{13}$C NMR spectrum of **2** in DMSO-$d_6$ at 125 MHz. | **S13** |
| **12** | Figure S10. $^1$H–$^1$H COSY spectrum of **2** in DMSO-$d_6$ at 500 MHz. | **S14** |
| **13** | Figure S11. HMBC spectrum of **2** in DMSO-$d_6$ at 500 MHz. | **S15** |
| **14** | Figure S12. HSQC spectrum of **2** in DMSO-$d_6$ at 500 MHz. | **S16** |
| **15** | Figure S13. ROESY spectrum of **2** in DMSO-$d_6$ at 500 MHz. | **S17** |
| **16** | Figure S14. LR-ESI-MS spectrum of **3**. | **S18** |
| **17** | Figure S15. HR-ESI-MS spectrum of **3**. | **S19** |
| **18** | Figure S16. LR-ESI-MS spectrum of **4**. | **S20** |
| **19** | Figure S17. HR-ESI-MS spectrum of **4**. | **S21** |
| **20** | Figure S18. $^1$H NMR spectrum of **4** in DMSO-$d_6$ at 500 MHz. | **S22** |
| **21** | Figure S19. $^1$H–$^1$H COSY spectrum of **4** in DMSO-$d_6$ at 500 MHz. | **S23** |
| **22** | Figure S20. LR-ESI-MS spectrum of **5**. | **S24** |
| **23** | Figure S21. $^1$H NMR spectrum of **5** in DMSO-$d_6$ at 500 MHz. | **S25** |
| **24** | Figure S22. $^1$H–$^1$H COSY spectrum of **5** in DMSO-$d_6$ at 500 MHz. | **S26** |
| **25** | Figure S23. LR-ESI-MS spectrum of **6**. | **S27** |
| **26** | Figure S24. HR-ESI-MS spectrum of **6**. | **S28** |
| **27** | Figure S25. $^1$H NMR spectrum of **6** in DMSO-$d_6$ at 500 MHz. | **S29** |
| **28** | Figure S26. LR-ESI-MS spectrum of **7**. | **S30** |
| **29** | Figure S27. HR-ESI-MS spectrum of **7**. | **S31** |
| **30** | Figure S28. $^1$H NMR spectrum of **7** in DMSO-$d_6$ at 500 MHz. | **S32** |
| **31** | Figure S29. $^1$H–$^1$H COSY spectrum of **7** in DMSO-$d_6$ at 500 MHz. | **S33** |
| **32** | Figure S30. HMBC spectrum of **7** in DMSO-$d_6$ at 500 MHz. | **S34** |
| **33** | Figure S31. HSQC spectrum of **7** in DMSO-$d_6$ at 500 MHz. | **S35** |
| **34** | Figure S32. LR-ESI-MS spectrum of **8**. | **S36** |
| **35** | Figure S33. HR-ESI-MS spectrum of **8**. | **S37** |
| **36** | Figure S34. $^1$H NMR spectrum of **8** in DMSO-$d_6$ at 500 MHz. | **S38** |
| **37** | Figure S35. LR-ESI-MS spectrum of **9**. | **S39** |
| **38** | Figure S36. HR-ESI-MS spectrum of **9**. | **S40** |
| **39** | Figure S37. $^1$H NMR spectrum of **9** in DMSO-$d_6$ at 500 MHz. | **S41** |
| **40** | Figure S38. $^{13}$C NMR spectrum of **9** in DMSO-$d_6$ at 125 MHz. | **S42** |
| **41** | Figure S39. $^1$H–$^1$H COSY spectrum of **9** in DMSO-$d_6$ at 500 MHz. | **S43** |
| **42** | Figure S40. HMBC spectrum of **9** in DMSO-$d_6$ at 500 MHz. | **S44** |
| **43** | Figure S41. HSQC spectrum of **9** in DMSO-$d_6$ at 500 MHz. | **S45** |

| | | |
|---|---|---|
| **44** | Figure S42. ROESY spectrum of **9** in DMSO-$d_6$ at 500 MHz. | **S46** |
| **45** | Figure S43. LR-ESI-MS spectrum of **10**. | **S47** |
| **46** | Figure S44. HR-ESI-MS spectrum of **10**. | **S48** |
| **47** | Figure S45. $^1$H NMR spectrum of **10** in DMSO-$d_6$ at 500 MHz. | **S49** |
| **48** | Figure S46. $^1$H–$^1$H COSY spectrum of **10** in DMSO-$d_6$ at 500 MHz. | **S50** |
| **49** | Figure S47. HMBC spectrum of **10** in DMSO-$d_6$ at 500 MHz. | **S51** |
| **50** | Figure S48. ROESY spectrum of **10** in DMSO-$d_6$ at 500 MHz. | **S52** |
| **51** | Figure S49. LR-ESI-MS spectrum of **11**. | **S53** |
| **52** | Figure S50. $^1$H NMR spectrum of **11** in DMSO-$d_6$ at 500 MHz. | **S54** |
| **53** | Figure S51. DEPTQ spectrum of **11** in DMSO-$d_6$ at 125 MHz. | **S55** |
| **54** | Figure S52. $^1$H–$^1$H COSY spectrum of **11** in DMSO-$d_6$ at 500 MHz. | **S56** |
| **55** | Figure S53. HMBC spectrum of **11** in DMSO-$d_6$ at 500 MHz. | **S57** |
| **56** | Figure S54. HSQC spectrum of **11** in DMSO-$d_6$ at 500 MHz. | **S58** |
| **57** | Figure S55. ROESY spectrum of **11** in DMSO-$d_6$ at 500 MHz. | **S59** |
| **58** | Figure S56. HR-ESI-MS spectrum of **12**. | **S60** |
| **59** | Figure S57. $^1$H NMR spectrum of **12** in DMSO-$d_6$ at 500 MHz. | **S61** |
| **60** | Figure S58. $^1$H–$^1$H COSY spectrum of **12** in DMSO-$d_6$ at 500 MHz. | **S62** |
| **61** | Figure S59. HMBC spectrum of **12** in DMSO-$d_6$ at 500 MHz. | **S63** |
| **61** | Figure S60. HSQC spectrum of **12** in DMSO-$d_6$ at 500 MHz. | **S64** |
| **62** | Table S3. Antimicrobial properties of the isolated metabolites. | **S65** |
| **63** | Table S4. Cytotoxic properties of the isolated metabolites. | **S65** |

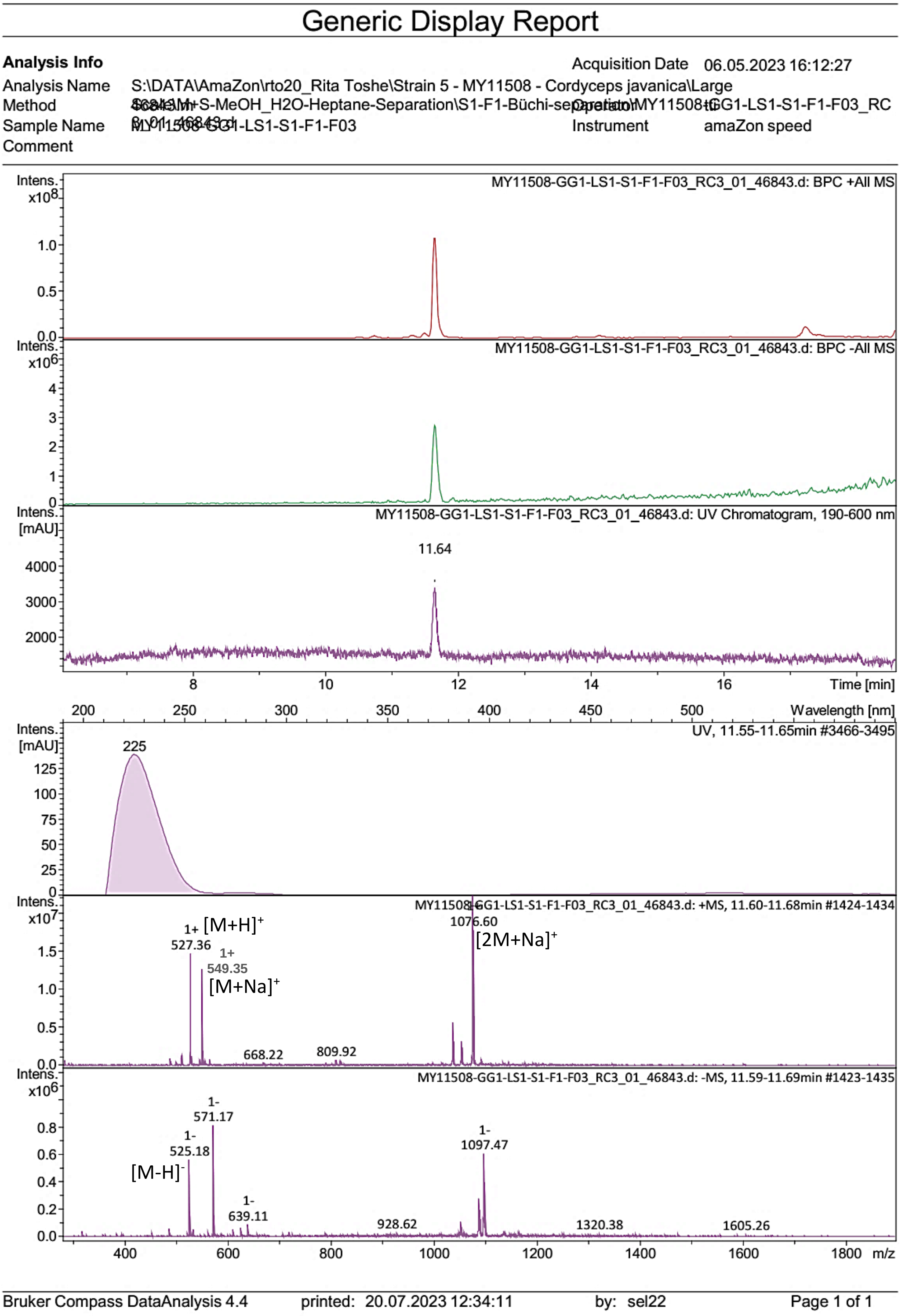


Figure S1. LR-ESI-MS spectrum of **1**.

#### Generic Display Report

| **Analysis Info** | | Acquisition Date | 10.05.2023 20:44:20 |
|---|---|---|---|
| Analysis Name | S:\DATA\MaXis\rto20_Rita Toshe\23_05\MY11508-GG1-LS1-S-F1-F03_23_01_11516.d | | |
| Method | pos_säure_10000_screening_ms_100_2500_line.m | Operator | ate06 |
| Sample Name | MY11508-GG1-LS1-S-F1-F03 | Instrument | maXis |
| Comment | Screening01<br>Waters Acquity UPLC BEH C_18 1,7um 2.1x50mm | | |

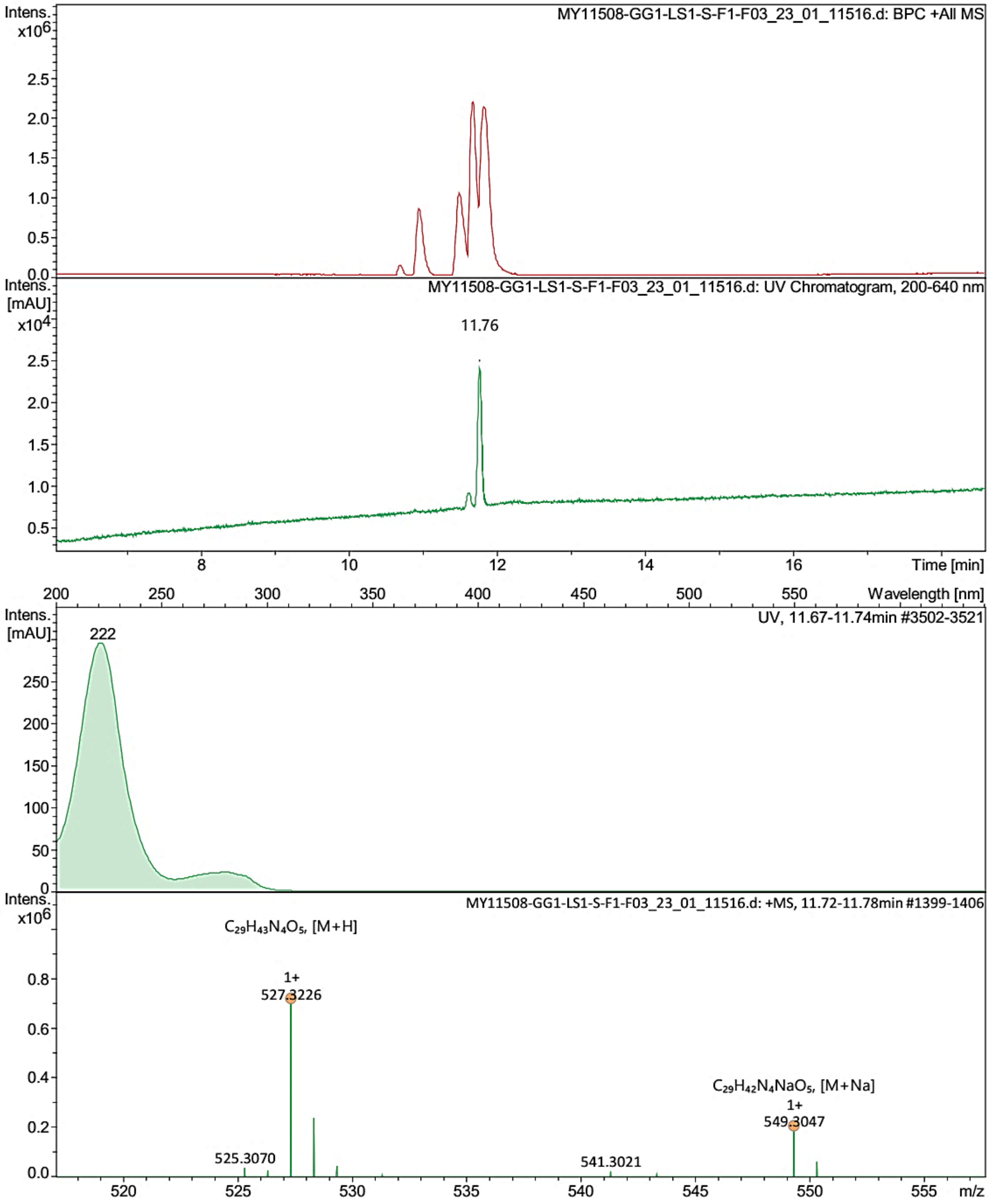




Figure S2. HR-ESI-MS spectrum of **1**.

**Structure determination of GRGDS, calibration of the data workflow for absolute structure determinaiton of Beauveriolide Q**

**GRGDS** is a five-amino-acid peptide used for the calibration of the absolute structure determination procedure with electron diffraction. The compound was purchased from GenScript Biotech (Leiden, Netherlands). Needle-like crystals, less than 0.5 µm wide and approximately 10 µm long, were grown from methanol. 3D ED data of **GRGDS** was collected using EPU-D and processed with PETS2, exactly as described for **Beauveriolide Q** in the Methods section of the main text.

The best-performing dataset was selected for structure analysis. The structure was solved with SHELXD and refined kinematically with SHELXL in the Olex2 environment. The crystallographic details are provided in Table S1.

Table S1. Crystallographic data for **GRGDS TFA** and **Beauveriolide Q**

| | **GRGDS TFA** | **Beauveriolide Q** |
|---|---|---|
| Crystal data | | |
| Chemical formula | $C_{17}H_{28}N_8O_8$ $F_3O_2C_2H_1$ | $C_{29}H_{42}N_4O_5$ |
| Mr | 586.47 | 526.67 |
| Crystal system, space group | Monoclinic, *C2* | Monoclinic, $P2_1$ |
| Temperature (K) | 77 | 77 |
| a, b, c (Å) | 29.231, 4.546, 19.640 | 16.586, 5.170, 17.340 |
| β (°) | 106.70 | 94.91 |
| V (Å$^3$) | 2499.8 | 1481.5 |
| Z, Z’ | 4, 1 | 2, 1 |
| Radiation type | electrons, λ = 0.0251 Å | electrons, λ = 0.0251 Å |
| Crystal size (mm) | 0.005 x 0.001 x 0.0002 | 0.008 x 0.001 x 0.0002 |
| Data collection | | |
| Diffractometer | Thermo Fisher GLACIOS, EPU-D | Thermo Fisher GLACIOS, EPU-D |
| No. of measured, independent and observed [I > 2σ(I)] reflections | 4085, 2632, 2101 | 5306, 3513, 3221 |
| Rint | 0.131 | 0.137 |
| θmax (°) | 0.8 | 0.8 |
| (sin θ/λ)max (Å$^{-1}$) | 0.550 | 0.569 |
| Data resolution, Å | 0.91 | 0.91 |
| Data completeness | 0.722 | 0.761 |
| Refinement | | |

| | | |
|---|---|---|
| $R[F^2 > 2\sigma(F^2)]$, wR(F2), S | 0.173, 0.429, 1.64 | 0.256, 0.637, 3.31 |
| No. of reflections | 2632 | 3513 |
| No. of parameters | 178 | 146 |
| No. of restraints | 1 | 1 |
| H-atom treatment | H atoms treated by a mixture of independent and constrained refinement | H-atom parameters constrained |
| Absolute structure: dynamical refinement in JANA, $R_{obs}$ | 15.62 / 12.50 | 20.79 / 22.09 |
| CSD deposition code | 2391399 | 2391744 |
| Original diffraction data | doi.org/10.5281/zenodo.13938423 | doi.org/10.5281/zenodo.13943915 |

Computer programs: *SHELXD* (Sheldrick, 2008), *SHELXL* 2018/3 (Sheldrick, 2015), Olex2 1.5 (Dolomanov *et al.*, 2009).

To our surprise, **GRGDS** co-crystallized with trifluoroacetic acid (**TFA**), which was evidently used in the peptide synthesis.

We then performed the absolute structure determination of the **GRGDS TFA** co-crystal through dynamical refinement from electron diffraction data using JANA. We did not carry out a full dynamical refinement of all structure parameters, focusing solely on the absolute structure determination.

The refinement statistics of the initial structure model obtained during structure determination showed worse figures of merit than the inverted enantiomer structure (Figure S3, Table S2).

Despite the higher R-factors obtained in the dynamical refinement (Table S2), the initial stereochemistry of the **GRGDS** molecule is correct. This suggests that somewhere in the data processing pipeline, a flipping of the diffraction patterns occurs. Instead of searching for the point where the data flipping occurs and correcting it, we now consider the entire procedure calibrated: the correct enantiomer should yield higher R-factors in the dynamical refinement.

Table S2. Dynamical refinement of **GRGDS TFA** and **Beauveriolide Q**

| | $R_{obs}$ | $R_{all}$ | $wR2_{all}$ |
|---|---|---|---|
| **GRGDS TFA** | 15.62 | 17.10 | 31.89 |
| **GRGDS TFA** inverted model | 12.50 | 13.98 | 24.50 |
| **Beauveriolide Q** | 20.79 | 22.19 | 40.44 |
| **Beauveriolide Q** inverted model | 22.09 | 23.46 | 42.80 |

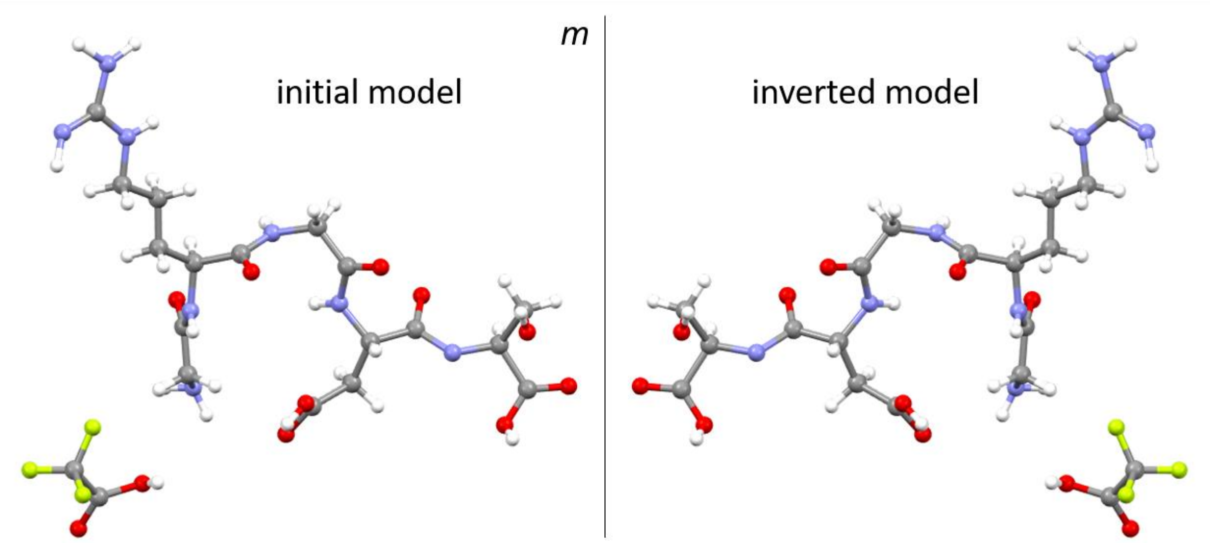


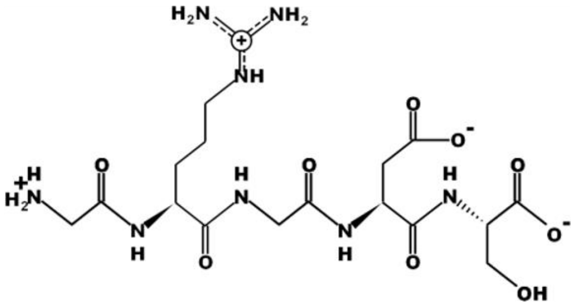


Figure S3. Absolute structure determination of **GRGDS TFA** – calibration of the data workflow. Left: the model as obtained in the structure determination procedure, right: the inverted model, bottom: the expected stereochemistry of **GRGDS**.

The same dynamical refinement procedure was then applied to **Beauveriolide Q**. The inverted model showed higher R-factors, indicating that it has the correct stereochemistry. The correct stereochemistry of MY and molecular packing within the crystal structure is illustrated in Figure S2.

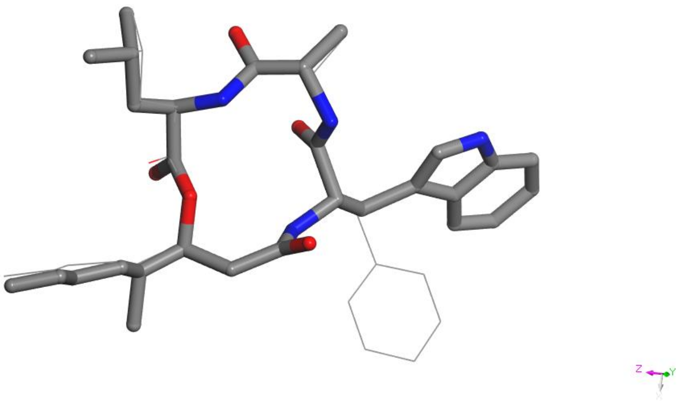

Figure S4. Overlay of the molecular conformations of compound **1** (stick representation) and beauveriolide I (**4**) (CSD code 2332378, wire representation) as determined from electron diffraction data.

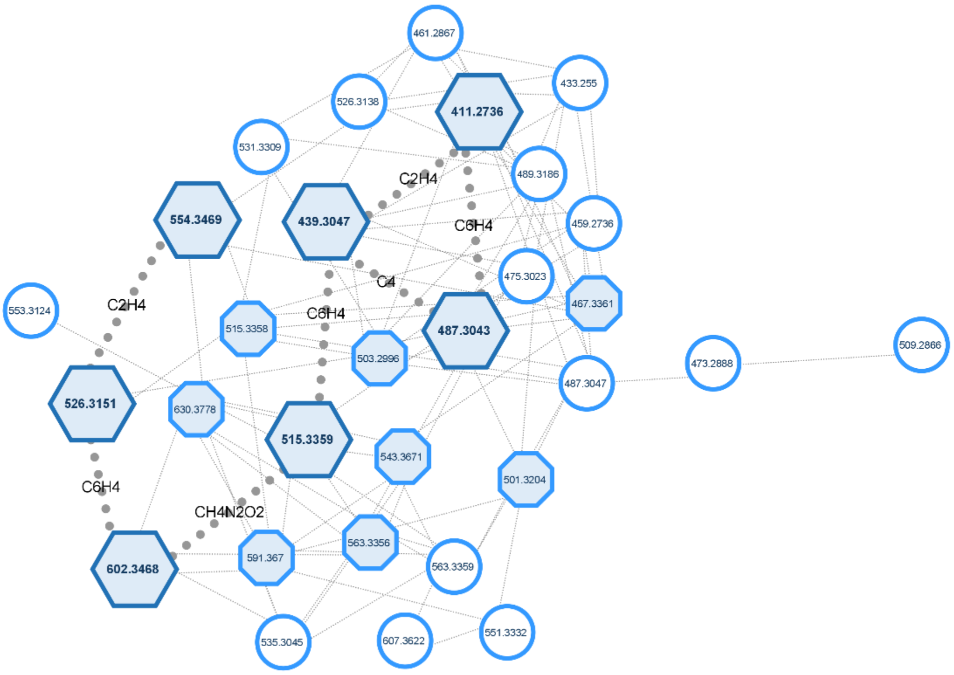


Figure S5. Molecular family encompassing beauveriolides produced by *Cordyceps javanica* BCC 82944. Hexagon-shaped nodes represent isolated congeners, while figure-eight–shaped nodes indicate dereplicated beauveriolides with putative annotations based on previously reported structures. Suggested structural additions or losses between nodes are highlighted for the isolated metabolites.

### Generic Display Report

| **Analysis Info** | | Acquisition Date | 06.05.2023 19:49:45 |
|---|---|---|---|
| Analysis Name | S:\DATA\AmaZon\rto20_Rita Toshe\Strain 5 - MY11508 - Cordyceps javanica\Large | | |
| Method | S6849Mh+S-MeOH_H2O-Heptane-Separation\S1-F1-Büchi-separation\MY11508-GG1-LS1-S1-F1-F09_RD | Operator | tu |
| Sample Name | MY11508-GG1-LS1-S1-F1-F09 1_01_46849.d | Instrument | amaZon speed |
| Comment | | | |

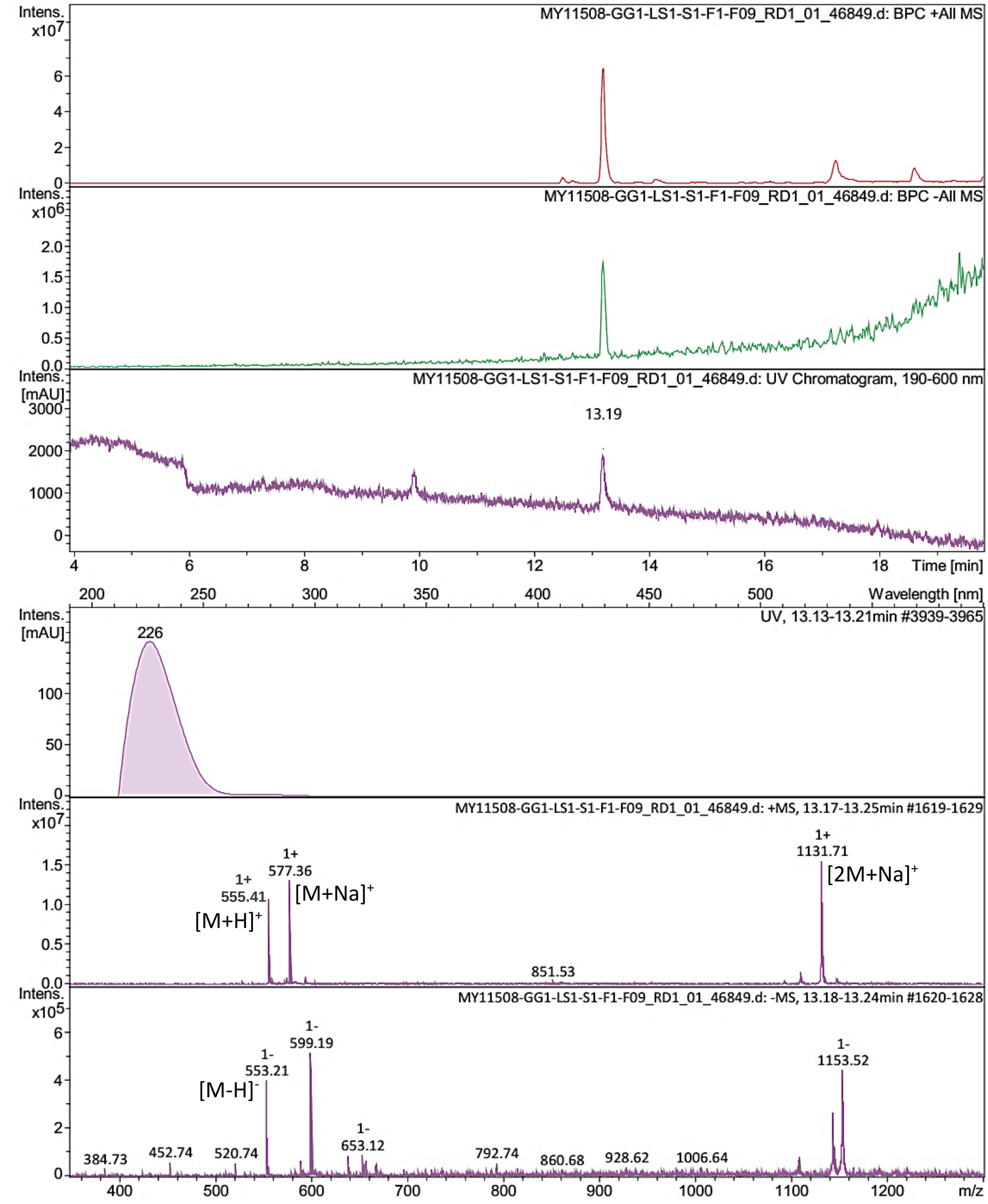


Bruker Compass DataAnalysis 4.4 printed: 12.07.2023 10:59:53 by: sel22 Page 1 of 1

Figure S6. LR-ESI-MS spectrum of **2**.

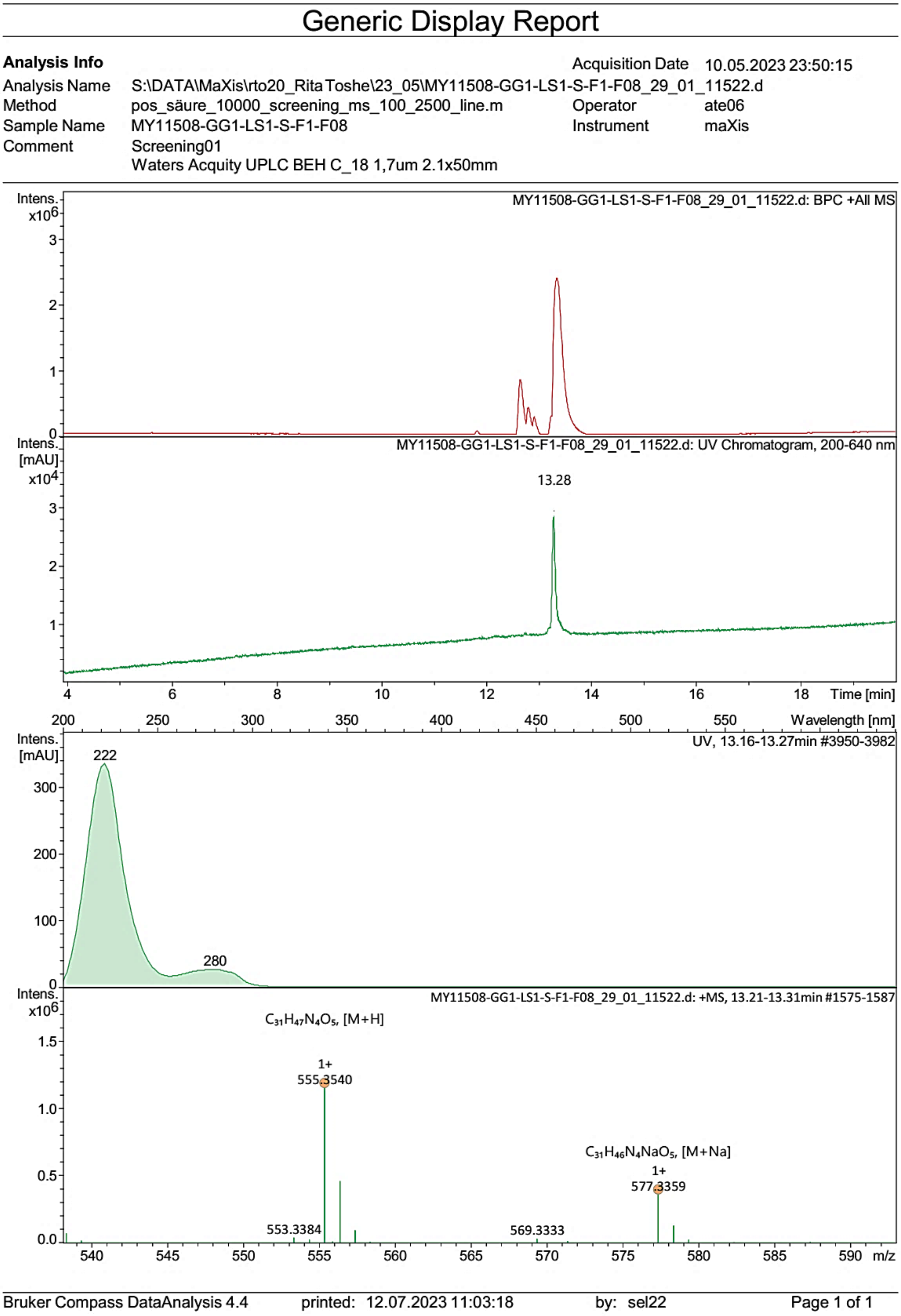


Figure S7. HR-ESI-MS spectrum of **2**.

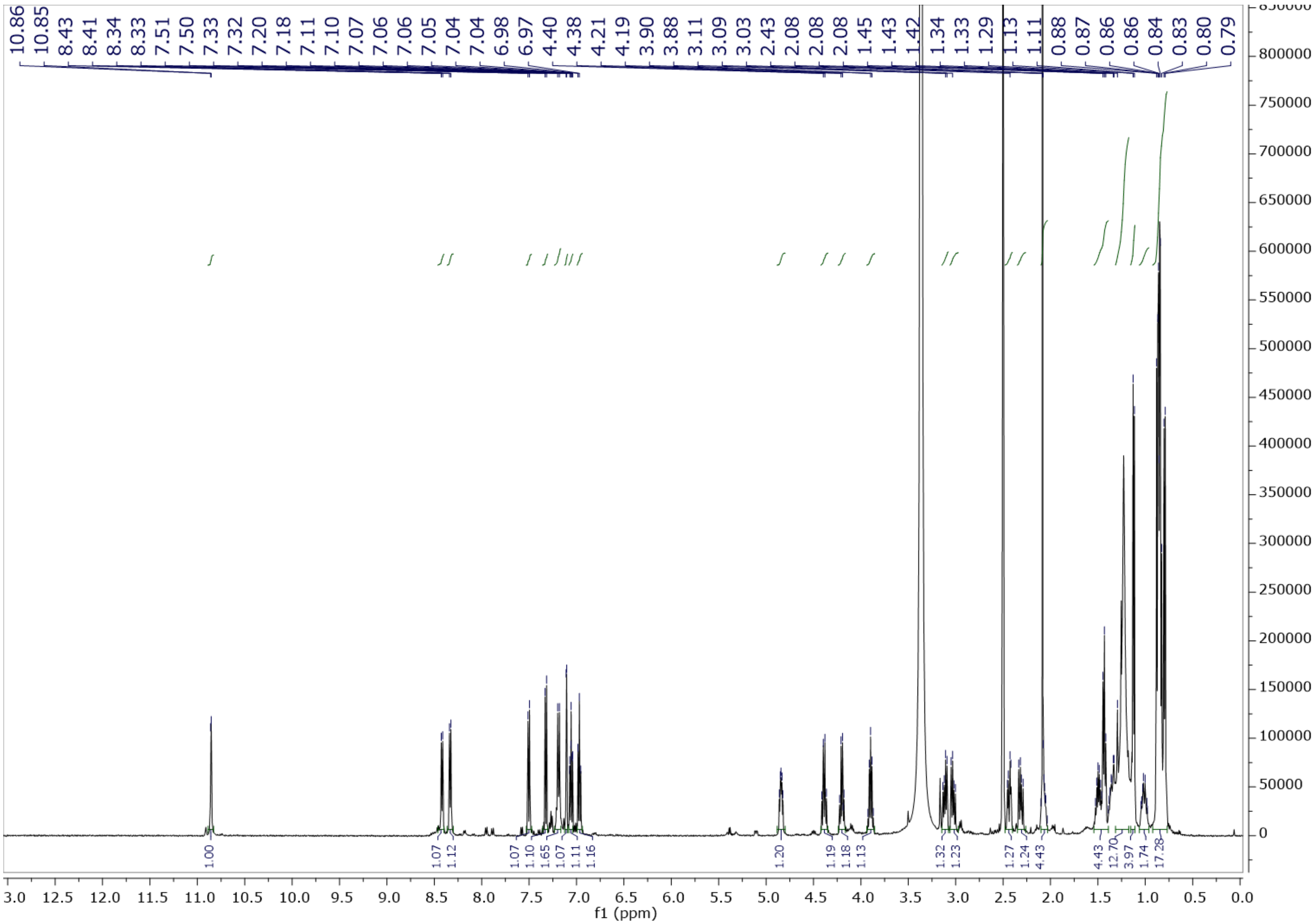

Figure S8. $^{1}H$ NMR spectrum of **2** in DMSO-$d_6$ at 500 MHz.

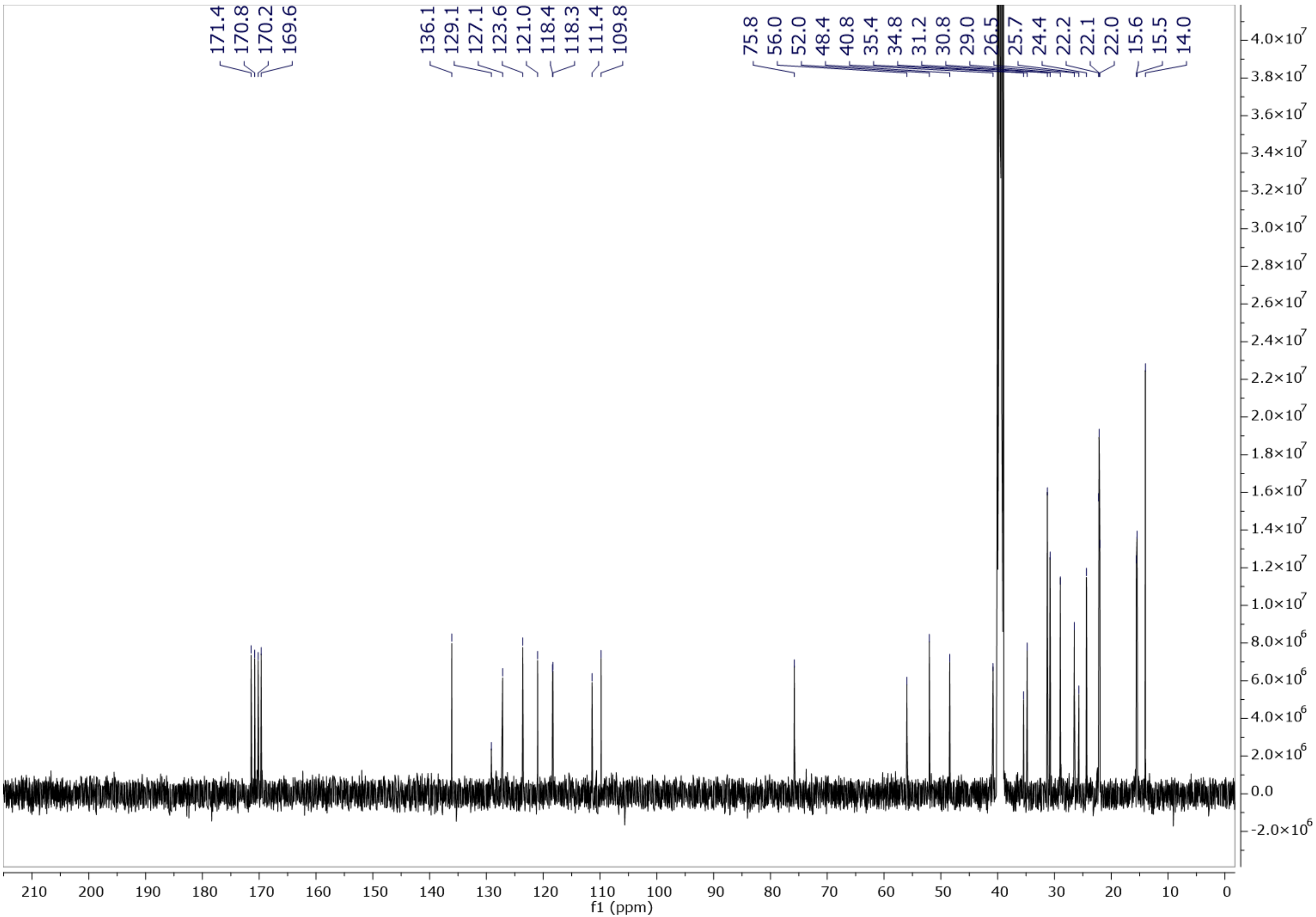


Figure S9. $^{13}$C NMR spectrum of **2** in DMSO-$d_6$ at 125 MHz.

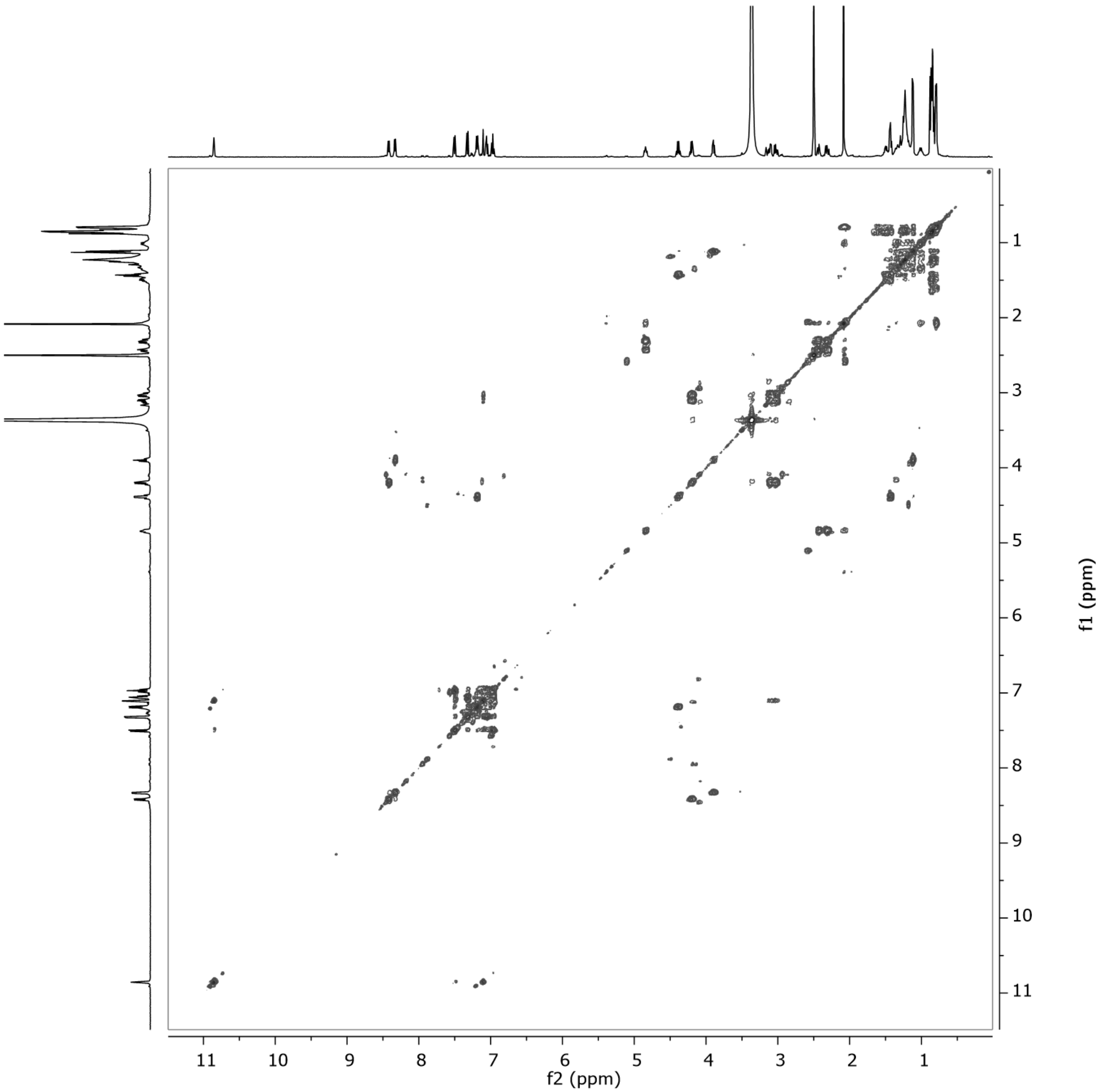


Figure S10. $^1H$–$^1H$ COSY spectrum of **2** in DMSO-$d_6$ at 500 MHz.

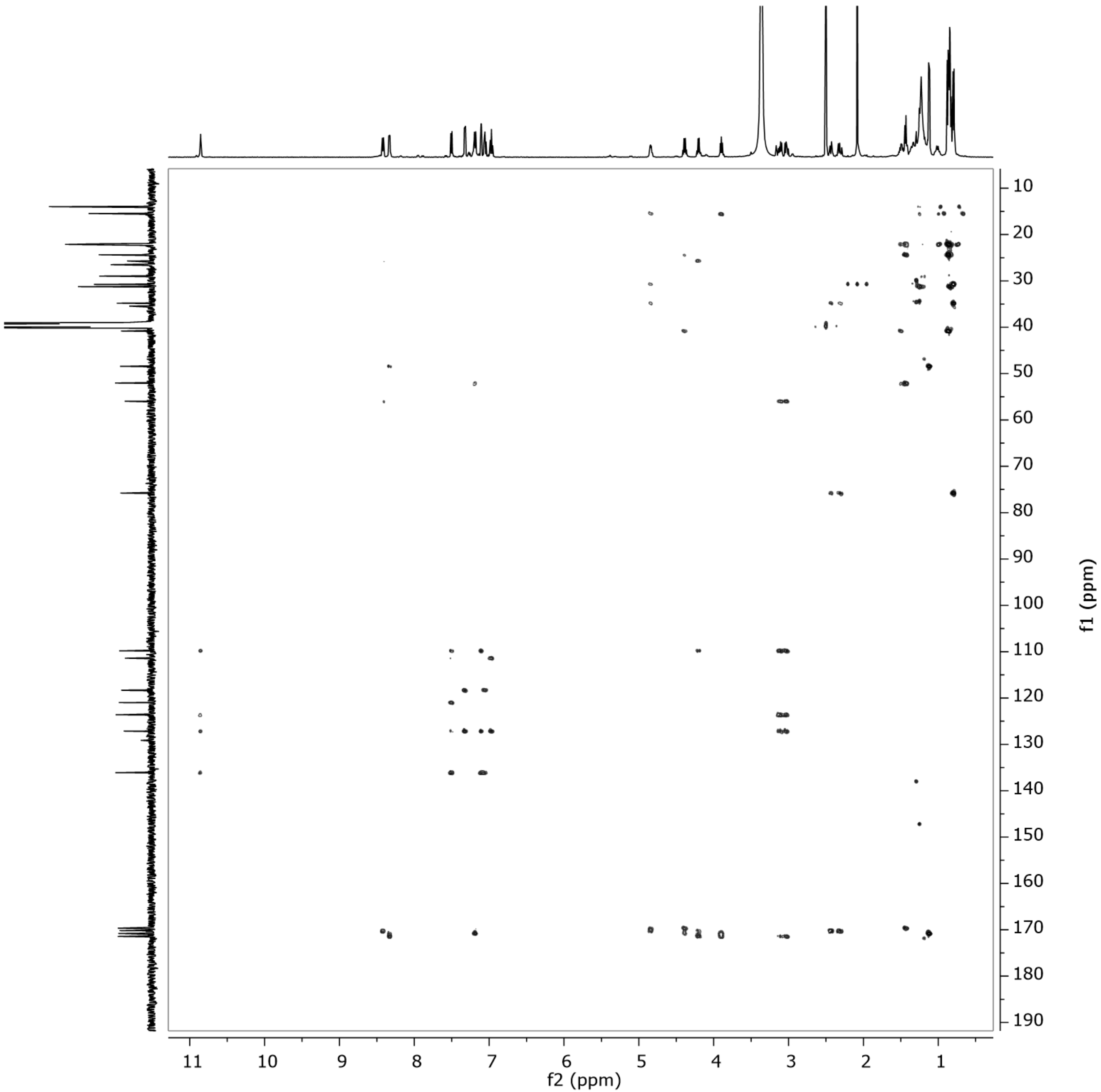


Figure S11. HMBC spectrum of **2** in DMSO-$d_6$ at 500 MHz.

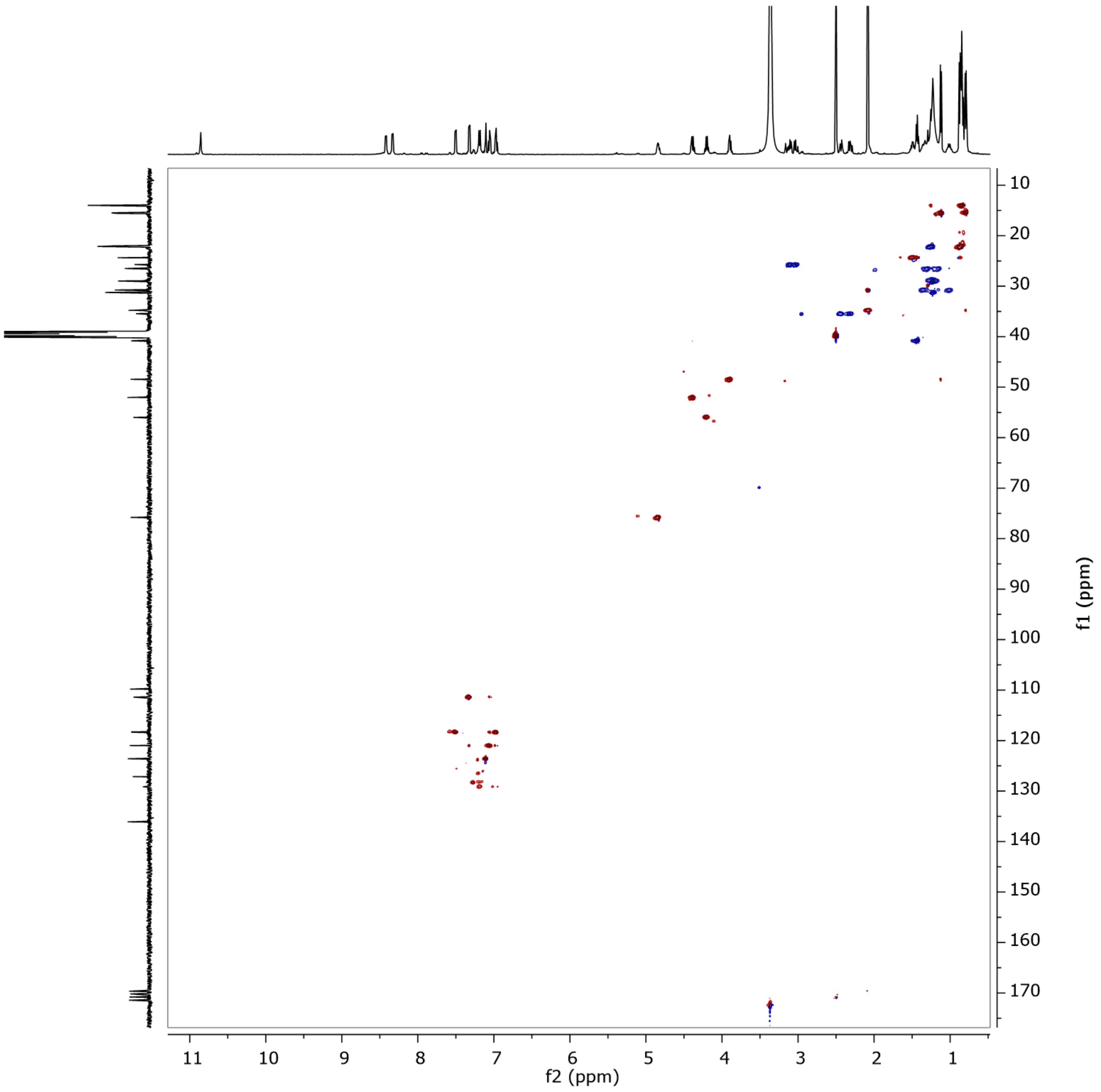


Figure S12. HSQC spectrum of **2** in DMSO-$d_6$ at 500 MHz.

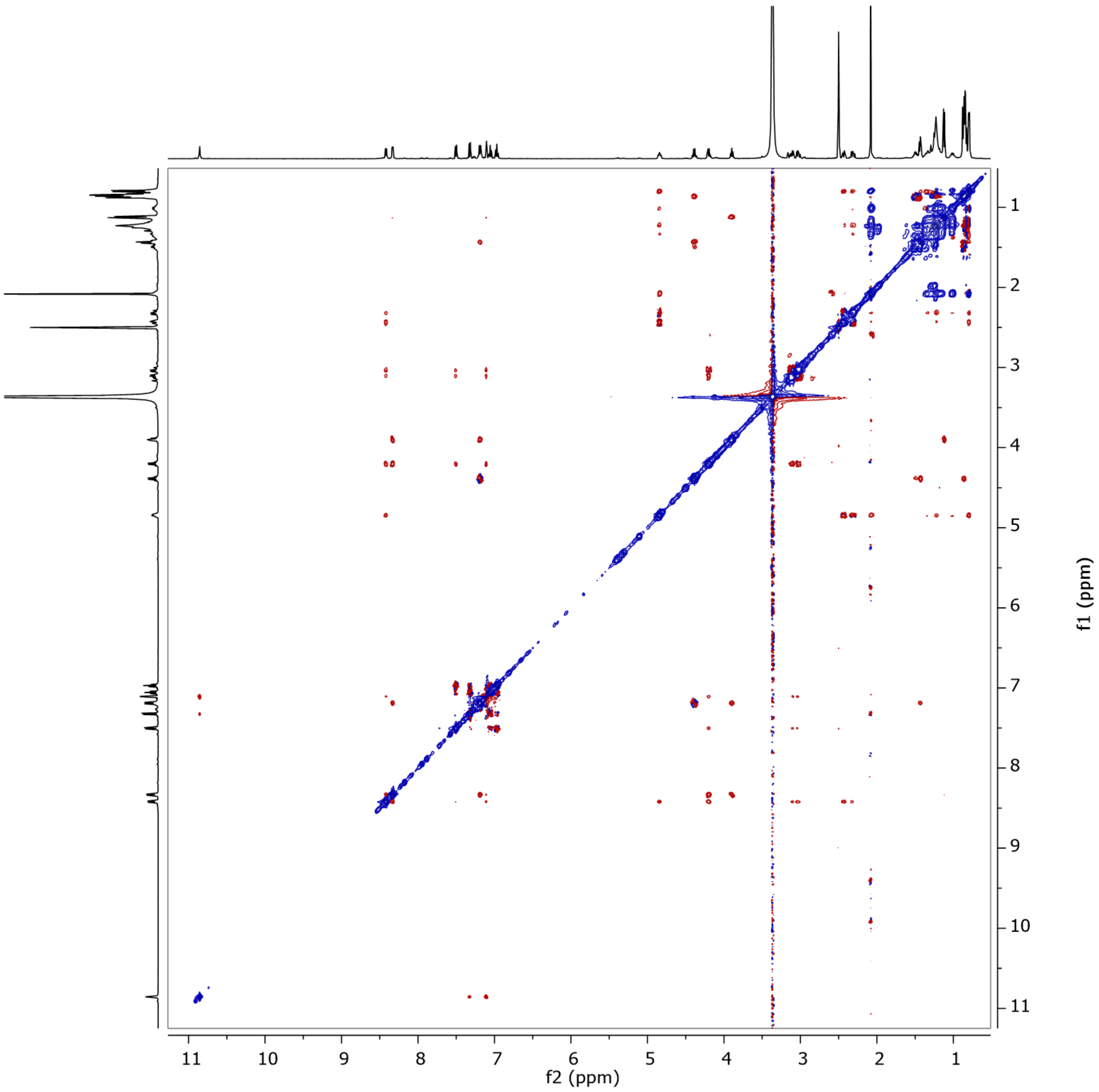


Figure S13. ROESY spectrum of **2** in DMSO-$d_6$ at 500 MHz.

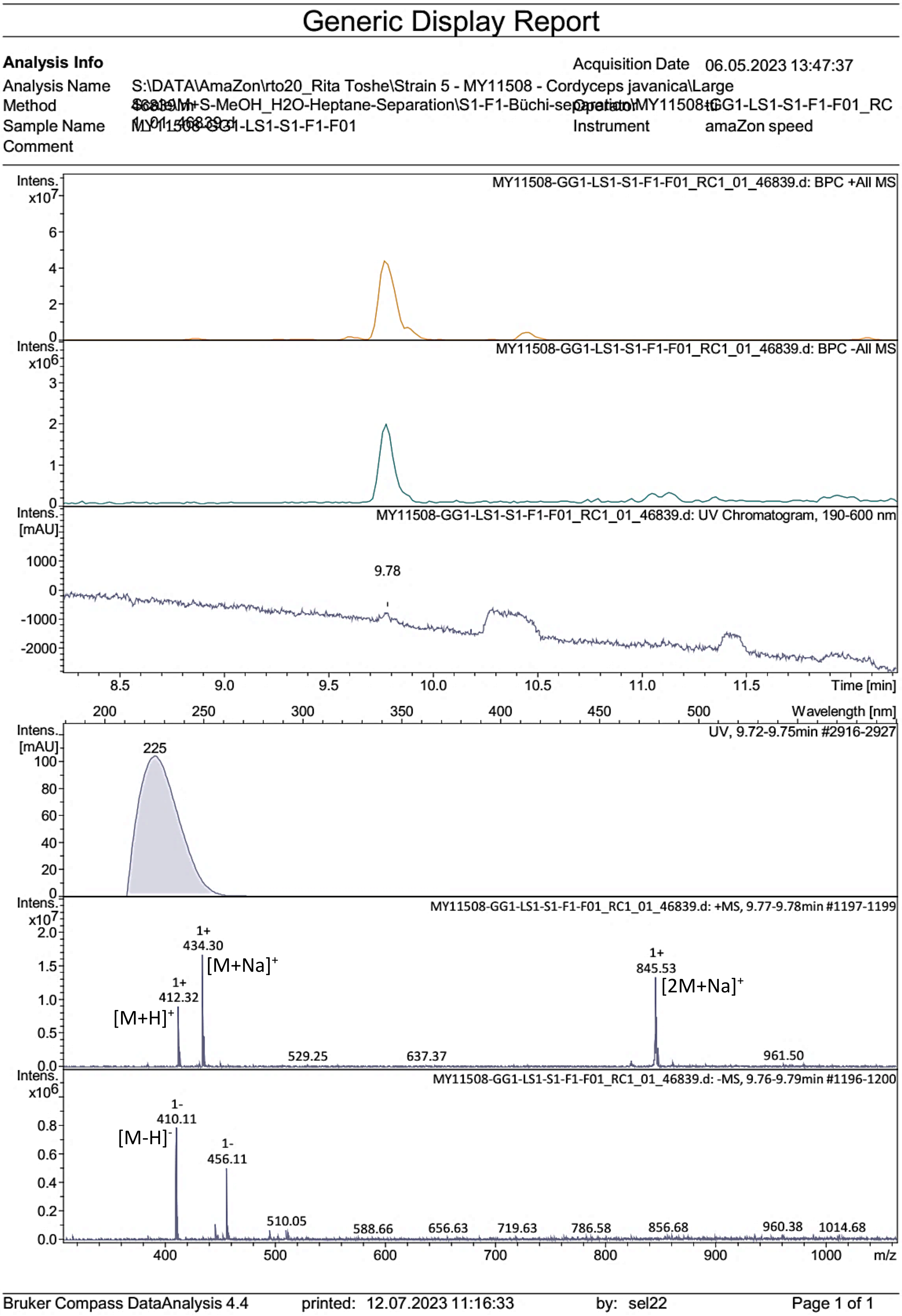


Figure S14. LR-ESI-MS spectrum of **3**.

### Generic Display Report

**Analysis Info**

| | |
|---|---|
| Acquisition Date | 10.05.2023 19:42:25 |
| Analysis Name | S:\DATA\MaXis\rto20_Rita Toshe\23_05\MY11508-GG1-LS1-S-F1-F01_21_01_11514.d |
| Method | pos_säure_10000_screening_ms_100_2500_line.m |
| Operator | ate06 |
| Sample Name | MY11508-GG1-LS1-S-F1-F01 |
| Instrument | maXis |
| Comment | Screening01<br>Waters Acquity UPLC BEH C_18 1,7um 2.1x50mm |

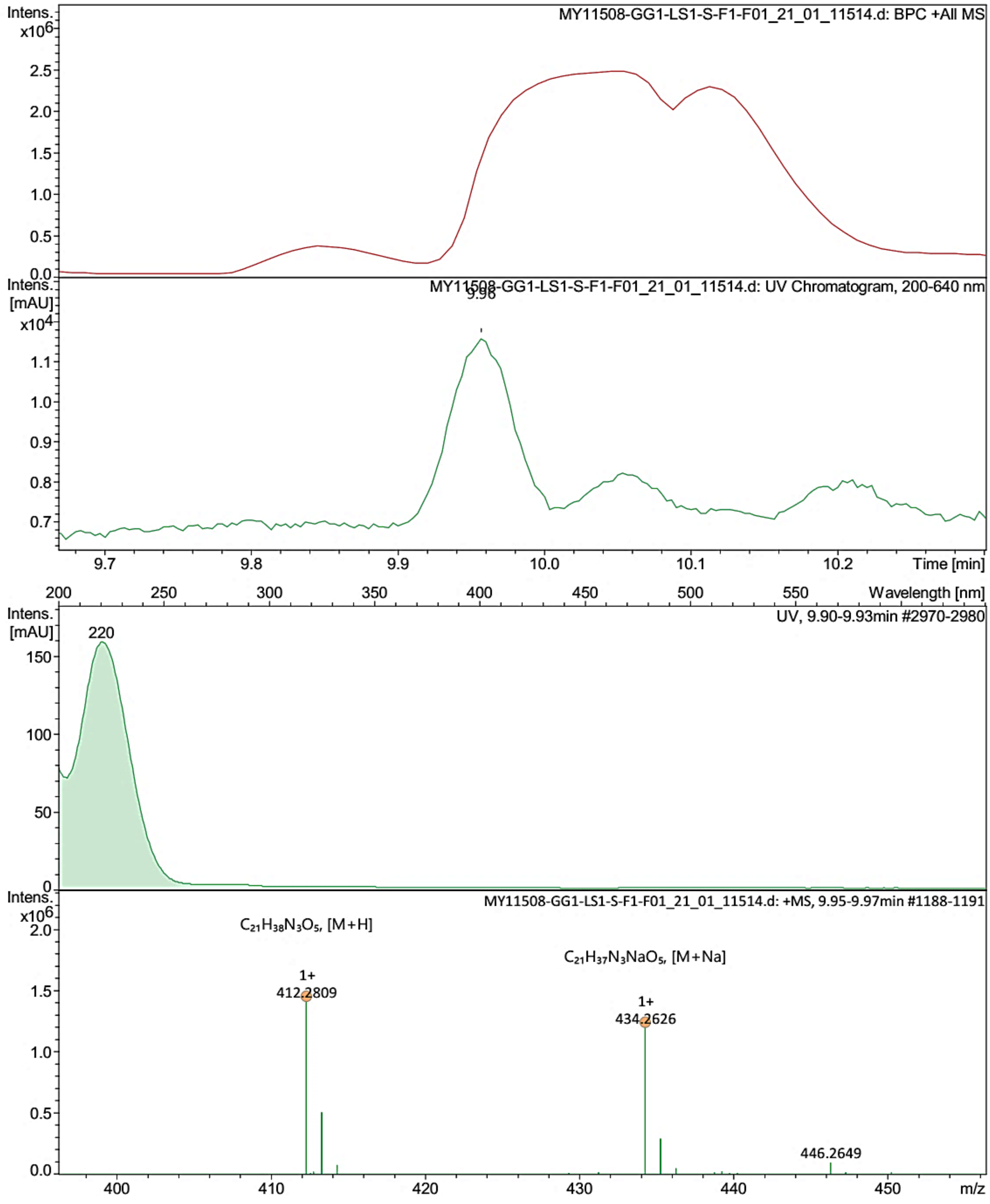


| | | | |
|---|---|---|---|
| Bruker Compass DataAnalysis 4.4 | printed: 12.07.2023 11:19:13 | by: sel22 | Page 1 of 1 |

Figure S15. HR-ESI-MS spectrum of **3**.

### Generic Display Report

**Analysis Info**

| | | | |
|---|---|---|---|
| Analysis Name | S:\DATA\AmaZon\rto20_Rita Toshe\23-05\MY11508-GG1-LS1-S1-F1-F04_RC4_01_46844.d | Acquisition Date | 06.05.2023 16:48:41 |
| Method | 46844.m | Operator | tti |
| Sample Name | MY11508-GG1-LS1-S1-F1-F04 | Instrument | amaZon speed |
| Comment | | | |

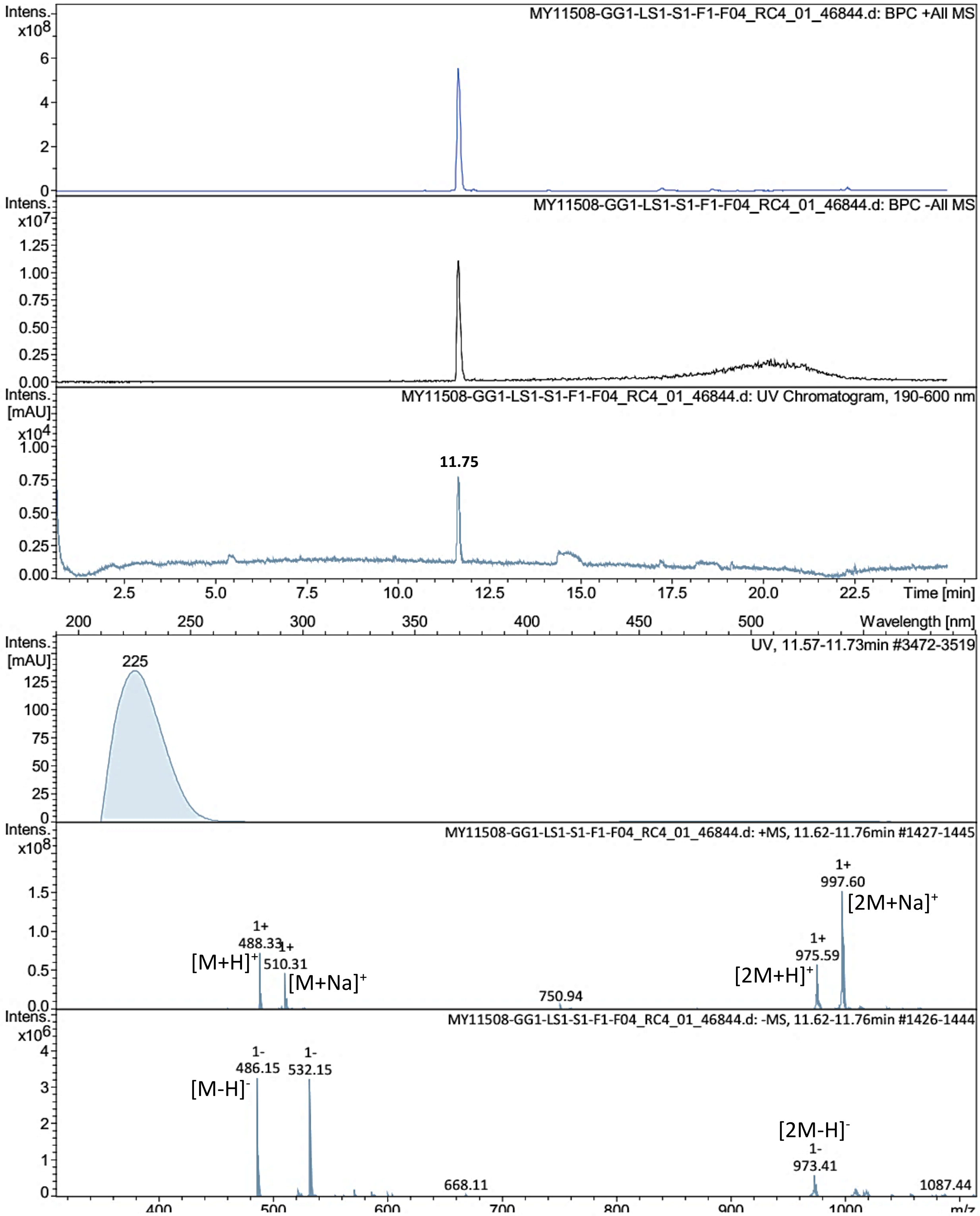


Bruker Compass DataAnalysis 4.4 printed: 09.05.2023 16:29:02 by: sel22 Page 1 of 1

Figure S16. LR-ESI-MS spectrum of **4**.

### Display Report

**Analysis Info**

| | | | |
|---|---|---|---|
| Analysis Name | S:\DATA\MaXis\rto20_Rita Toshe\Strain 5 - MY11508 - Cordyceps javanica\MY11508-GG1-LS1-S1-F4_59_01_11334.d | Acquisition Date | 10.03.2023 09:09:11 |
| Method | pos_säure_10000_screening_ms_100_2500_line.m | Operator | ate06 |
| Sample Name | MY11508-GG1-LS1-S1-F4 | Instrument | maXis 255552.00037 |
| Comment | Screening01<br>Waters Acquity UPLC BEH C_18 1,7um 2.1x50mm | | |

**Acquisition Parameter**

| | | | | | |
|---|---|---|---|---|---|
| Source Type | ESI | Ion Polarity | Positive | Set Nebulizer | 4.0 Bar |
| Focus | Not active | Set Capillary | 4500 V | Set Dry Heater | 200 °C |
| Scan Begin | 50 m/z | Set End Plate Offset | -500 V | Set Dry Gas | 10.0 l/min |
| Scan End | 2500 m/z | Set Collision Cell RF | 600.0 Vpp | Set Divert Valve | Waste |

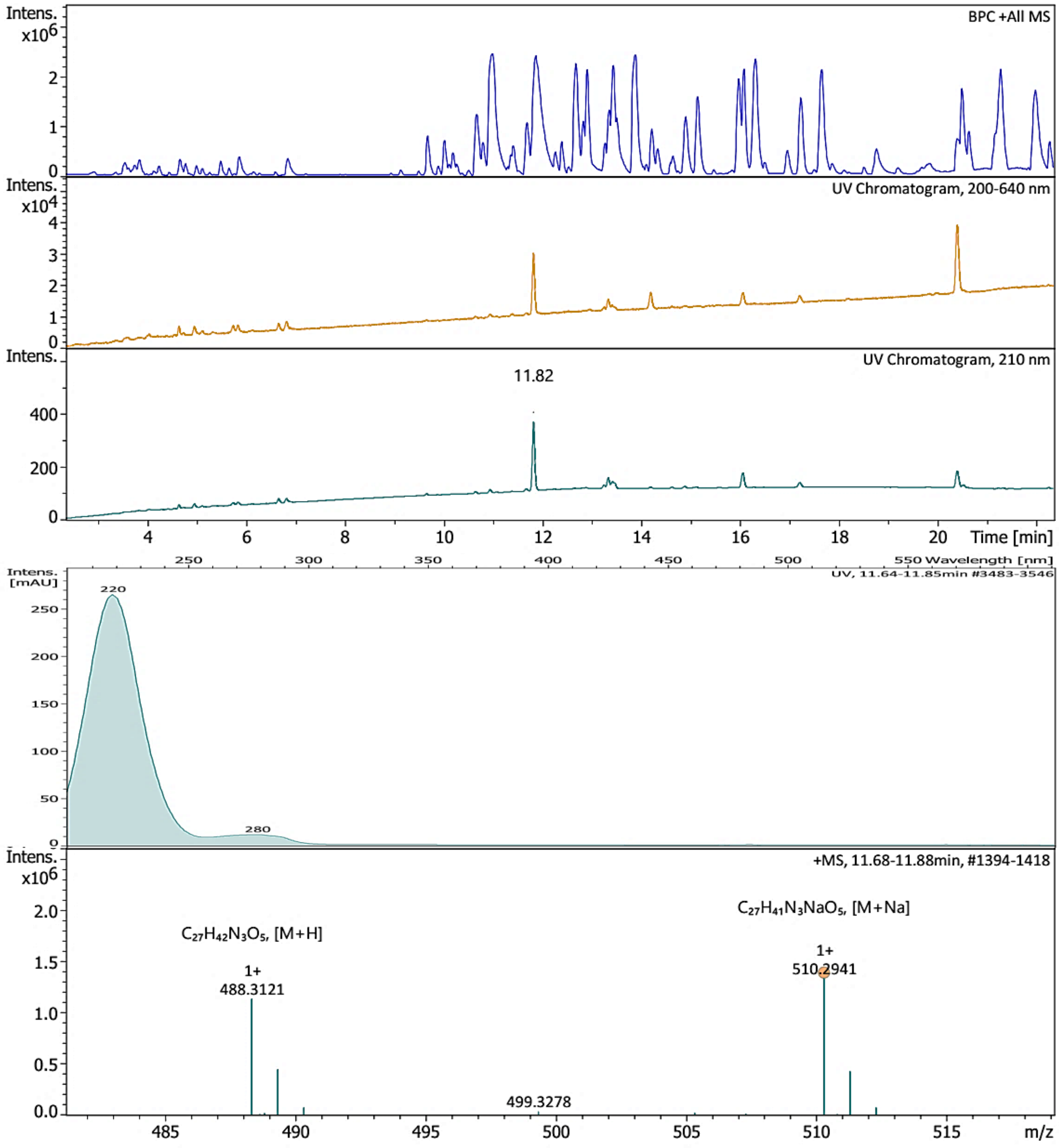


Bruker Compass DataAnalysis 6.1 printed: 13.10.2024 10:12:31 by: sel22 Page 1 of 1

Figure S17. HR-ESI-MS spectrum of **4**.

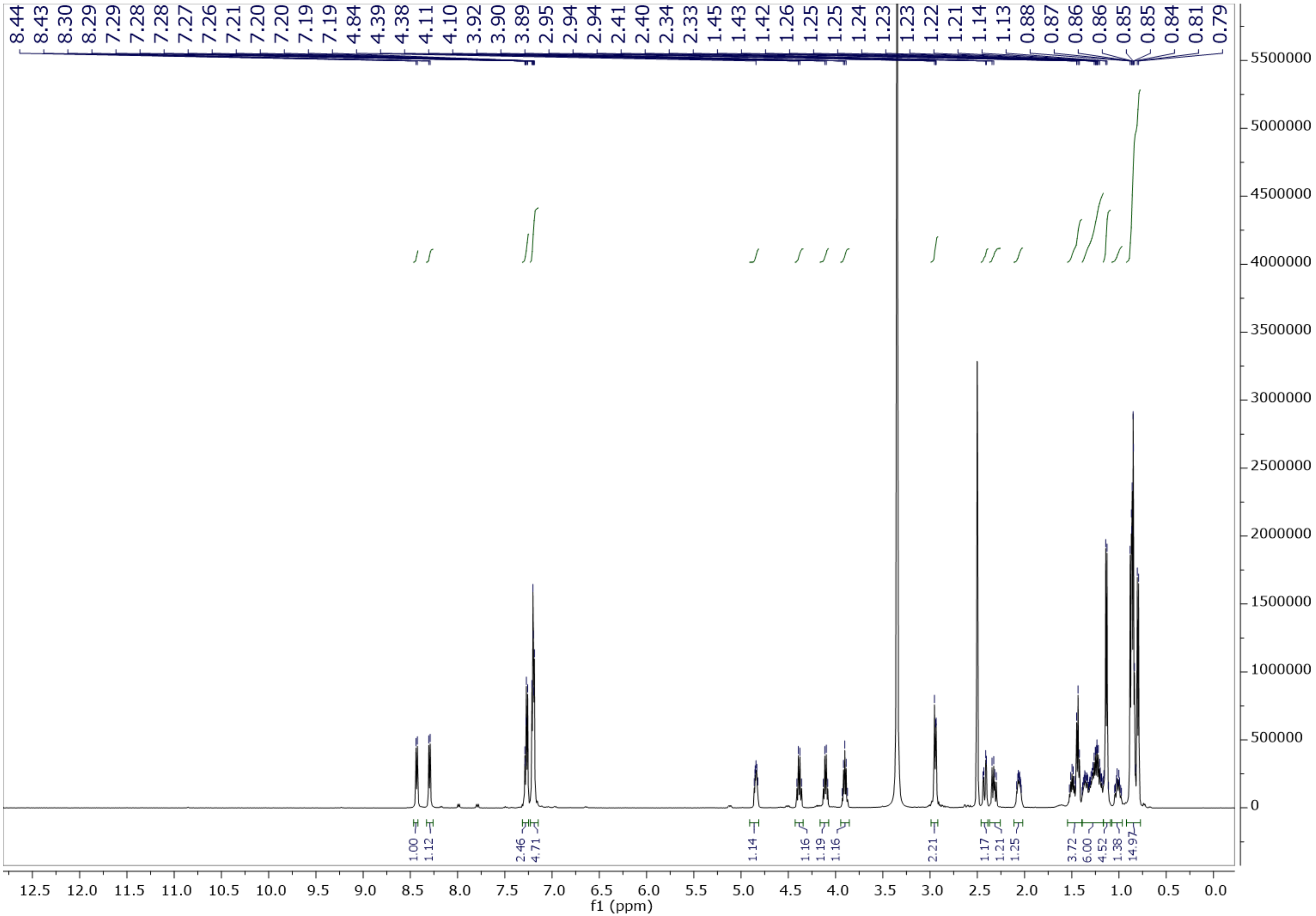

Figure S18. $^1$H NMR spectrum of **4** in DMSO-$d_6$ at 500 MHz.

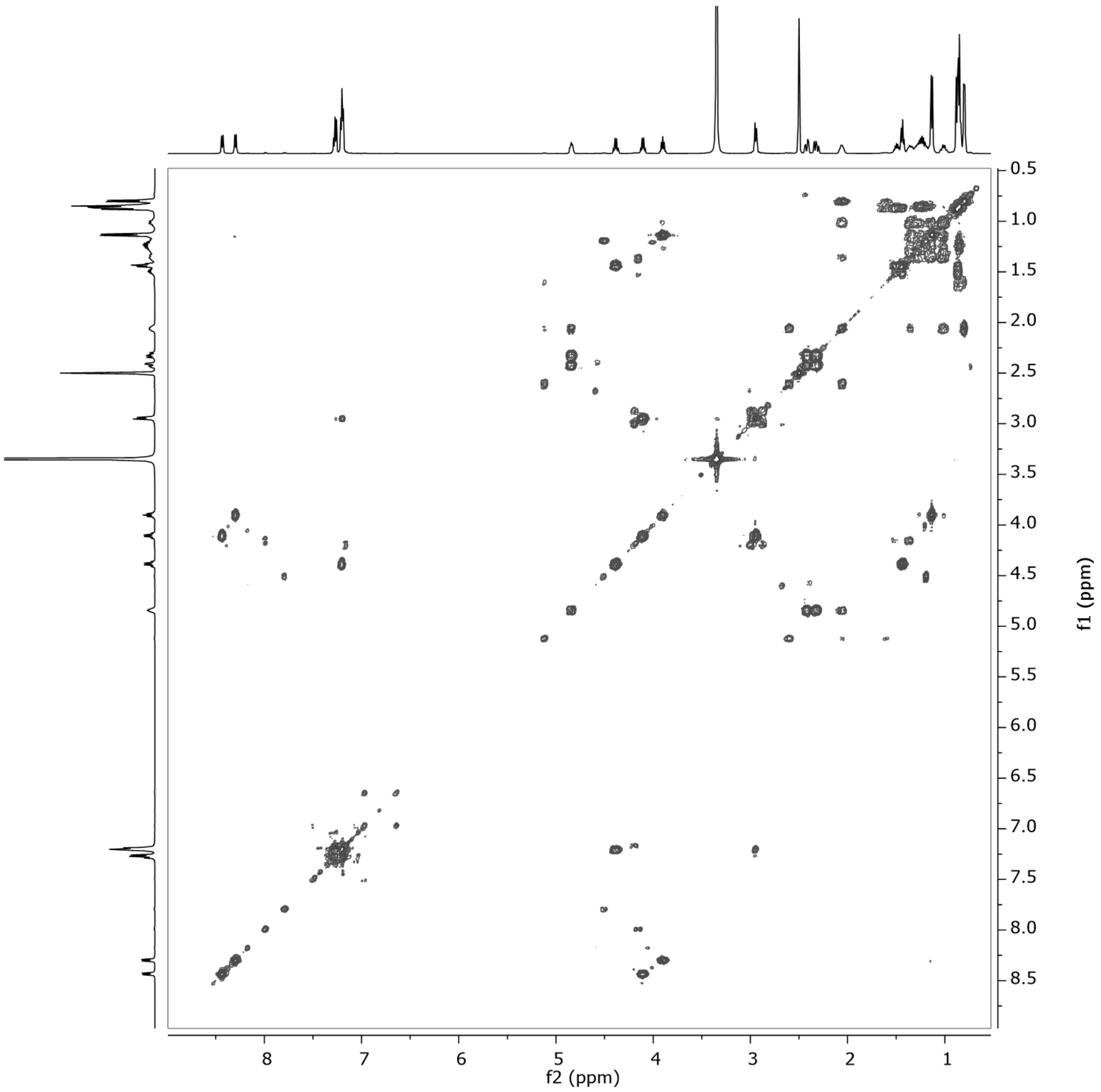


Figure S19. ${}^{1}H$–${}^{1}H$ COSY spectrum of **4** in DMSO-$d_6$ at 500 MHz.

### Generic Display Report

**Analysis Info**

| | | | |
|---|---|---|---|
| Analysis Name | S:\DATA\AmaZon\rto20_Rita Toshe\23-05\MY11508-GG1-LS1-S1-F1-F02_RC2_01_46840.d | Acquisition Date | 06.05.2023 14:23:50 |
| Method | 46840.m | Operator | tti |
| Sample Name | MY11508-GG1-LS1-S1-F1-F02 | Instrument | amaZon speed |
| Comment | | | |

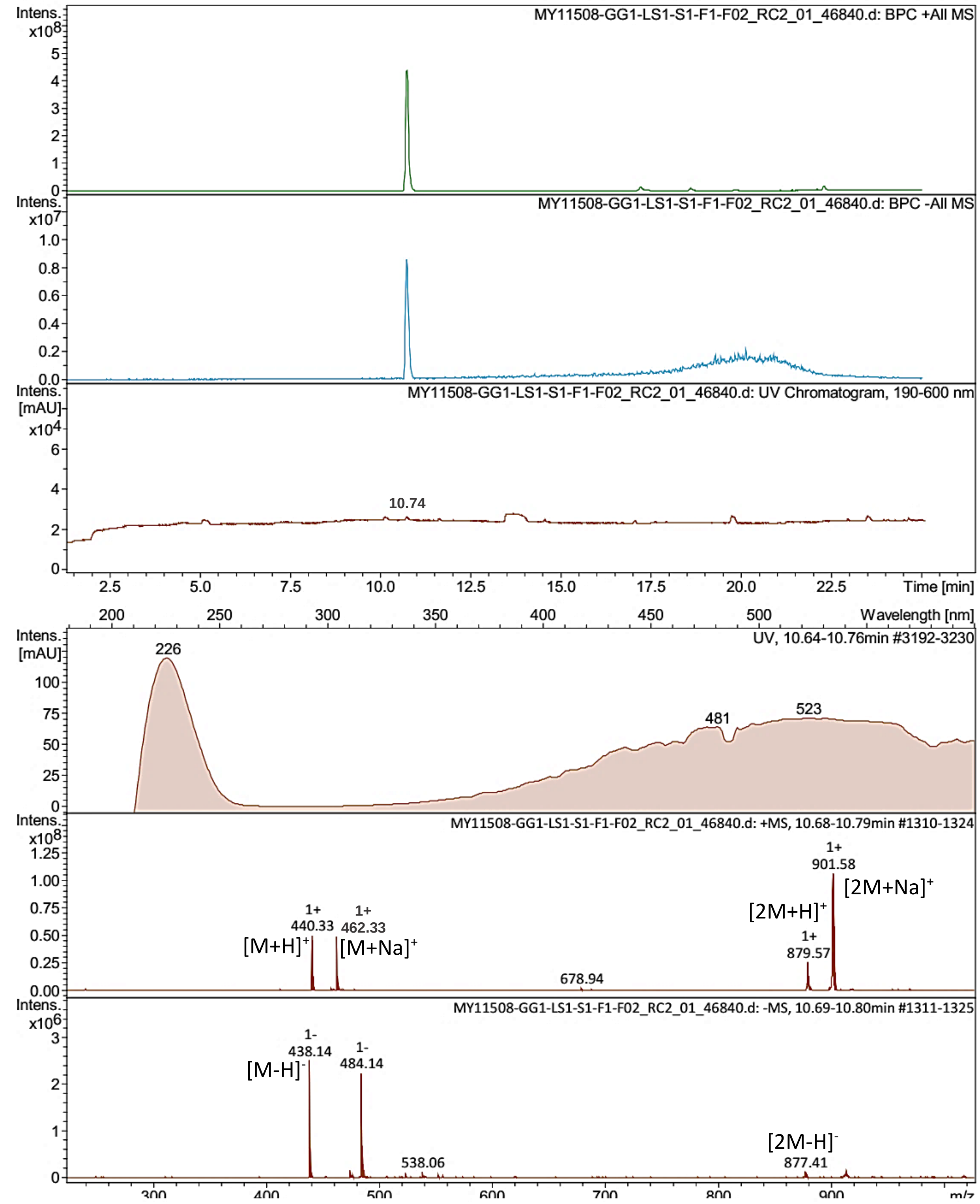


Bruker Compass DataAnalysis 4.4    printed: 09.05.2023 16:27:09    by: sel22    Page 1 of 1

Figure S20. LR-ESI-MS spectrum of **5**.

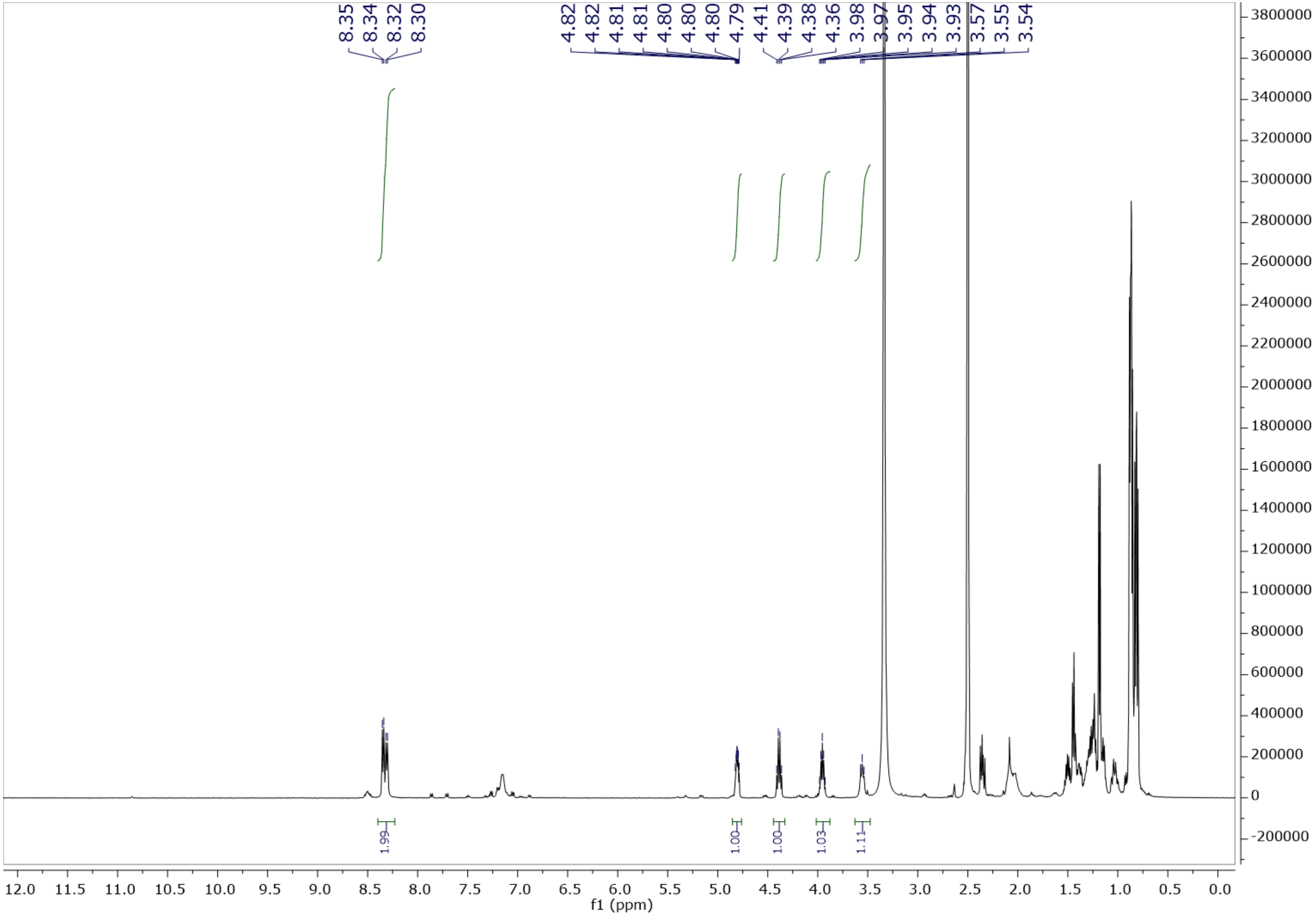


Figure S21. [1]H NMR spectrum of **5** in DMSO-$d_6$ at 500 MHz.

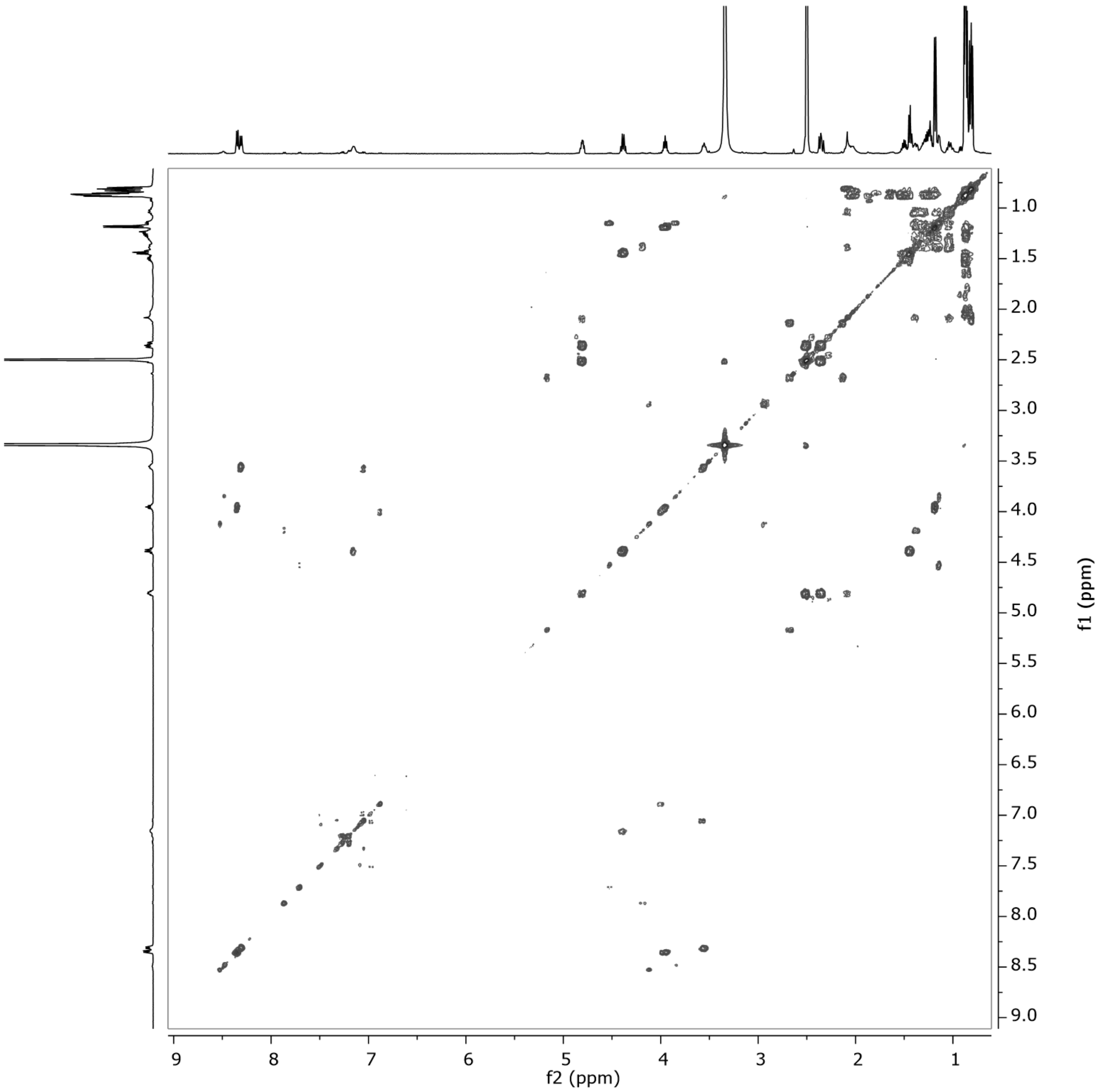


Figure S22. $^{1}H$–$^{1}H$ COSY spectrum of **5** in DMSO-$d_6$ at 500 MHz.

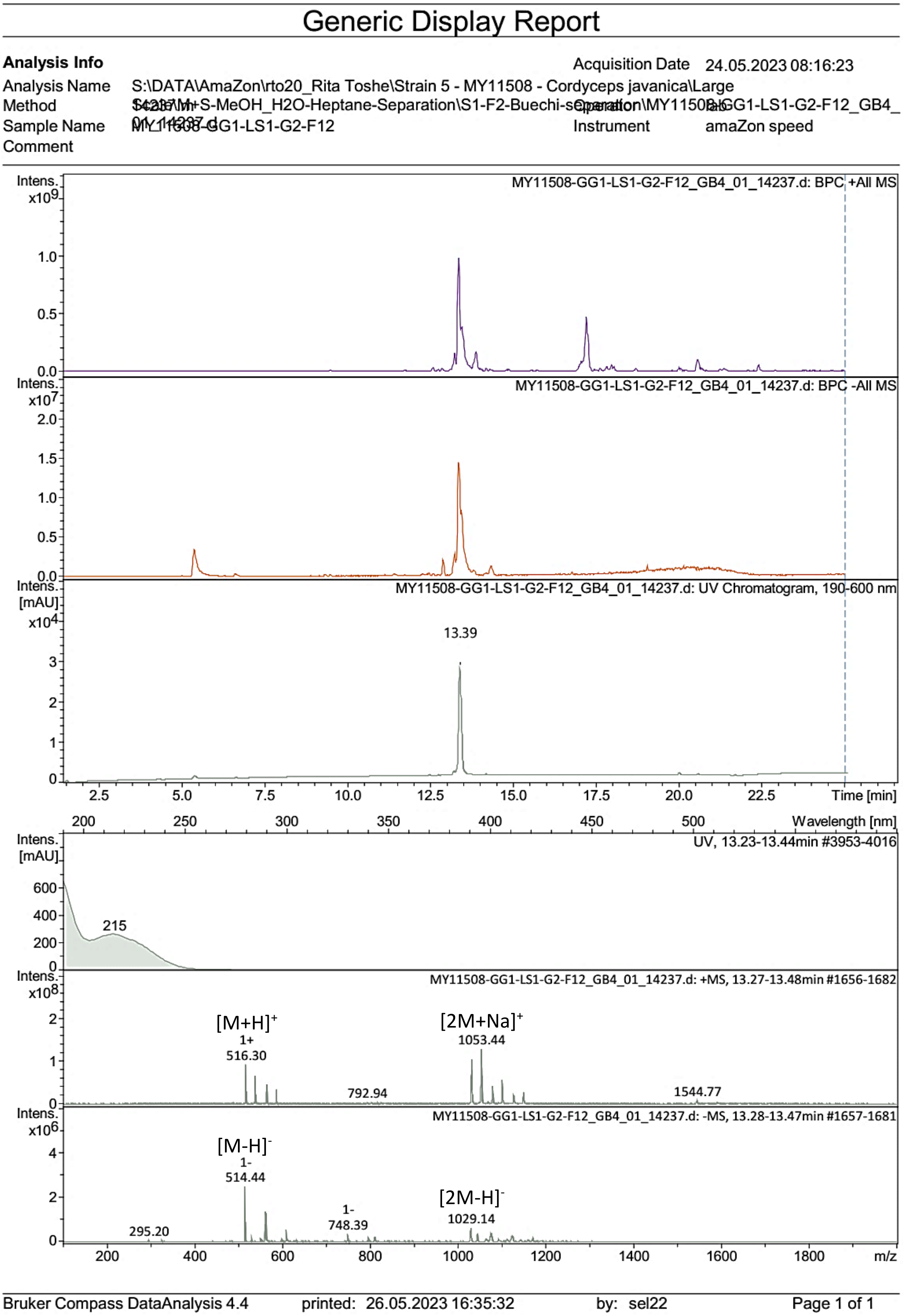


Figure S23. LR-ESI-MS spectrum of **6**.

### Generic Display Report

| | | | |
|---|---|---|---|
| **Analysis Info** | | Acquisition Date | 27.05.2023 20:06:11 |
| Analysis Name | S:\DATA\MaXis\rto20_RitaToshe\23_05\23_05_27\MY11508-GG1-LS1-F02-F12_72_01_11691.d | | |
| Method | pos_säure_10000_screening_ms_100_2500_line.m | Operator | ate06 |
| Sample Name | MY11508-GG1-LS1-F02-F12 | Instrument | maXis |
| Comment | Screening01<br>Waters Acquity UPLC BEH C_18 1,7um 2.1x50mm | | |

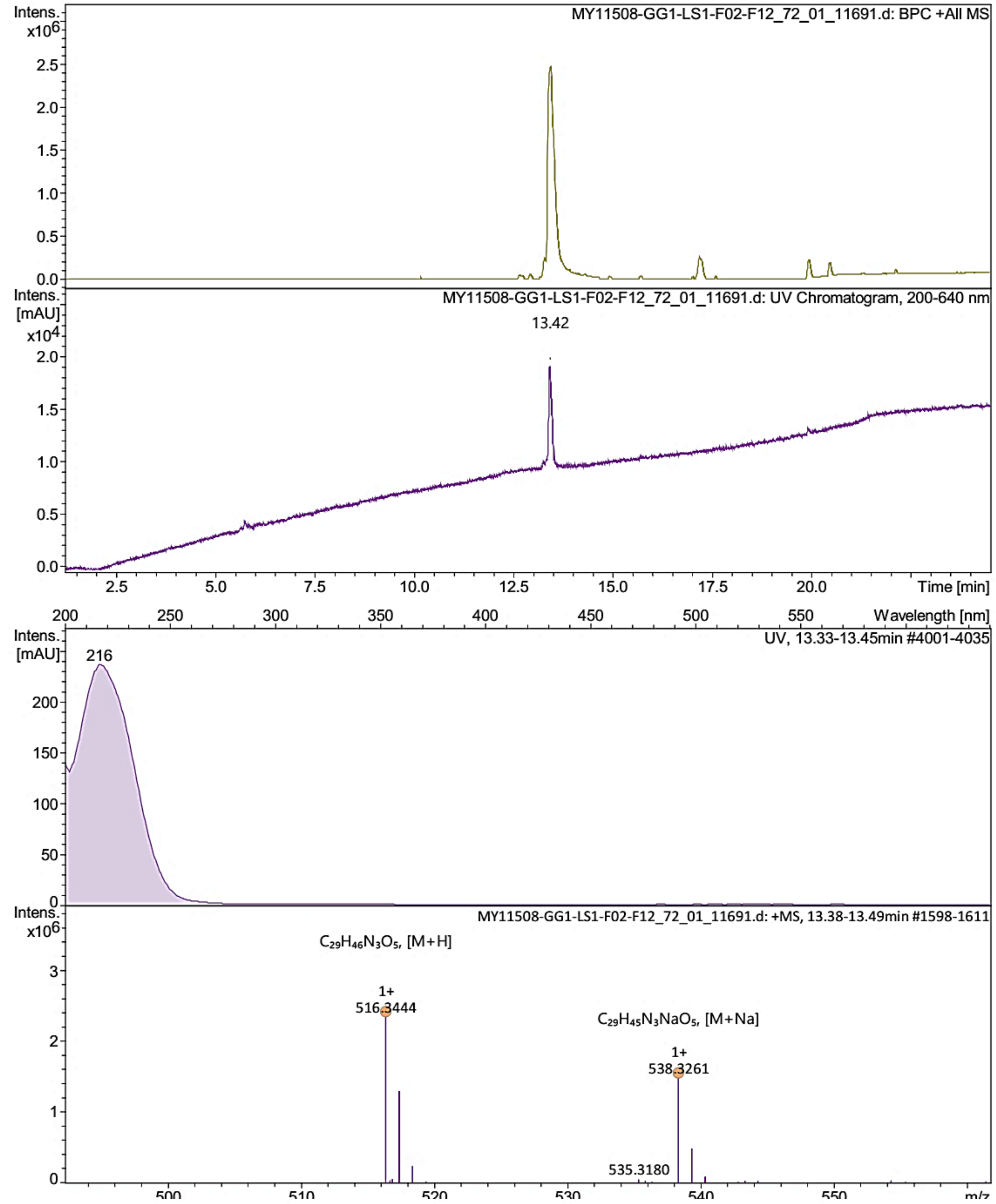


Bruker Compass DataAnalysis 4.4 printed: 02.06.2023 16:33:27 by: sel22 Page 1 of 1

Figure S24. HR-ESI-MS spectrum of **6**.

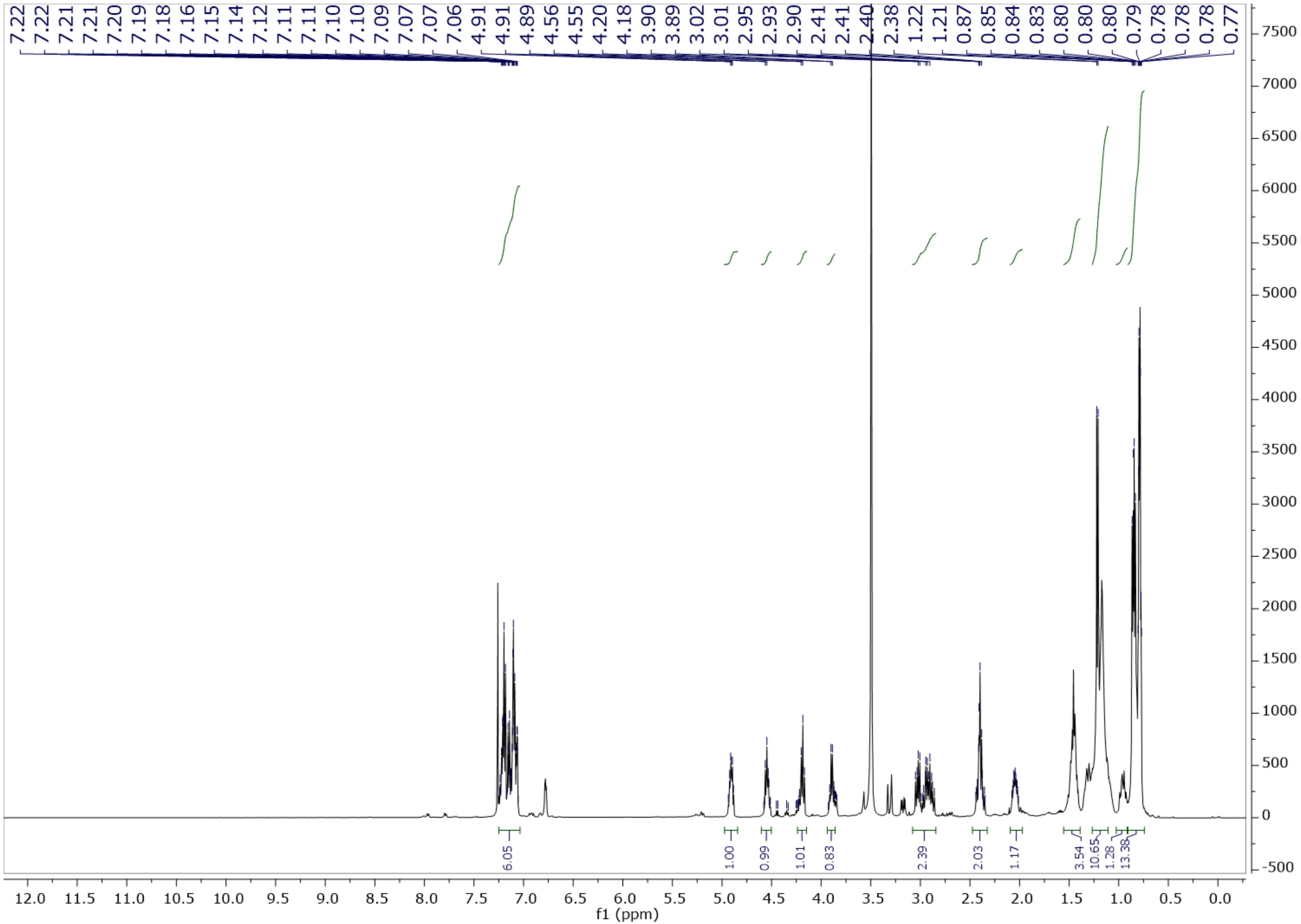


Figure S25. $^{1}$H NMR spectrum of **6** in DMSO-$d_6$ at 500 MHz.

### Generic Display Report

**Analysis Info**

| | | | |
|---|---|---|---|
| Analysis Name | S:\DATA\AmaZon\rto20_Rita Toshe\Strain 5 - MY11508 - Cordyceps javanica\Large | Acquisition Date | 23.05.2023 22:36:46 |
| Method | Scale M+S-MeOH_H2O-Heptane-Separation\S1-F2-Buechi-separation\MY11508-GG1-LS1-G2-F10_GB2_01_14221.d | Operator | lab |
| Sample Name | MY11508-GG1-LS1-G2-F10 | Instrument | amaZon speed |
| Comment | | | |

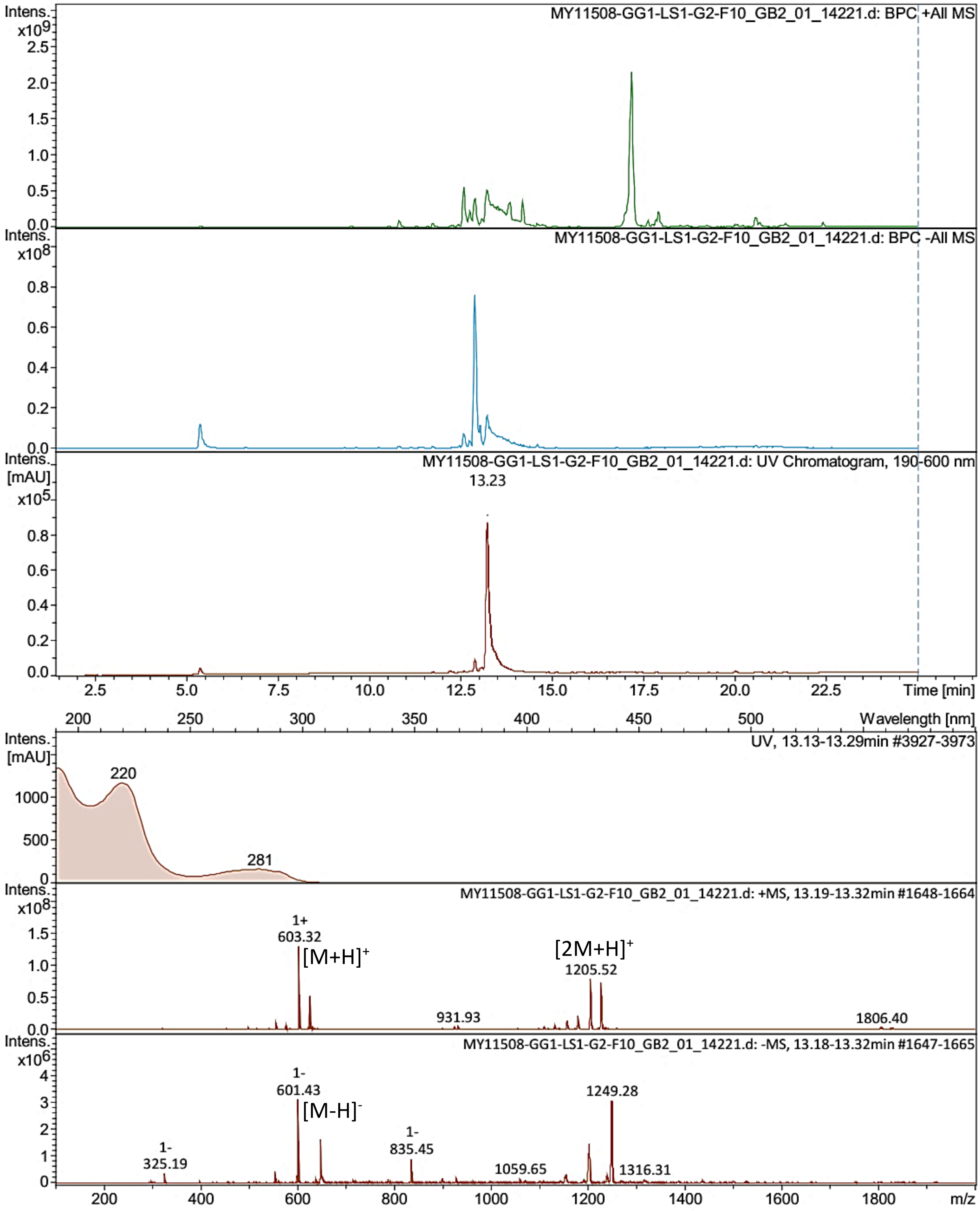




Figure S26. LR-ESI-MS spectrum of **7**.

### Generic Display Report

**Analysis Info**

| | |
|---|---|
| Analysis Name | S:\DATA\MaXis\rto20_RitaToshe\23_05\23_05_27\MY11508-GG1-LS1-F02-F10_70_01_11689.d |
| Method | pos_säure_10000_screening_ms_100_2500_line.m |
| Sample Name | MY11508-GG1-LS1-F02-F10 |
| Comment | Screening01<br>Waters Acquity UPLC BEH C_18 1,7um 2.1x50mm |

| | |
|---|---|
| Acquisition Date | 27.05.2023 19:04:12 |
| Operator | ate06 |
| Instrument | maXis |

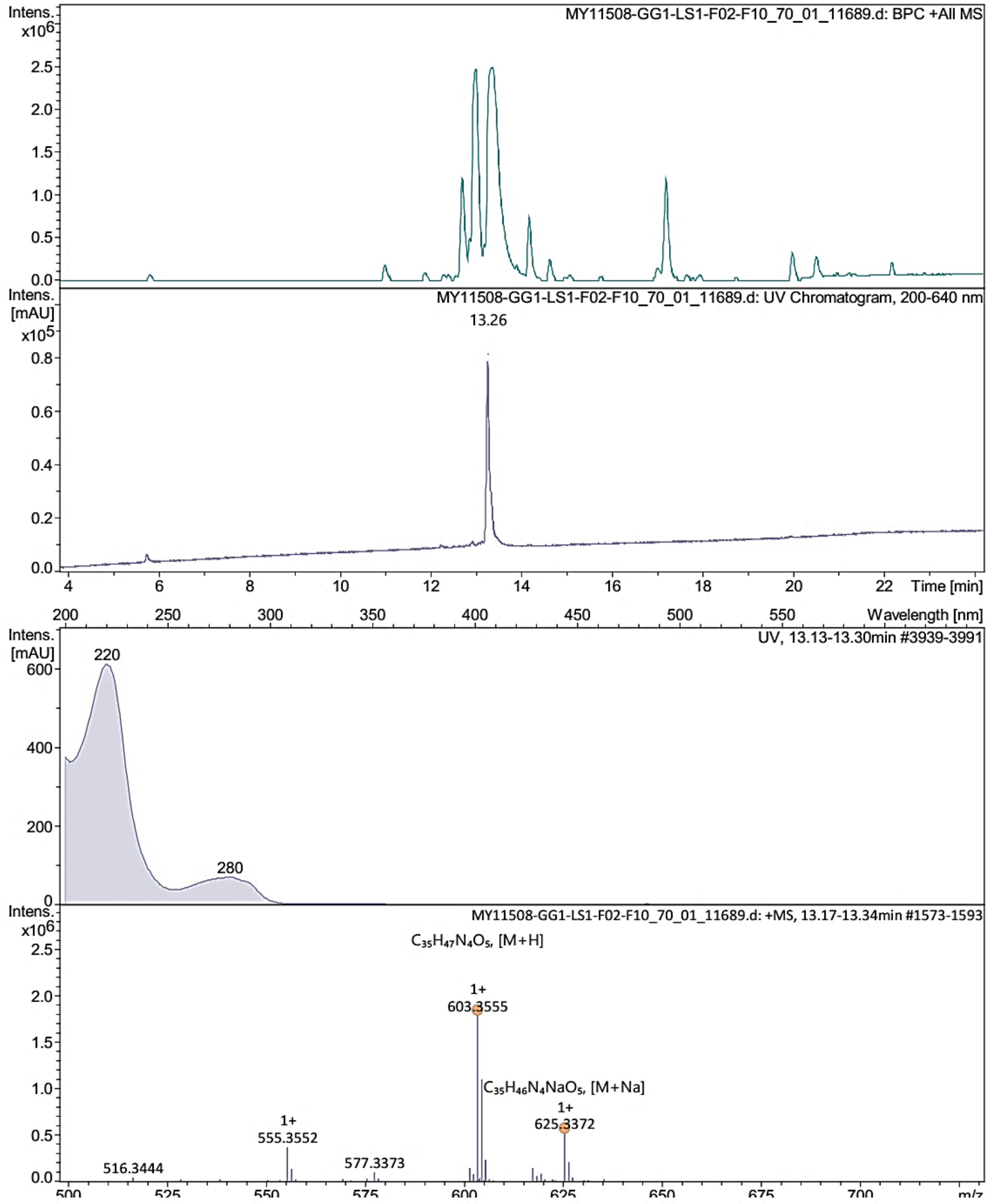


Bruker Compass DataAnalysis 4.4 printed: 01.06.2023 10:45:41 by: sel22 Page 1 of 1

Figure S27. HR-ESI-MS spectrum of **7**.

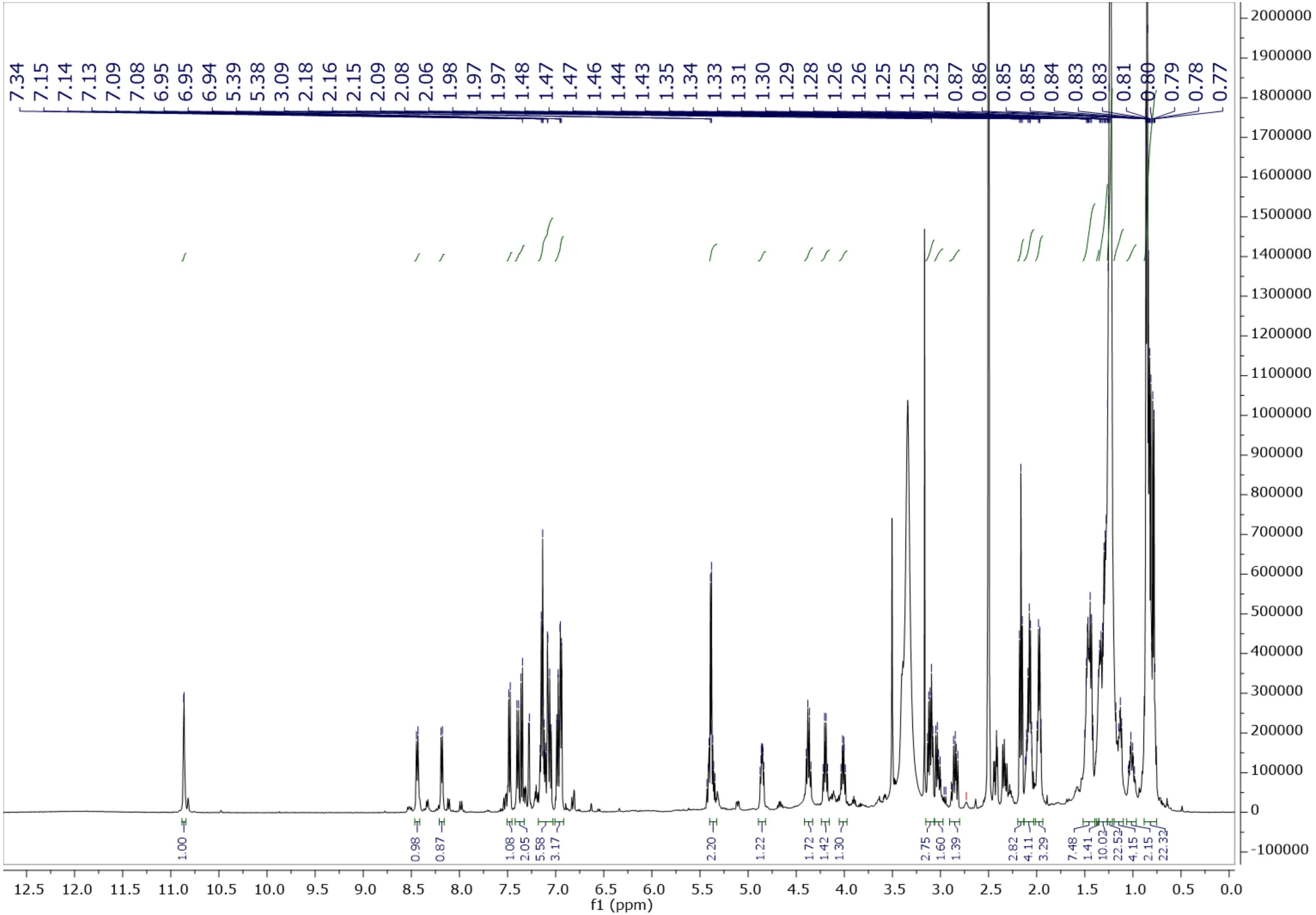

Figure S28. $^{1}$H NMR spectrum of **7** in DMSO-$d_6$ at 500 MHz.

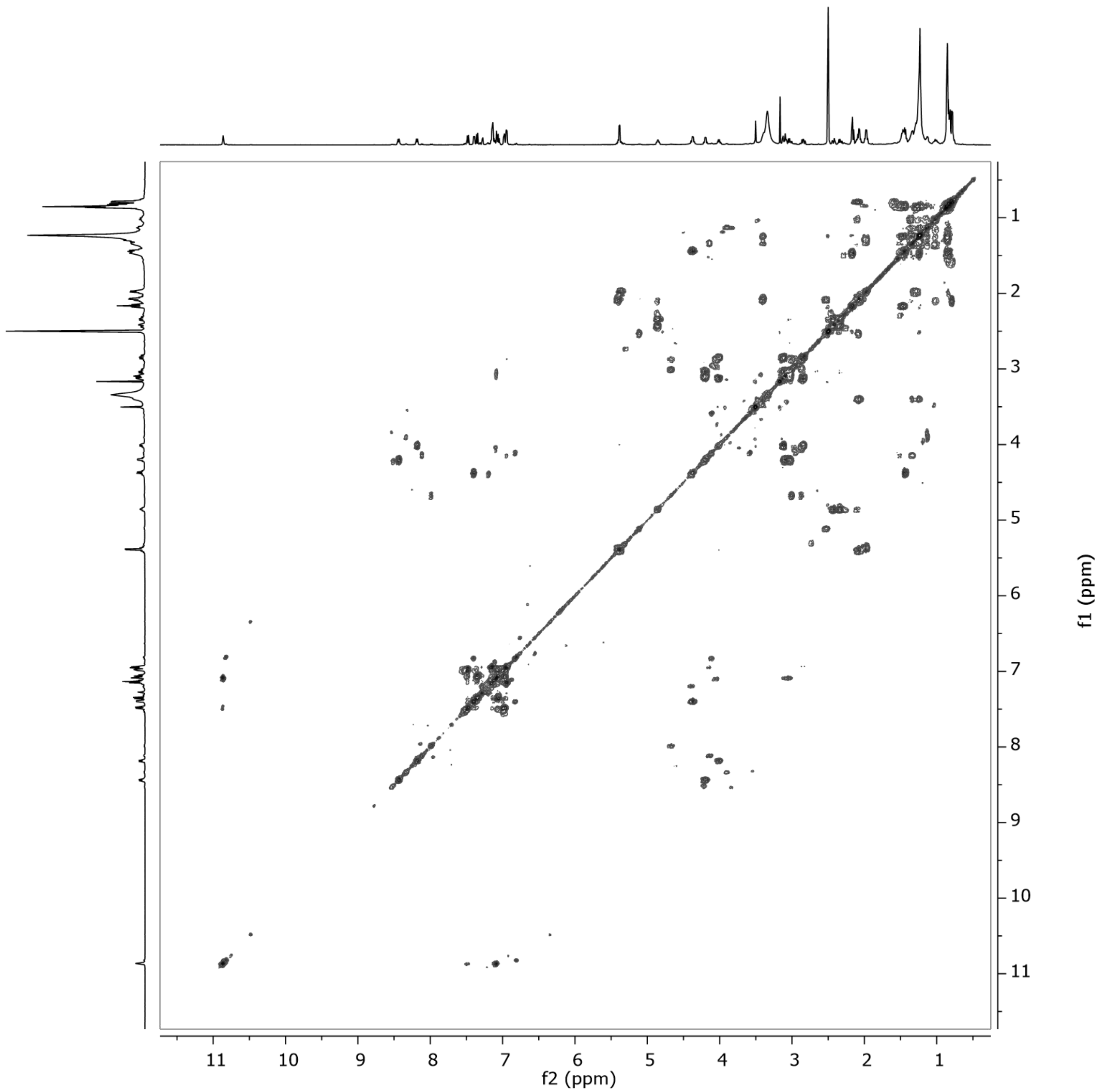


Figure S29. $^1H$–$^1H$ COSY spectrum of **7** in DMSO-$d_6$ at 500 MHz.

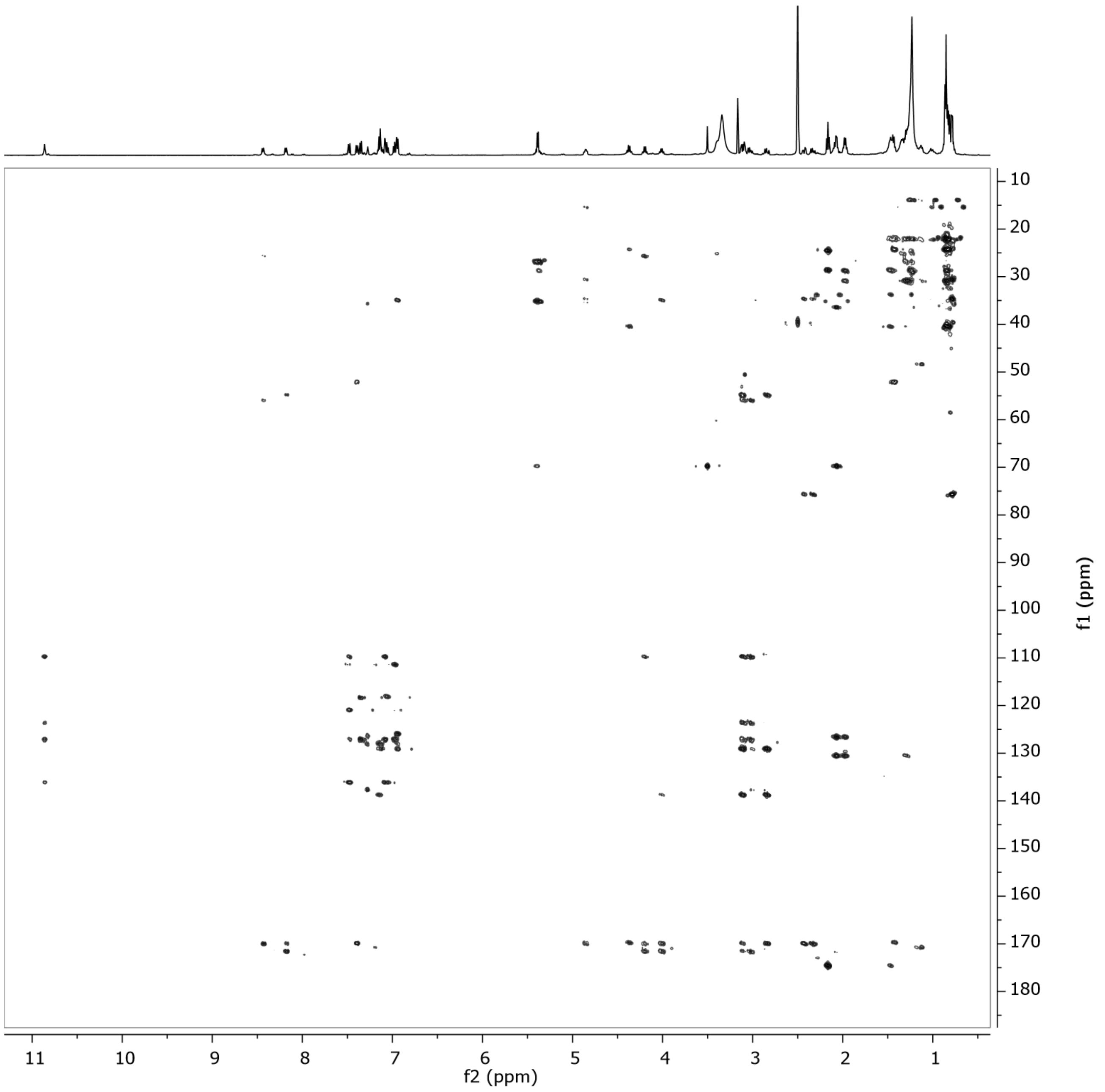


Figure S30. HMBC spectrum of **7** in DMSO-$d_6$ at 500 MHz.

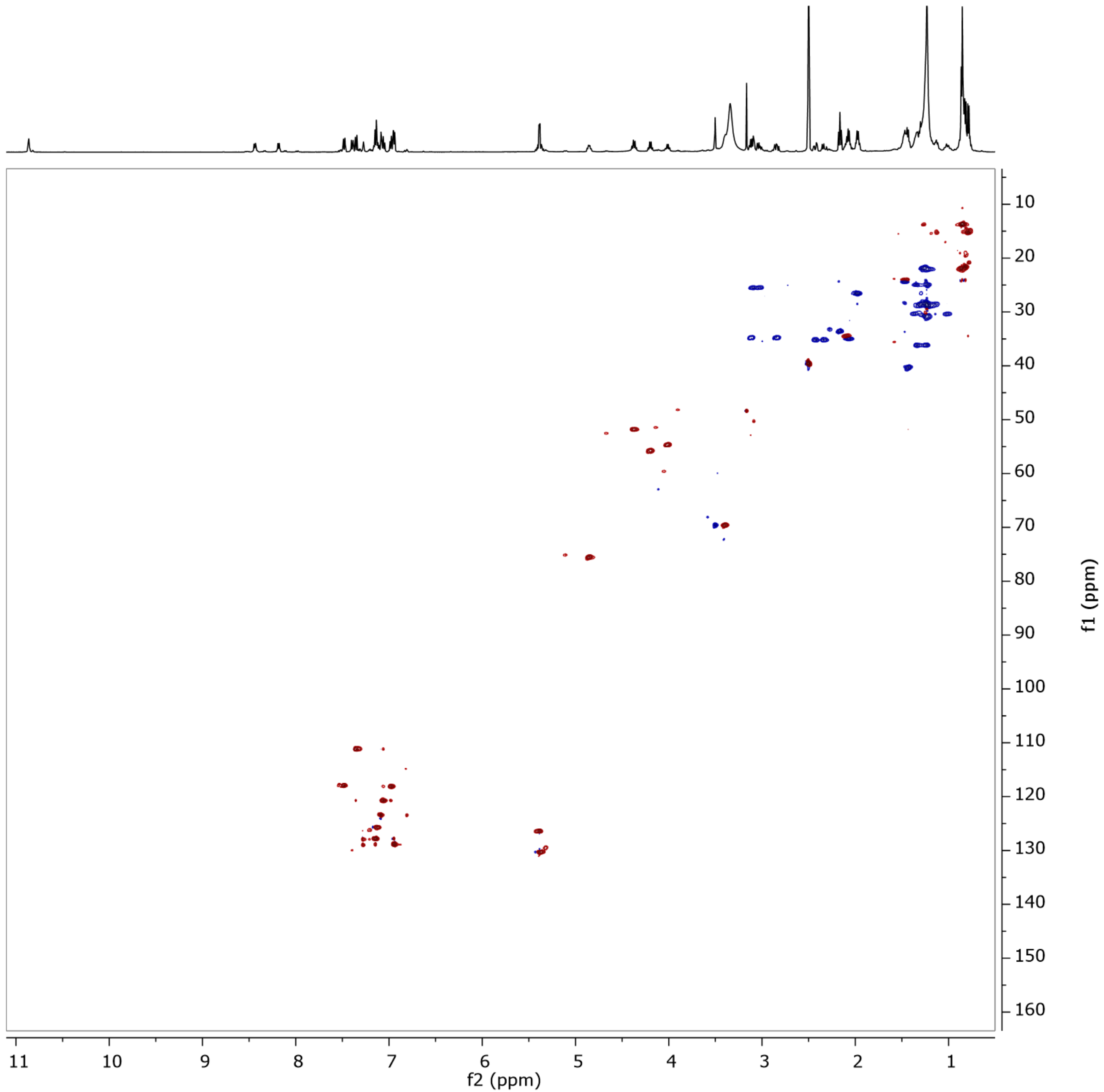


Figure S31. HSQC spectrum of **7** in DMSO-$d_6$ at 500 MHz.

### Display Report

**Analysis Info**

| | | | |
|---|---|---|---|
| Analysis Name | S:\DATA\AmaZon\rto20_Rita Toshe\Hit extracts\MY04954-YM-Cordyceps blackwelliae\MY04954-YM-S-F12_RC5_01_14540.d | Acquisition Date | 20.06.2023 10:00:58 |
| Method | 14540.m | Operator | lab |
| Sample Name | MY04954-YM-S-F12 | Instrument | amaZon speed 8374444.06119 |
| Comment | | | |

**Acquisition Parameter**

Ion Polarity Positive

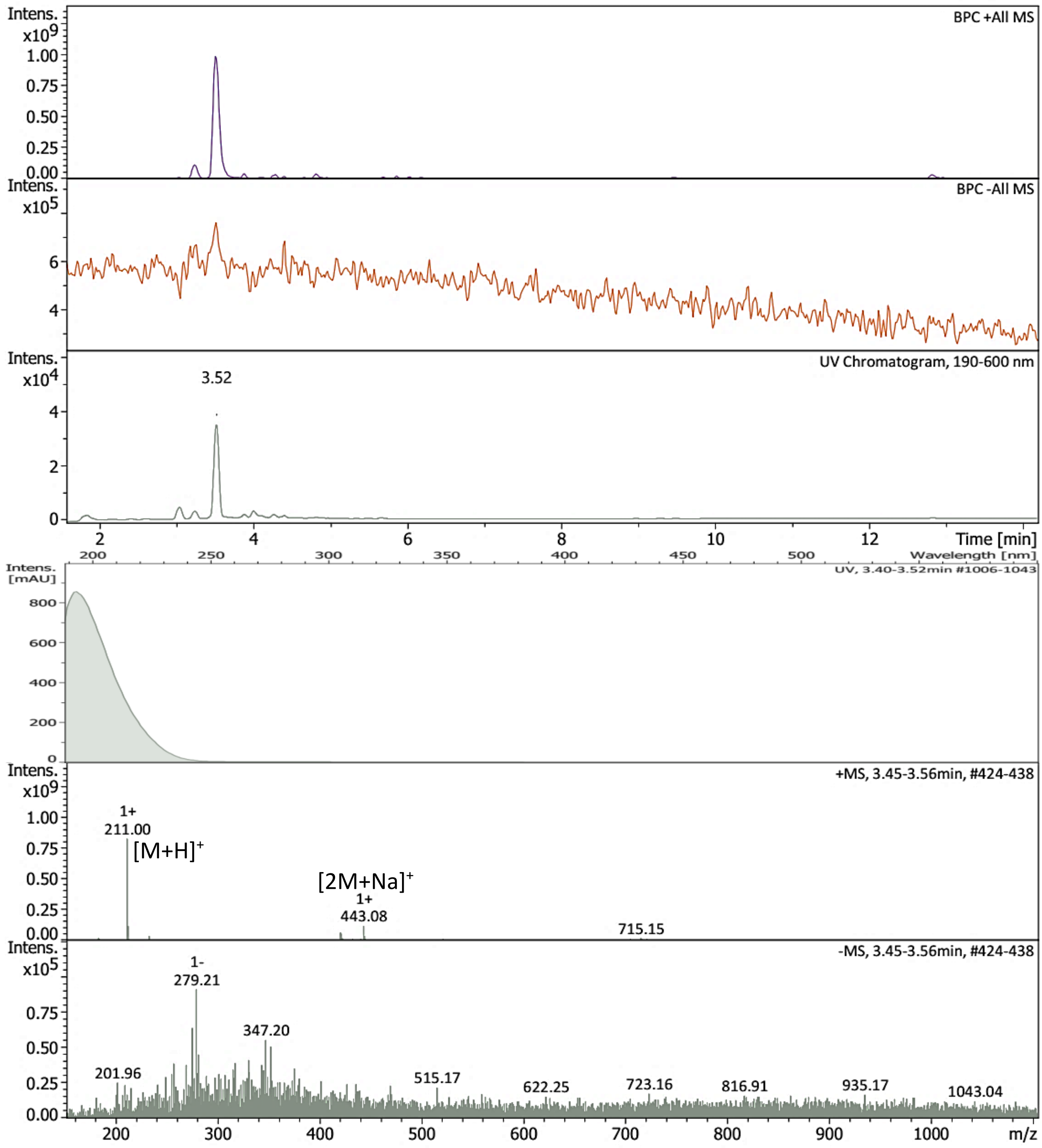


Bruker Compass DataAnalysis 6.1 printed: 13.10.2024 13:36:41 by: sel22 Page 1 of 1

Figure S32. LR-ESI-MS spectrum of **8**.

### Display Report

**Analysis Info**

| | | | |
|---|---|---|---|
| Analysis Name | S:\DATA\MaXis\rto20_Rita Toshe\Hit extracts\MY04954-YM-Cordyceps blackwelliae\MY04954-YM-S-F12_20_01_12018.d | Acquisition Date | 27.06.2023 20:57:48 |
| Method | pos_säure_10000_screening_ms_100_2500_line.m | Operator | ate06 |
| Sample Name | MY04954-YM-S-F12 | Instrument | maXis 255552.00037 |
| Comment | Screening01<br>Waters Acquity UPLC BEH C_18 1,7um 2.1x50mm | | |

**Acquisition Parameter**

| | | | | | |
|---|---|---|---|---|---|
| Source Type | ESI | Ion Polarity | Positive | Set Nebulizer | 4.0 Bar |
| Focus | Not active | Set Capillary | 4500 V | Set Dry Heater | 200 °C |
| Scan Begin | 50 m/z | Set End Plate Offset | -500 V | Set Dry Gas | 10.0 l/min |
| Scan End | 2500 m/z | Set Collision Cell RF | 600.0 Vpp | Set Divert Valve | Waste |

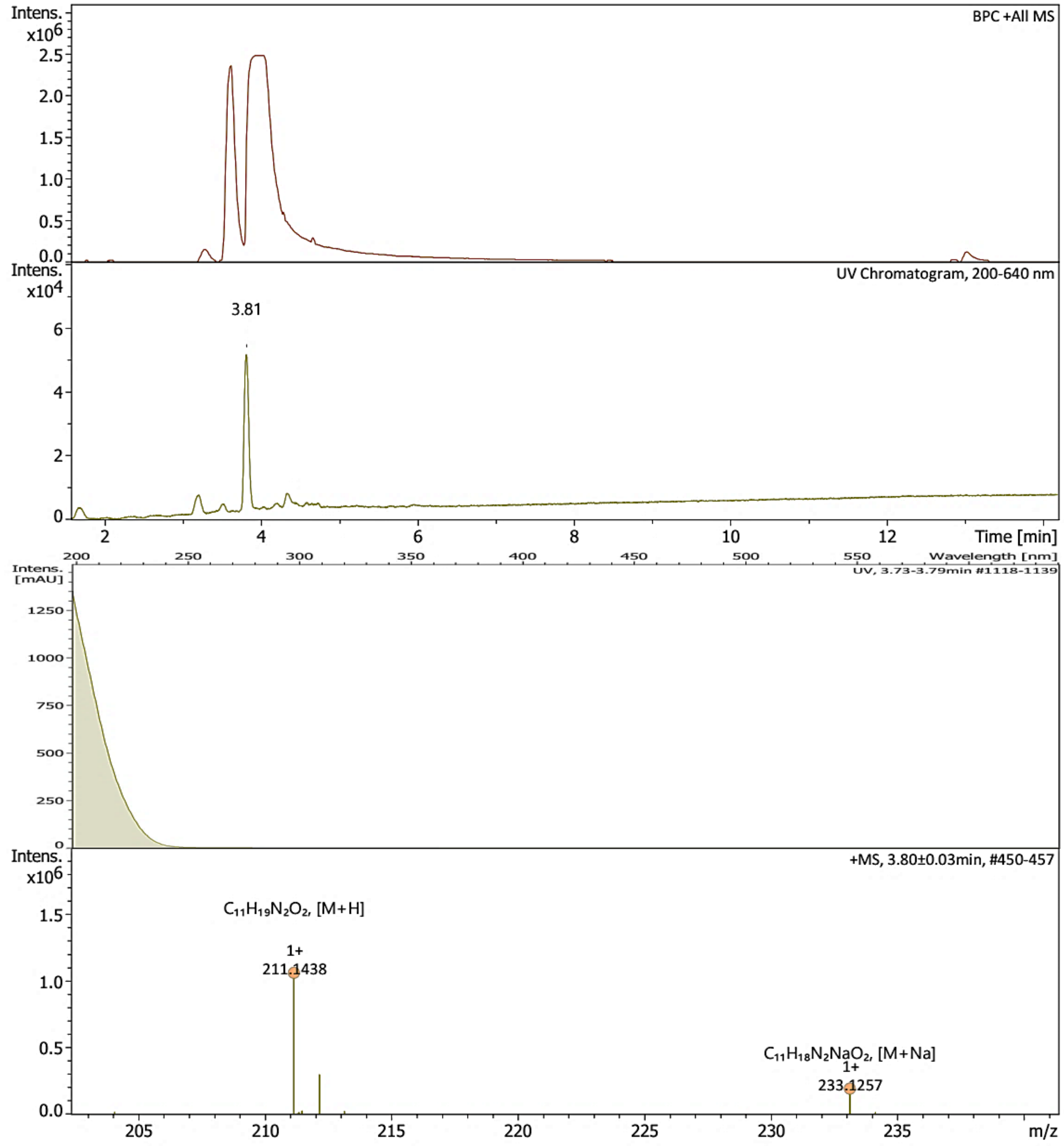




Figure S33. HR-ESI-MS spectrum of **8**.

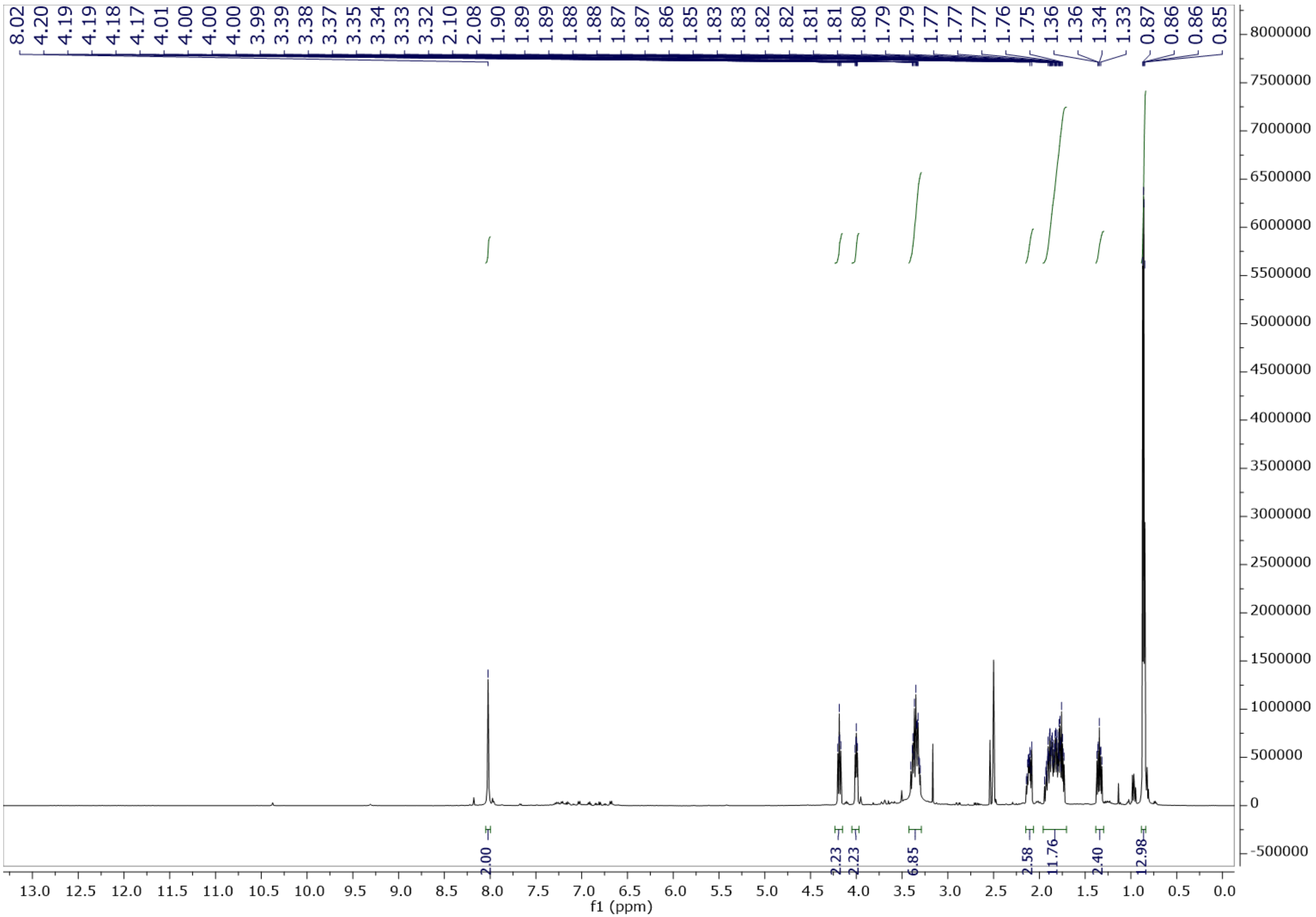


Figure S34. $^{1}$H NMR spectrum of **8** in DMSO-$d_6$ at 500 MHz.

### Generic Display Report

**Analysis Info**

| | | | |
|---|---|---|---|
| | | Acquisition Date | 20.06.2023 09:24:47 |
| Analysis Name | S:\DATA\AmaZon\rto20_Rita Toshe\23_06\MY04954-YM-S-F11_RC4_01_14539.d | | |
| Method | 14539.m | Operator | lab |
| Sample Name | MY04954-YM-S-F11 | Instrument | amaZon speed |
| Comment | | | |

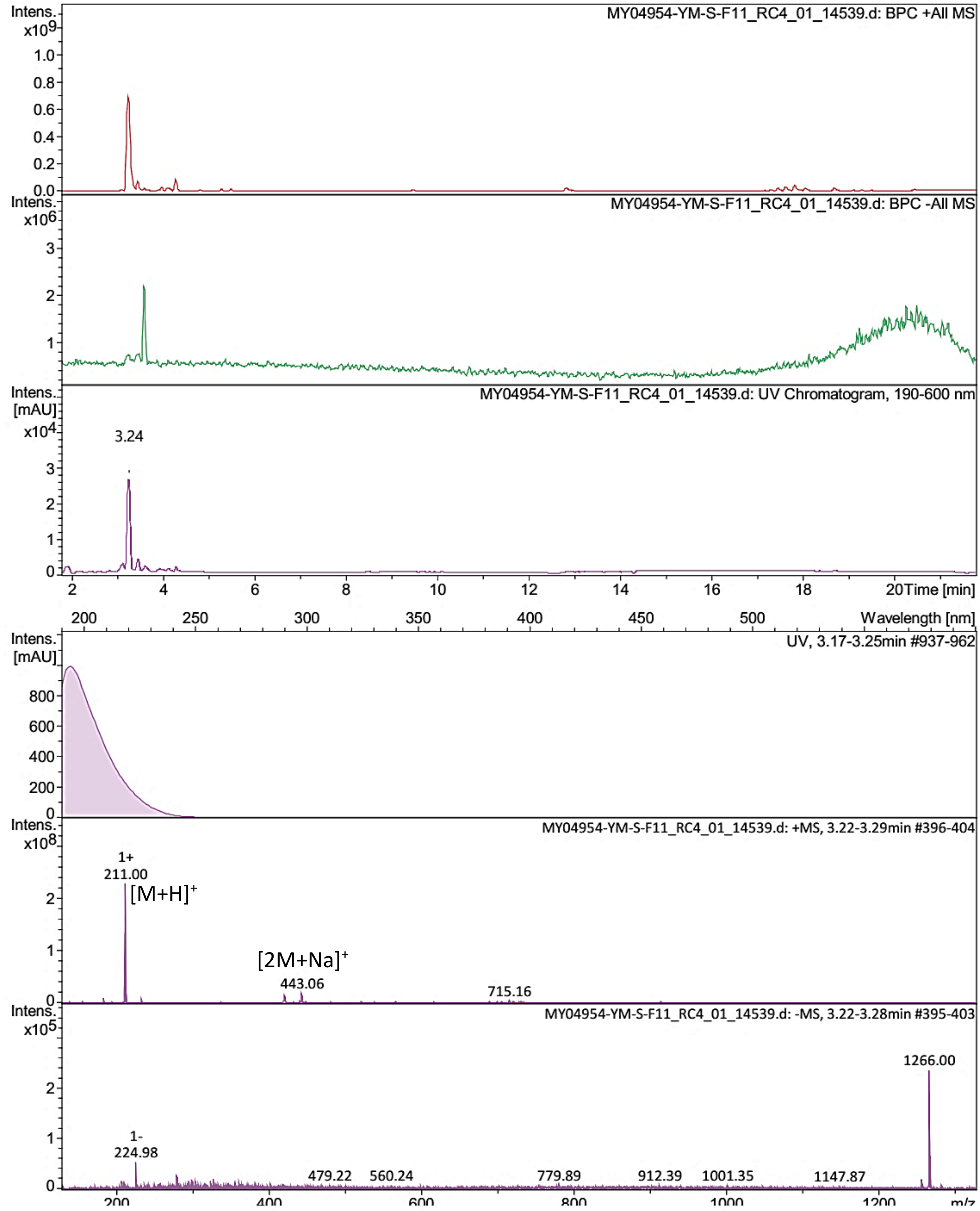


Bruker Compass DataAnalysis 4.4 printed: 21.06.2023 16:59:58 by: sel22 Page 1 of 1

Figure S35. LR-ESI-MS spectrum of **9**.

### Display Report

**Analysis Info**

| | | | |
|---|---|---|---|
| Analysis Name | S:\DATA\MaXis\rto20_Rita Toshe\23_06\MY04954-YM-S-F12_20_01_12018.d | Acquisition Date | 27.06.2023 20:57:48 |
| Method | pos_säure_10000_screening_ms_100_2500_line.m | Operator | ate06 |
| Sample Name | MY04954-YM-S-F12 | Instrument | maXis |
| Comment | Screening01<br>Waters Acquity UPLC BEH C_18 1,7um 2.1x50mm | | |

**Acquisition Parameter**

Ion Polarity Positive

SPS Target Mass

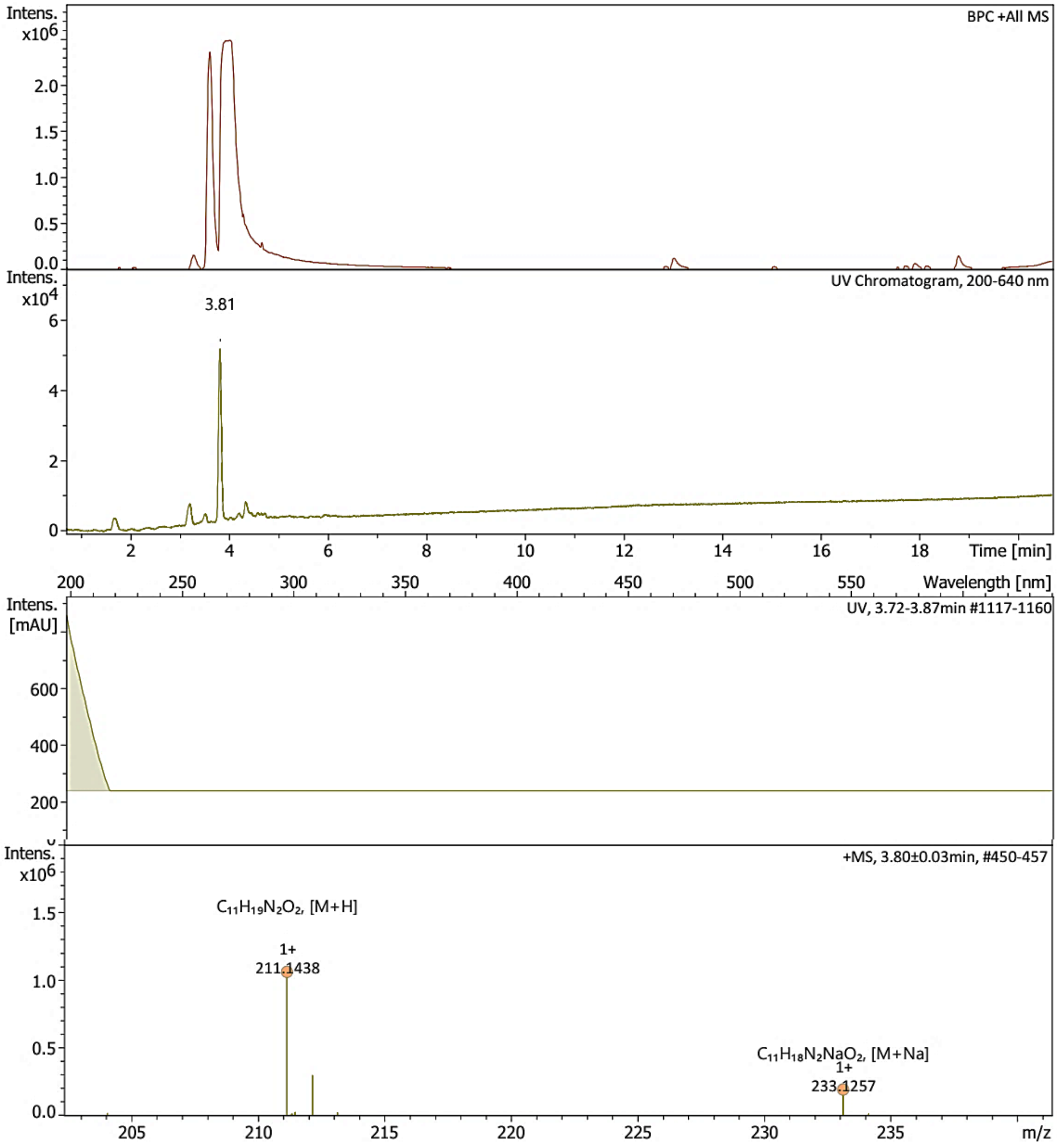


Bruker Compass DataAnalysis 6.1 printed: 13.10.2024 13:34:53 by: sel22 Page 1 of 1

Figure S36. HR-ESI-MS spectrum of **9**.

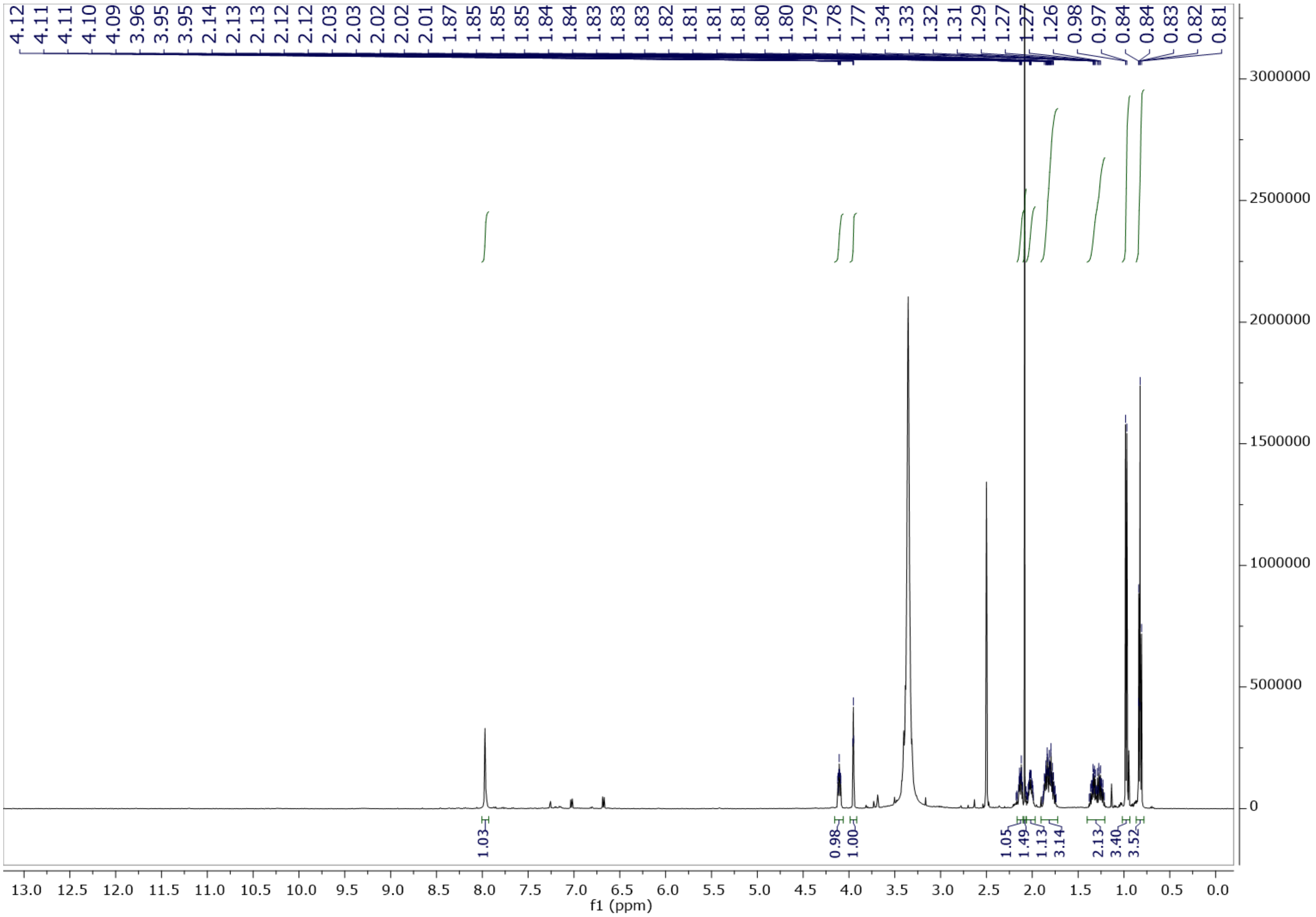


Figure S37. $^{1}H$ NMR spectrum of **9** in DMSO-$d_6$ at 500 MHz.

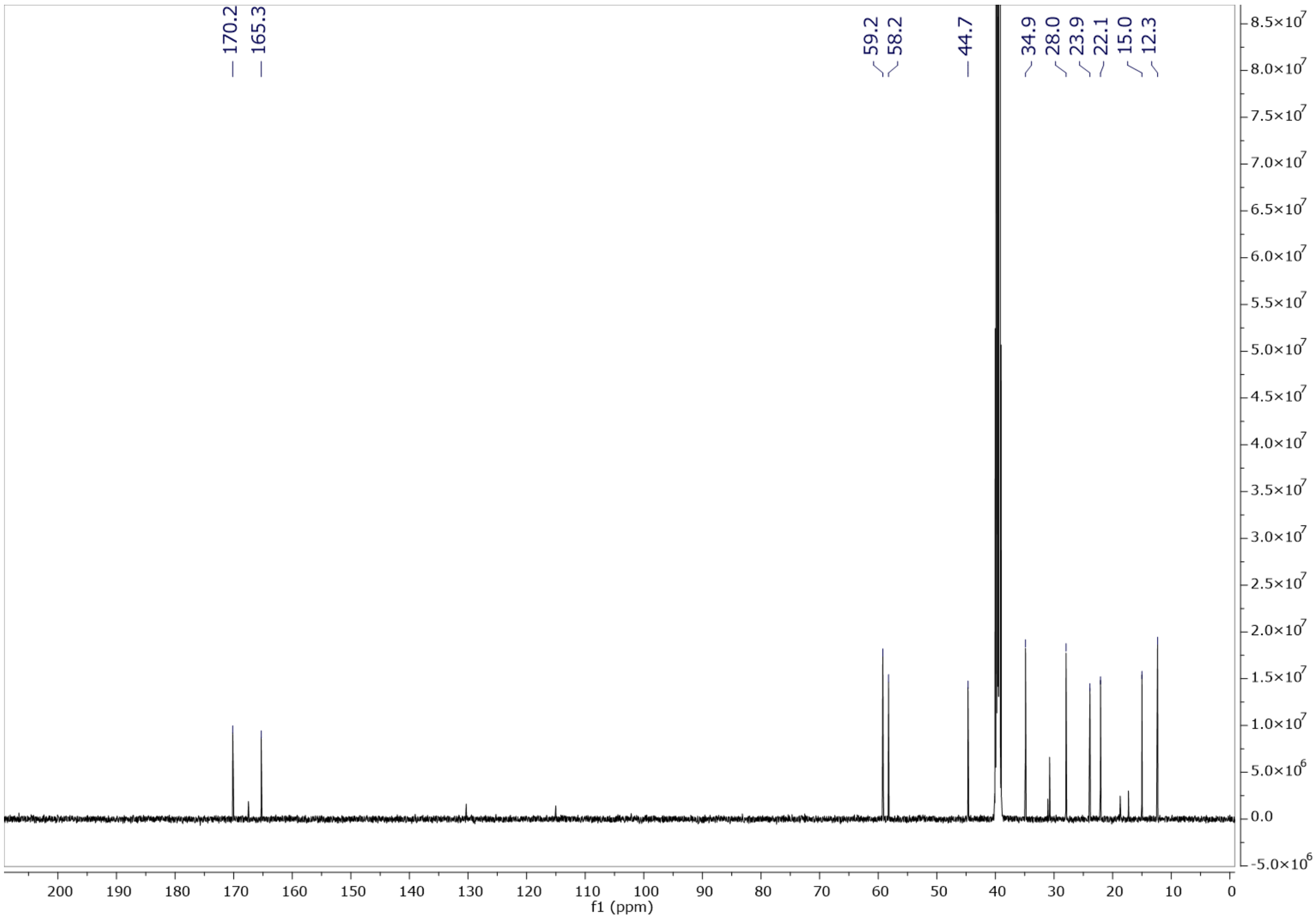


Figure S38. $^{13}C$ NMR spectrum of **9** in DMSO-$d_6$ at 125 MHz.

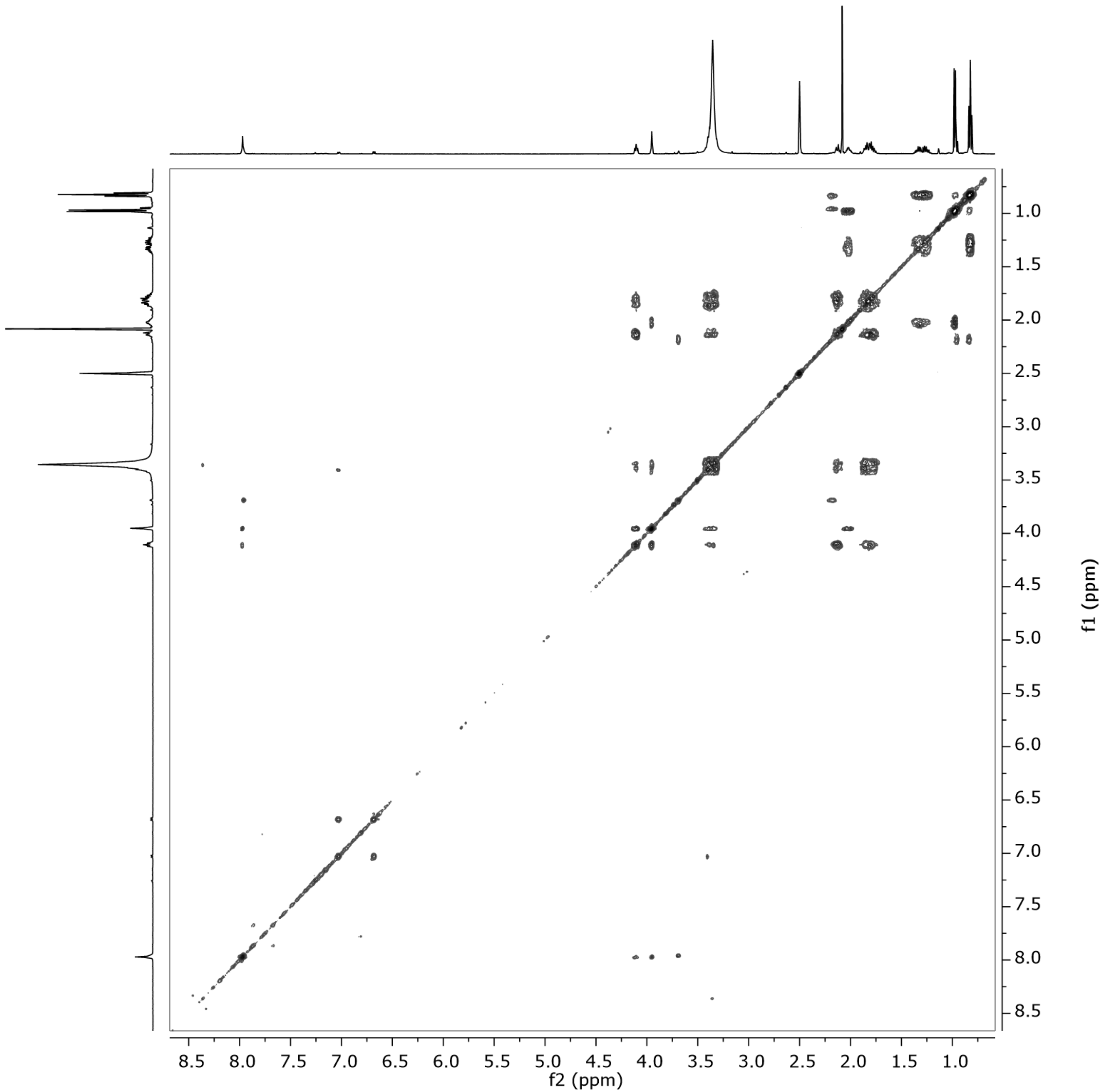


Figure S39. $^{1}H$–$^{1}H$ COSY spectrum of **9** in DMSO-$d_6$ at 500 MHz.

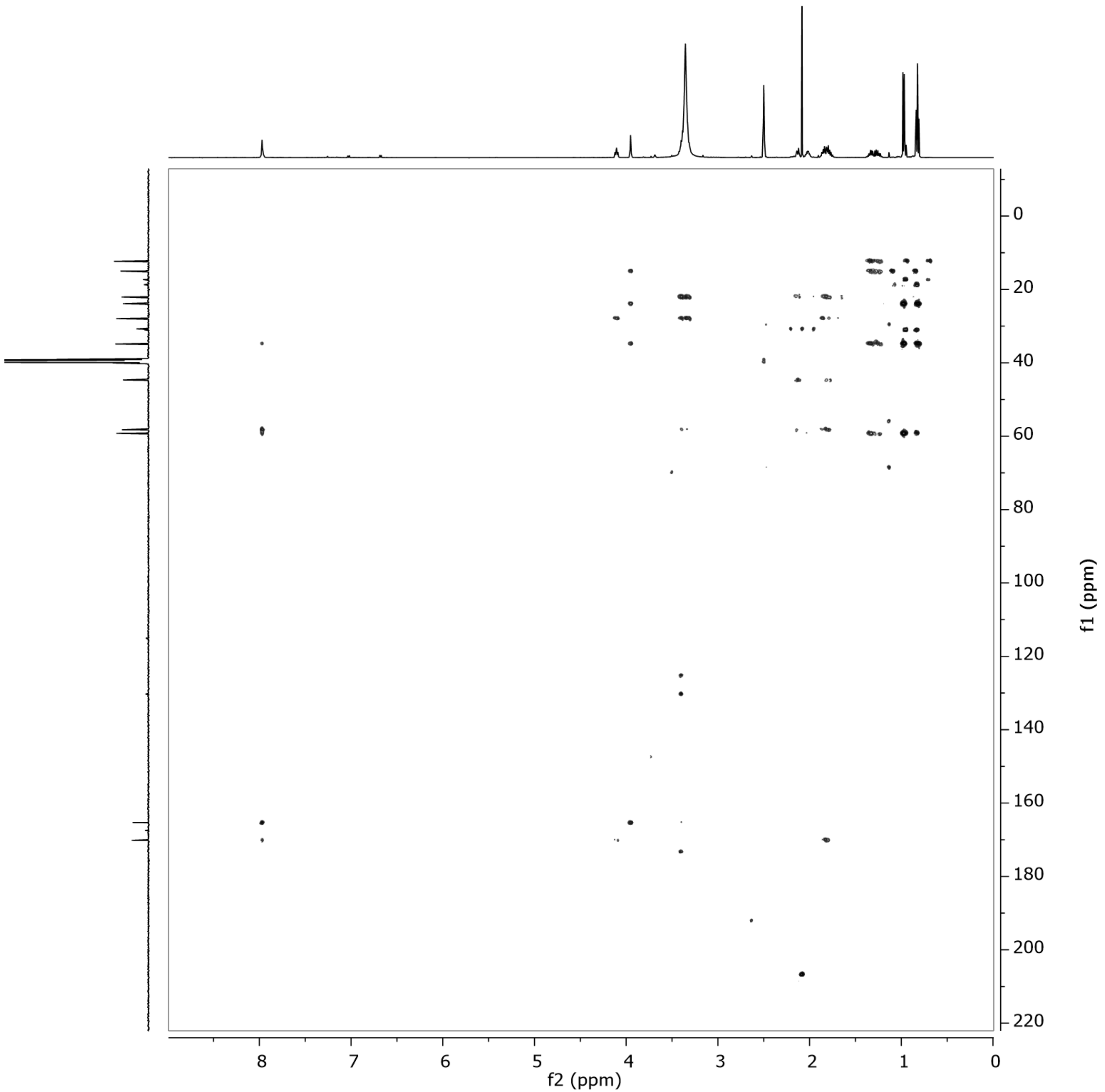


Figure S40. HMBC spectrum of **9** in DMSO-$d_6$ at 500 MHz.

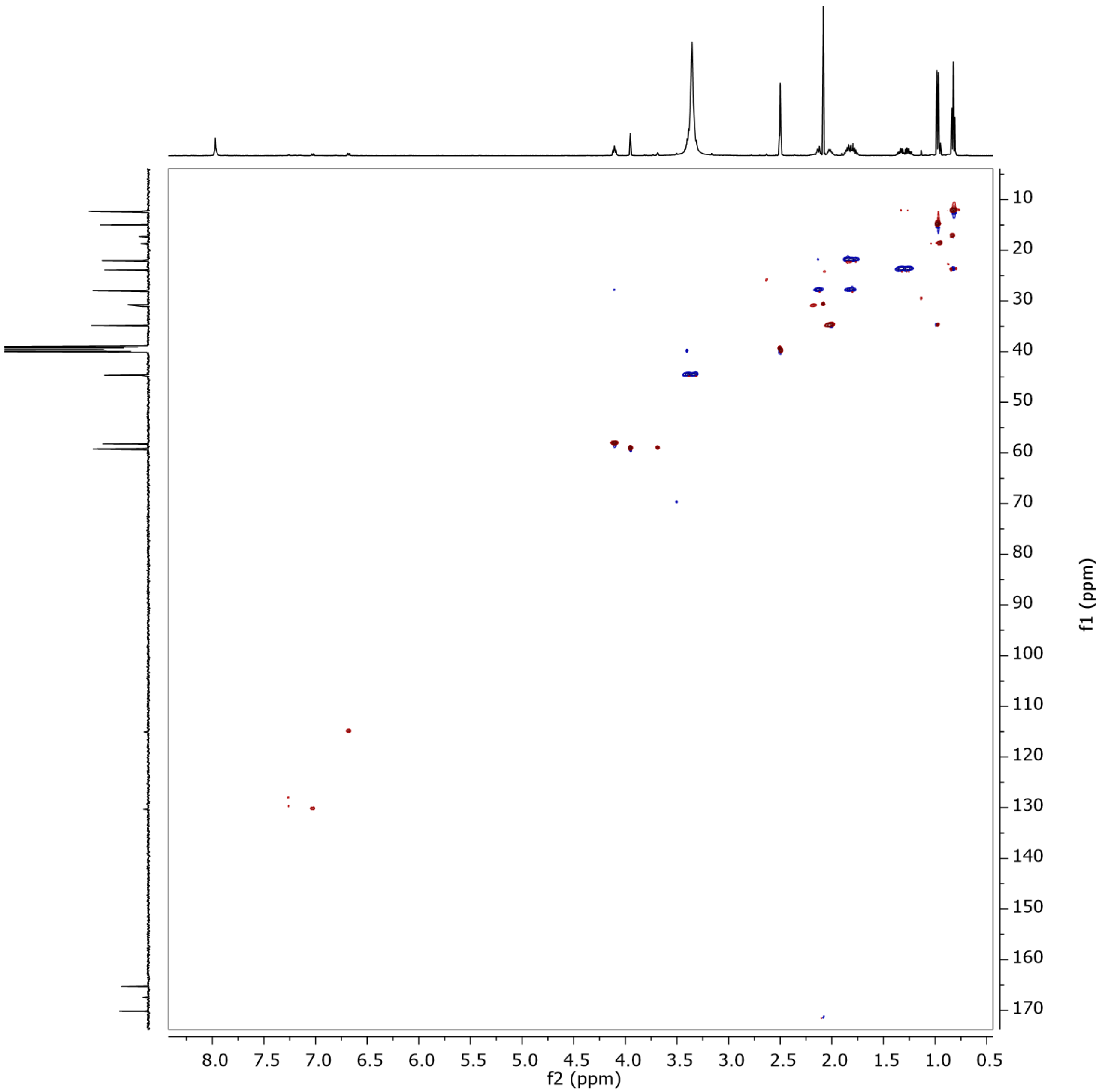


Figure S41. HSQC spectrum of **9** in DMSO-$d_6$ at 500 MHz.

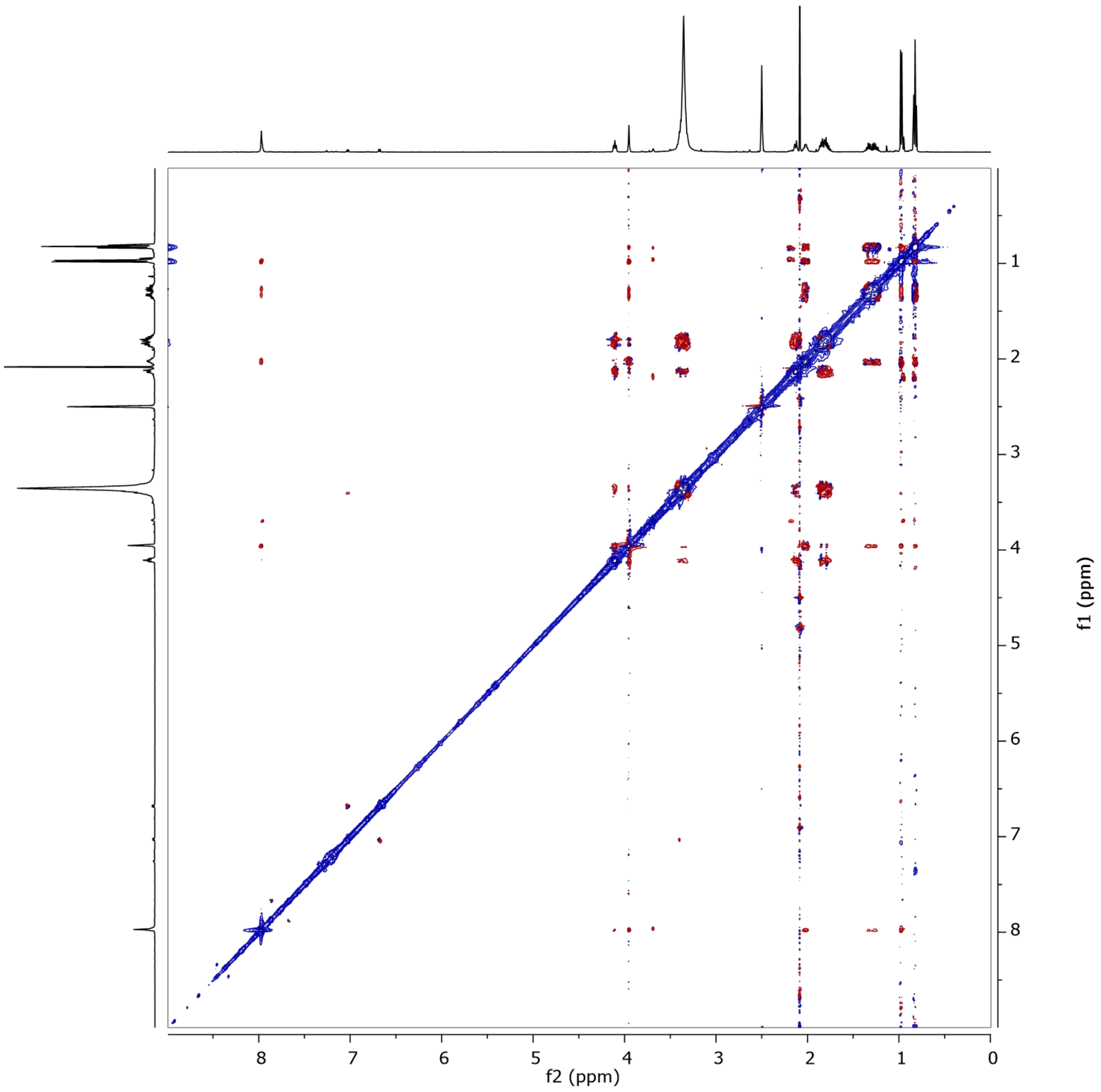


Figure S42. ROESY spectrum of **9** in DMSO-$d_6$ at 500 MHz.

### Generic Display Report

**Analysis Info**

Acquisition Date 20.06.2023 06:59:59

| | |
|---|---|
| Analysis Name | S:\DATA\AmaZon\rto20_Rita Toshe\23_06\MY04954-YM-S-F09_RC2_01_14535.d |
| Method | 14535.m |
| Sample Name | MY04954-YM-S-F09 |
| Comment | |
| Operator | lab |
| Instrument | amaZon speed |

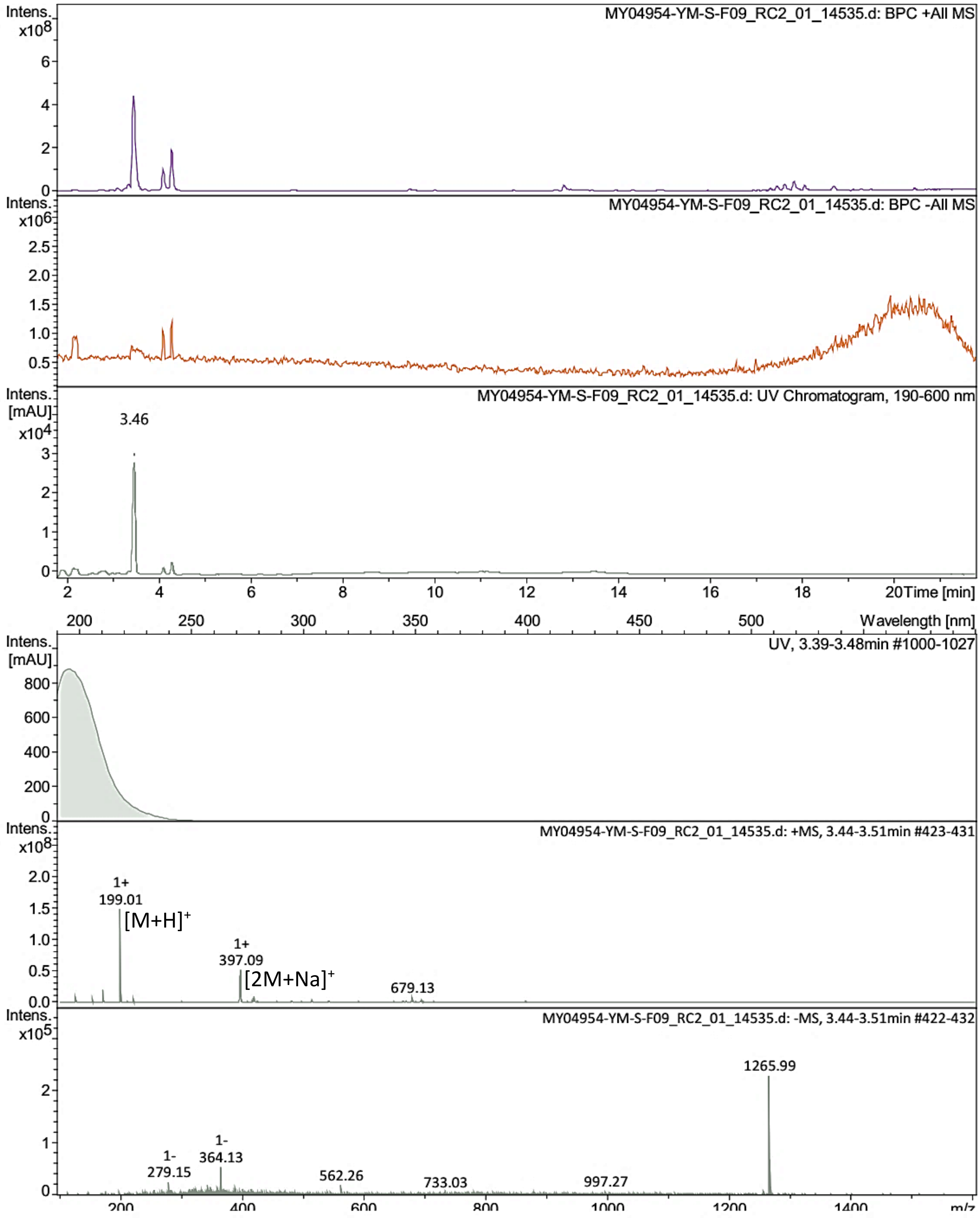


Bruker Compass DataAnalysis 4.4 printed: 21.06.2023 16:55:11 by: sel22 Page 1 of 1

Figure S43. LR-ESI-MS spectrum of **10**.

### Display Report

**Analysis Info**

| | | | |
|---|---|---|---|
| | | Acquisition Date | 27.06.2023 19:55:52 |
| Analysis Name | S:\DATA\MaXis\rto20_Rita Toshe\23_06\23_06\MY04954-YM-S-F09_18_01_12016.d | | |
| Method | pos_säure_10000_screening_ms_100_2500_line.m | Operator | ate06 |
| Sample Name | MY04954-YM-S-F09 | Instrument | maXis |
| Comment | Screening01<br>Waters Acquity UPLC BEH C_18 1,7um 2.1x50mm | | |

**Acquisition Parameter**

Ion Polarity Positive

SPS Target Mass

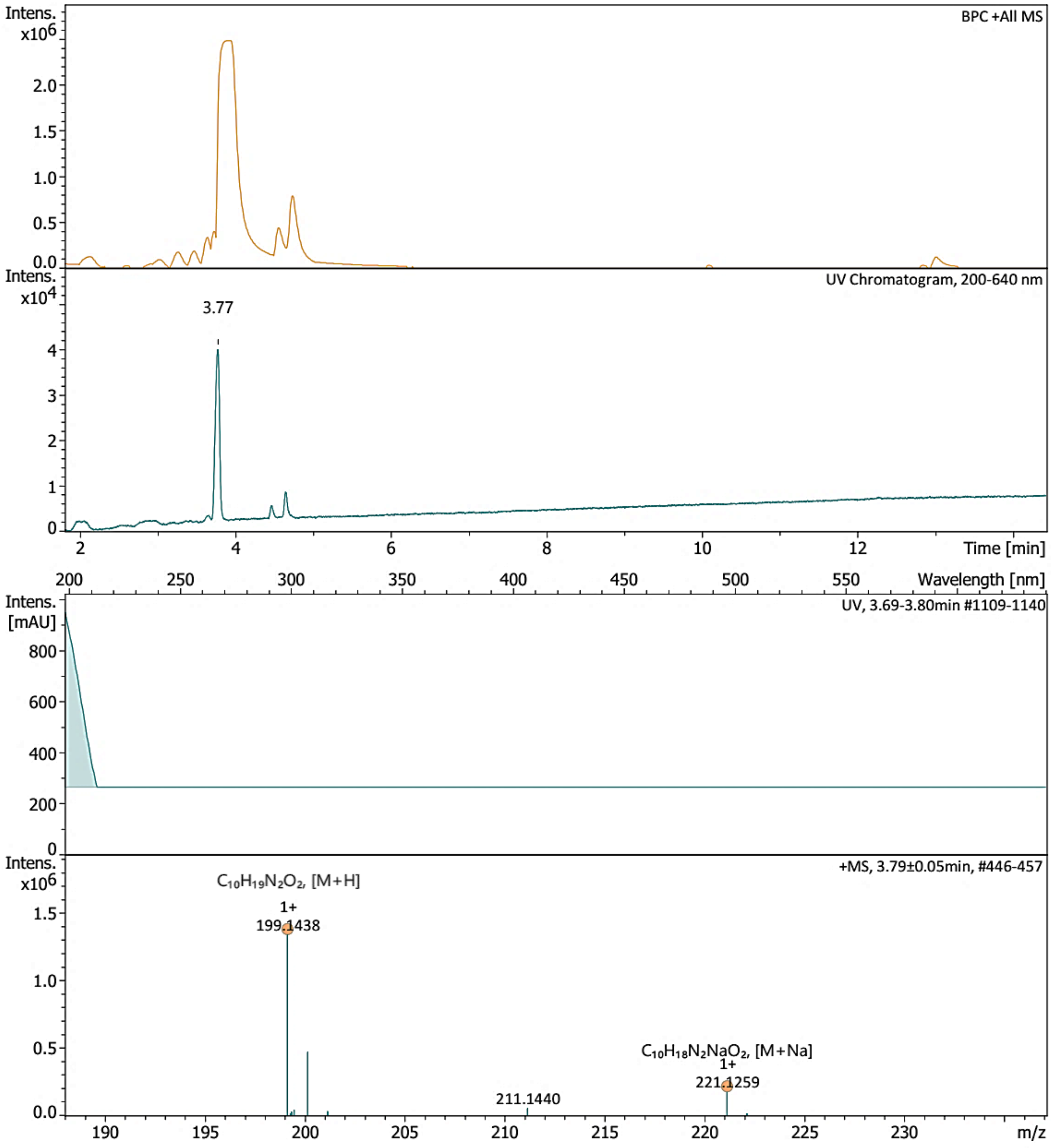


Bruker Compass DataAnalysis 6.1 printed: 28.11.2023 09:22:50 by: sel22 Page 1 of 1

Figure S44. HR-ESI-MS spectrum of **10**.

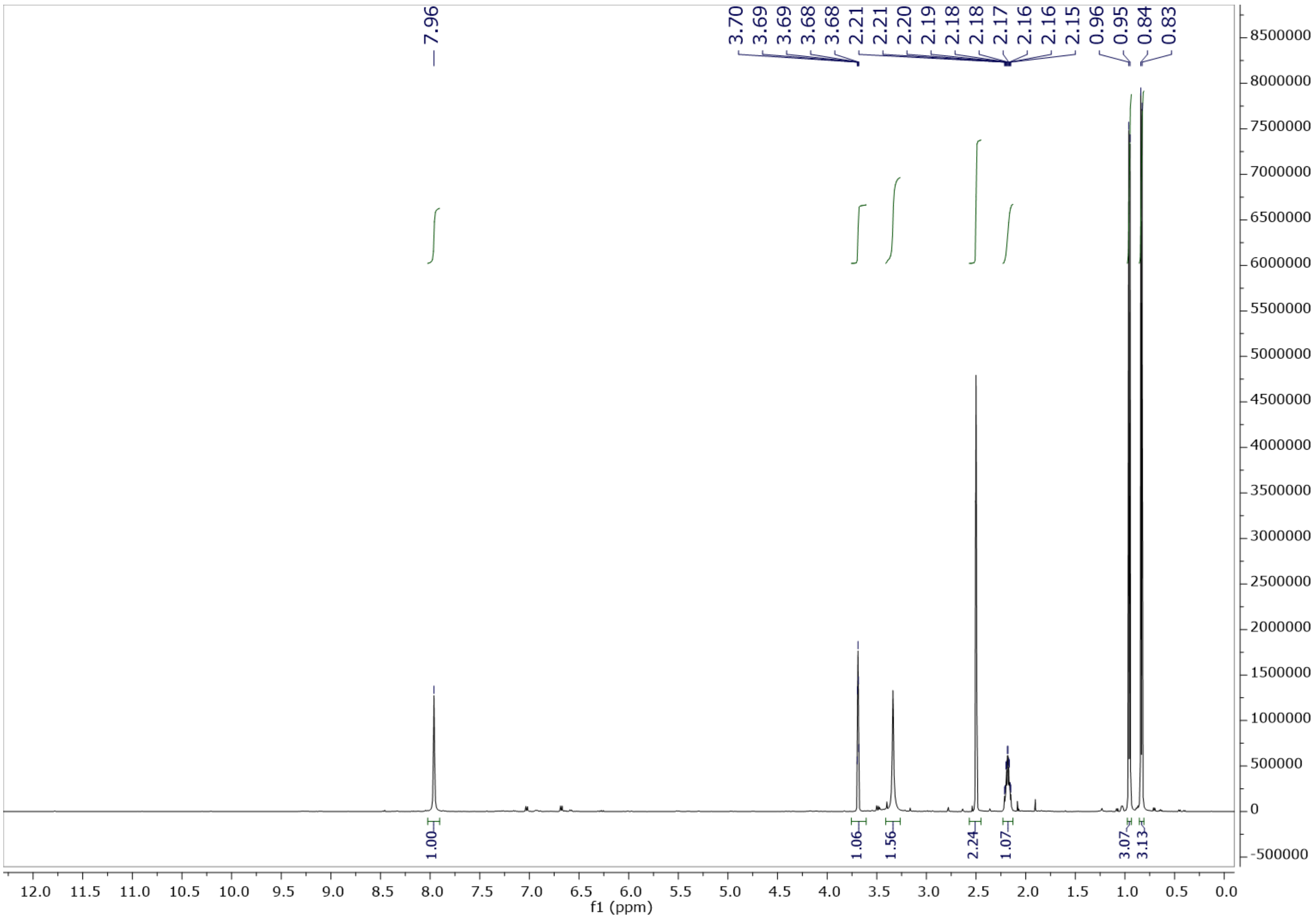


Figure S45. $^{1}$H NMR spectrum of **10** in DMSO-$d_6$ at 500 MHz.

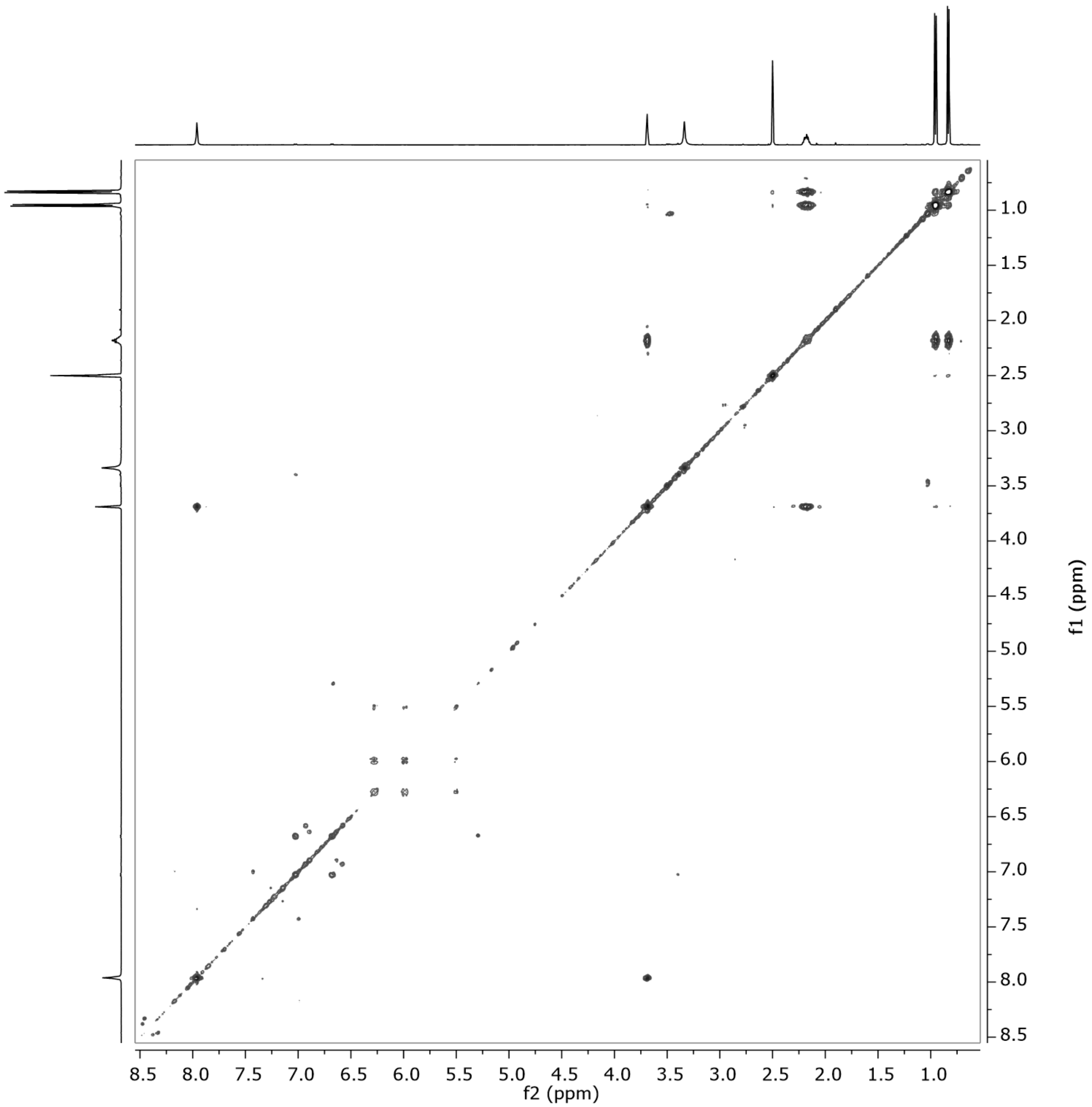


Figure S46. $^1H$–$^1H$ COSY spectrum of **10** in DMSO-$d_6$ at 500 MHz.

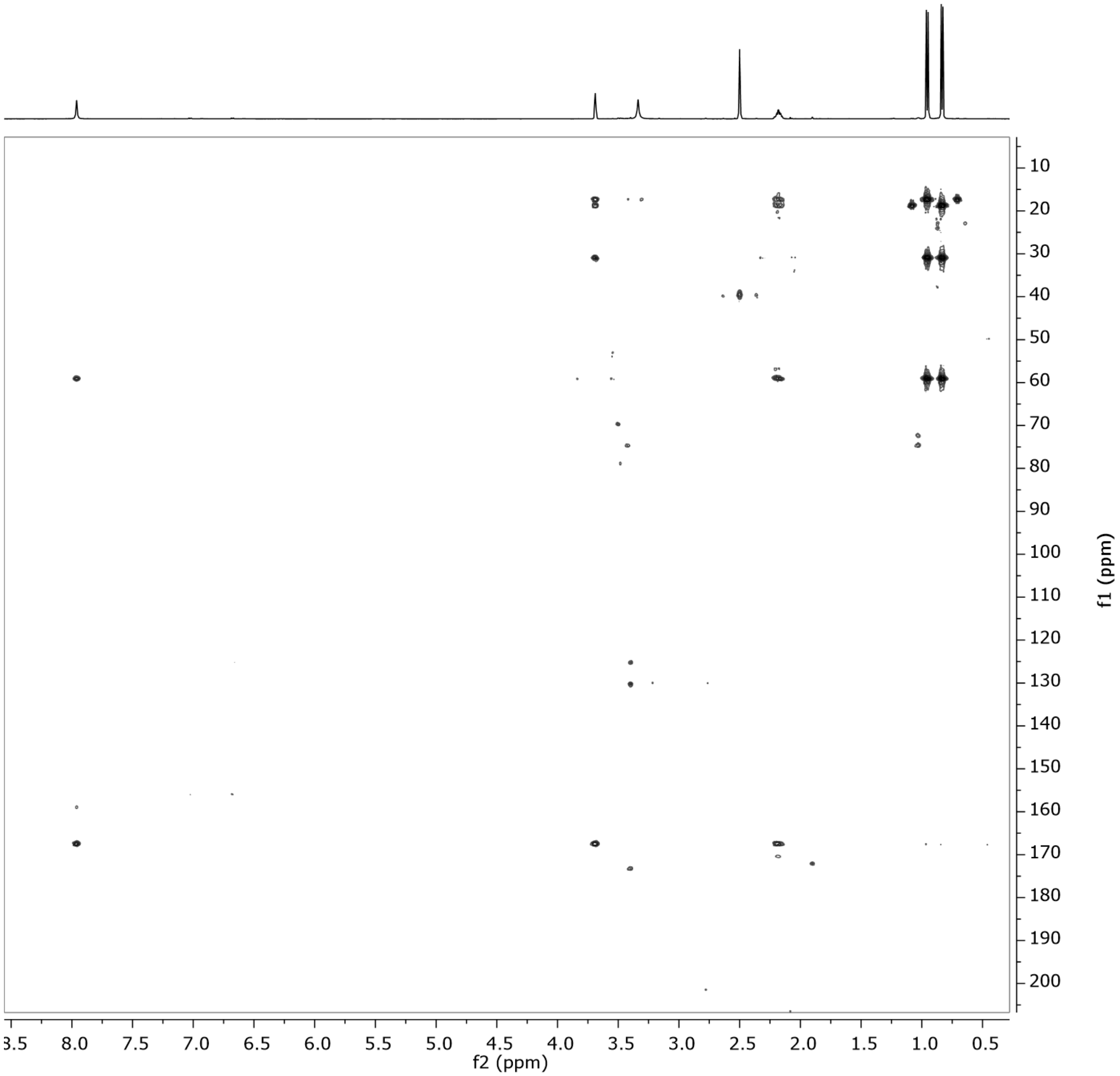


Figure S47. HMBC spectrum of **10** in DMSO-$d_6$ at 500 MHz.

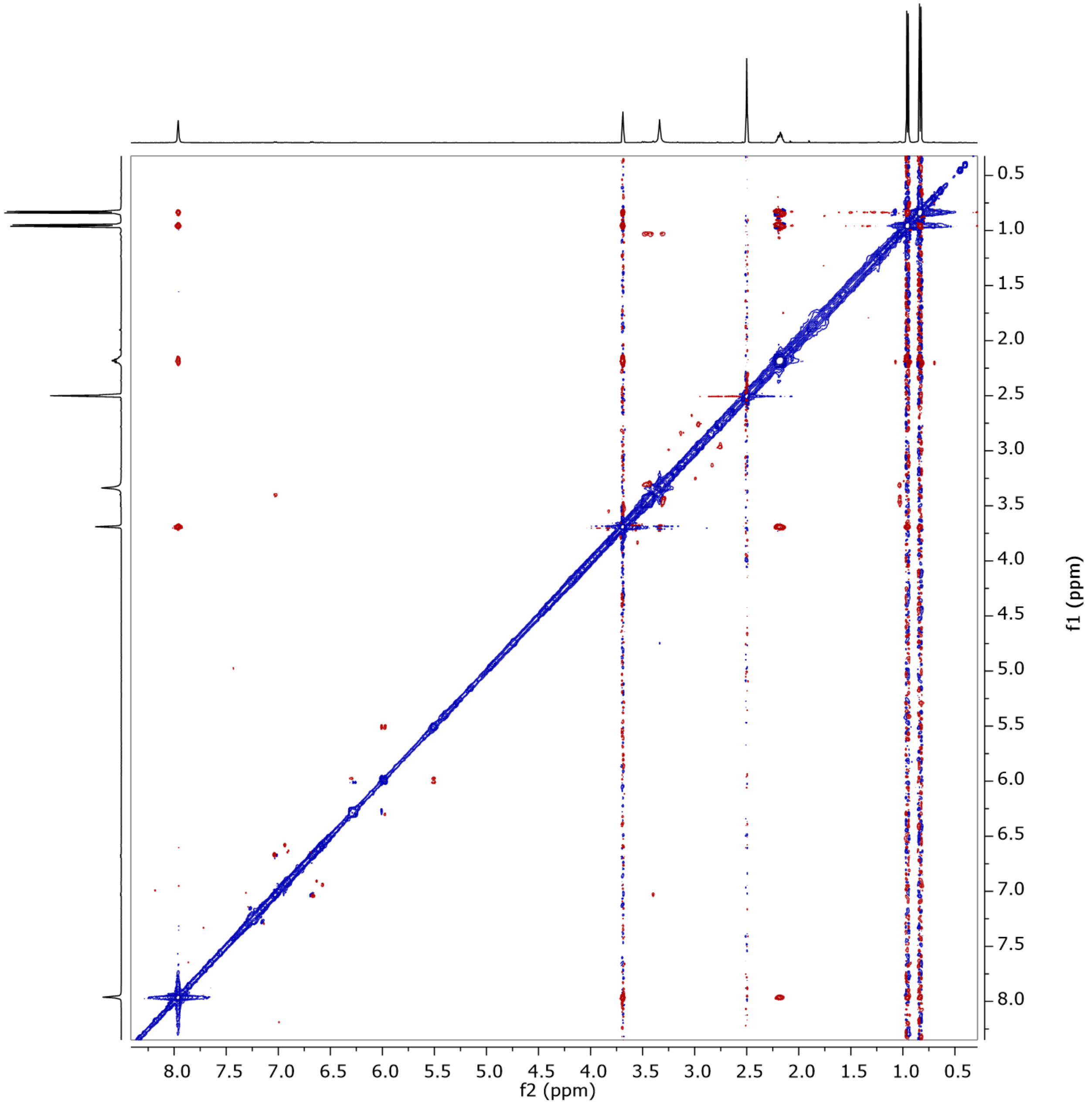


Figure S48. ROESY spectrum of **10** in DMSO-$d_6$ at 500 MHz.

### Generic Display Report

**Analysis Info**

| | | | |
|---|---|---|---|
| | | Acquisition Date | 20.06.2023 13:01:58 |
| Analysis Name | S:\DATA\AmaZon\rto20_Rita Toshe\23_06\MY04954-YM-S-F17_RD2_01_14545.d | | |
| Method | 14545.m | Operator | lab |
| Sample Name | MY04954-YM-S-F17 | Instrument | amaZon speed |
| Comment | | | |

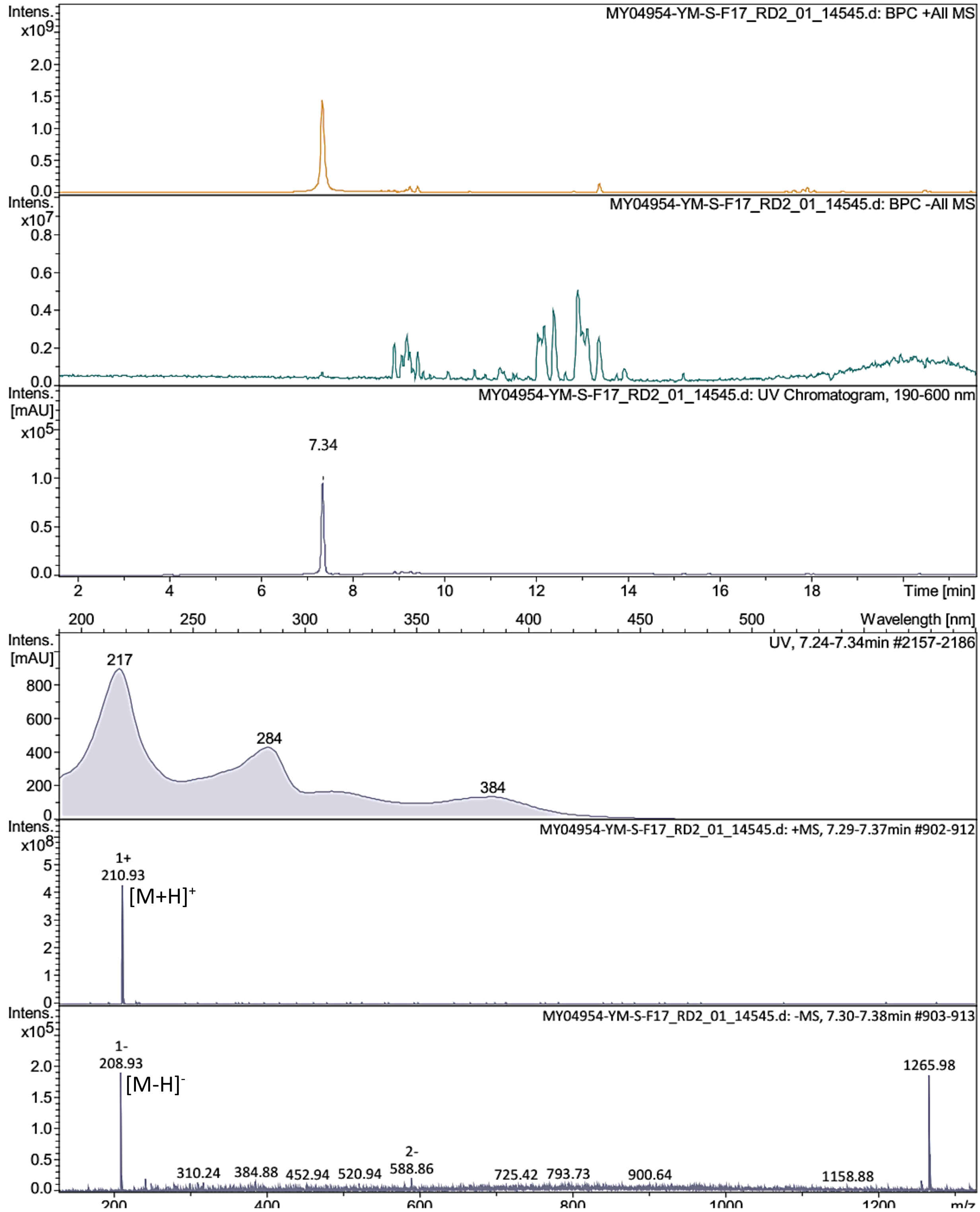


Bruker Compass DataAnalysis 4.4 printed: 21.06.2023 17:06:20 by: sel22 Page 1 of 1

Figure S49. LR-ESI-MS spectrum of **11**.

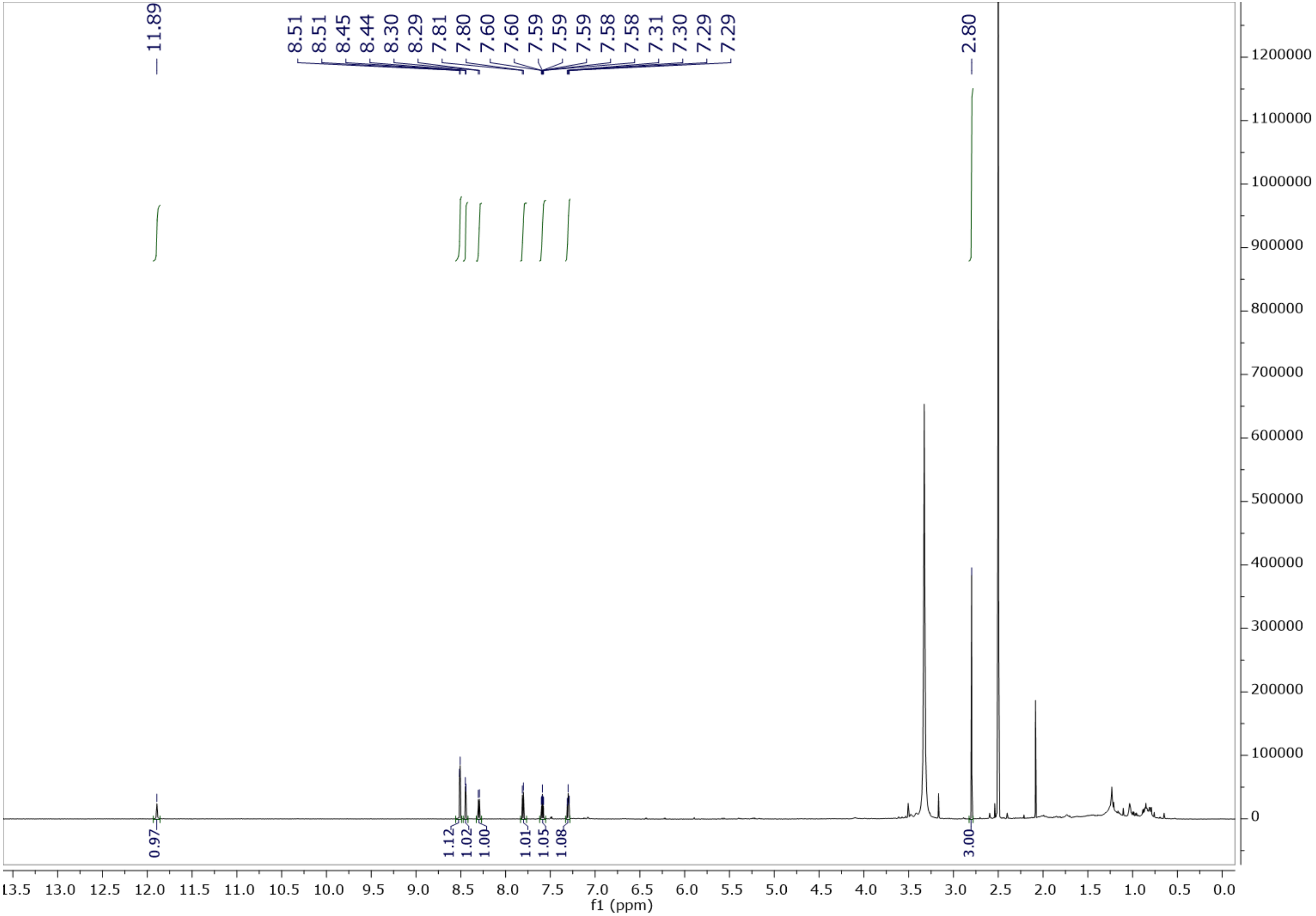


Figure S50. [1]H NMR spectrum of **11** in DMSO-$d_6$ at 500 MHz.

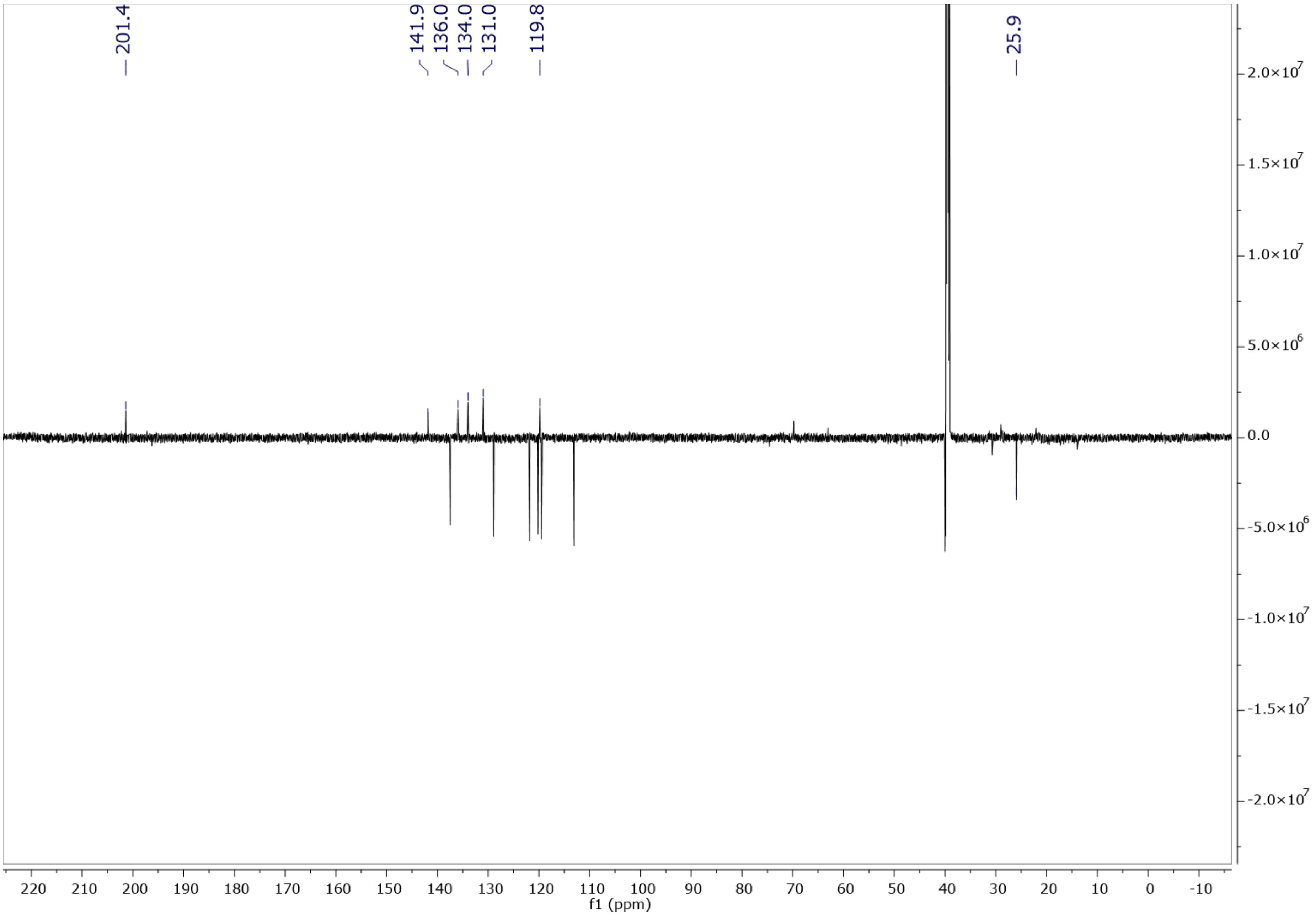


Figure S51. DEPTQ spectrum of **11** in DMSO-$d_6$ at 125 MHz.

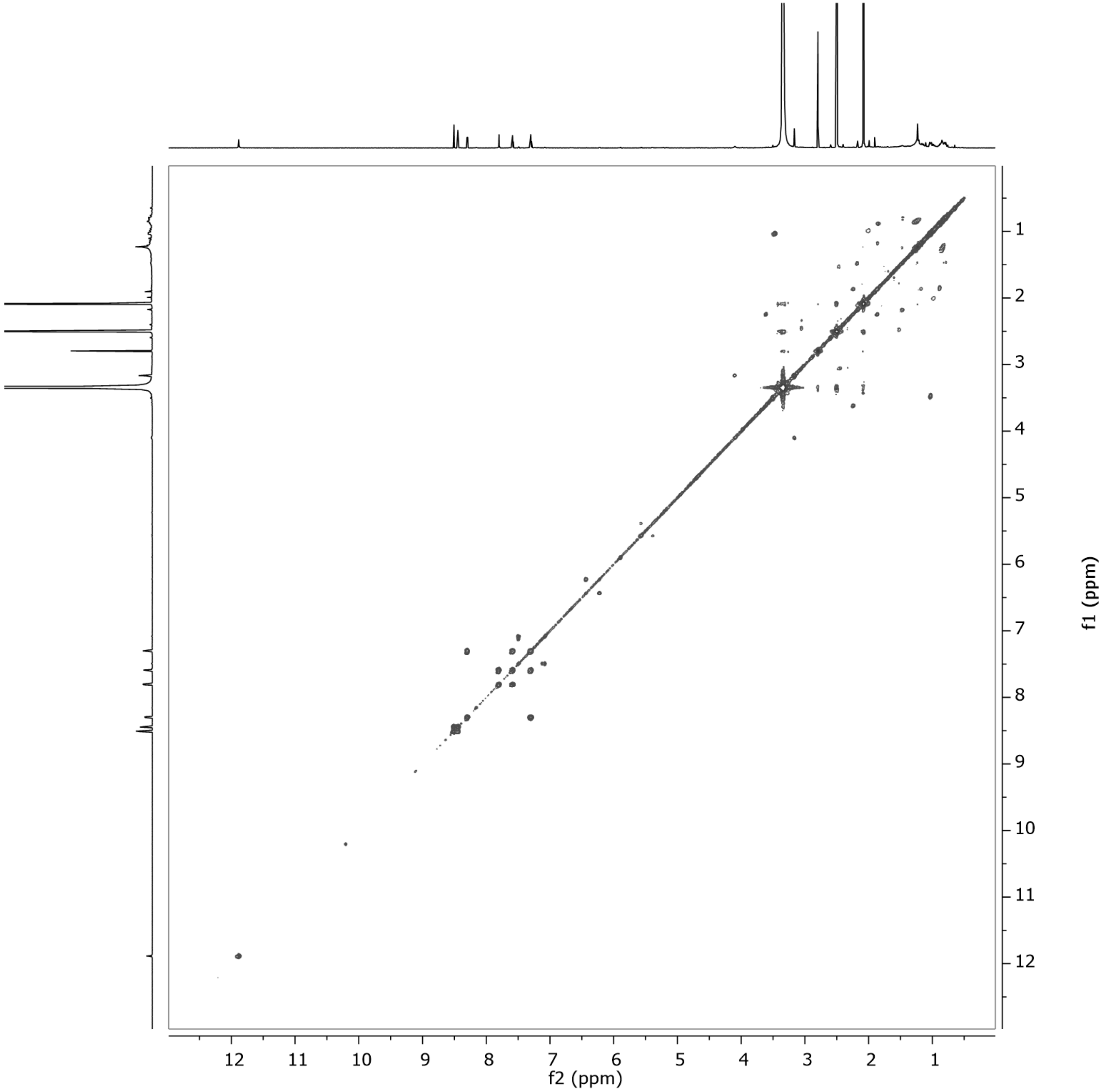


Figure S52. $^1H$–$^1H$ COSY spectrum of **11** in DMSO-$d_6$ at 500 MHz.

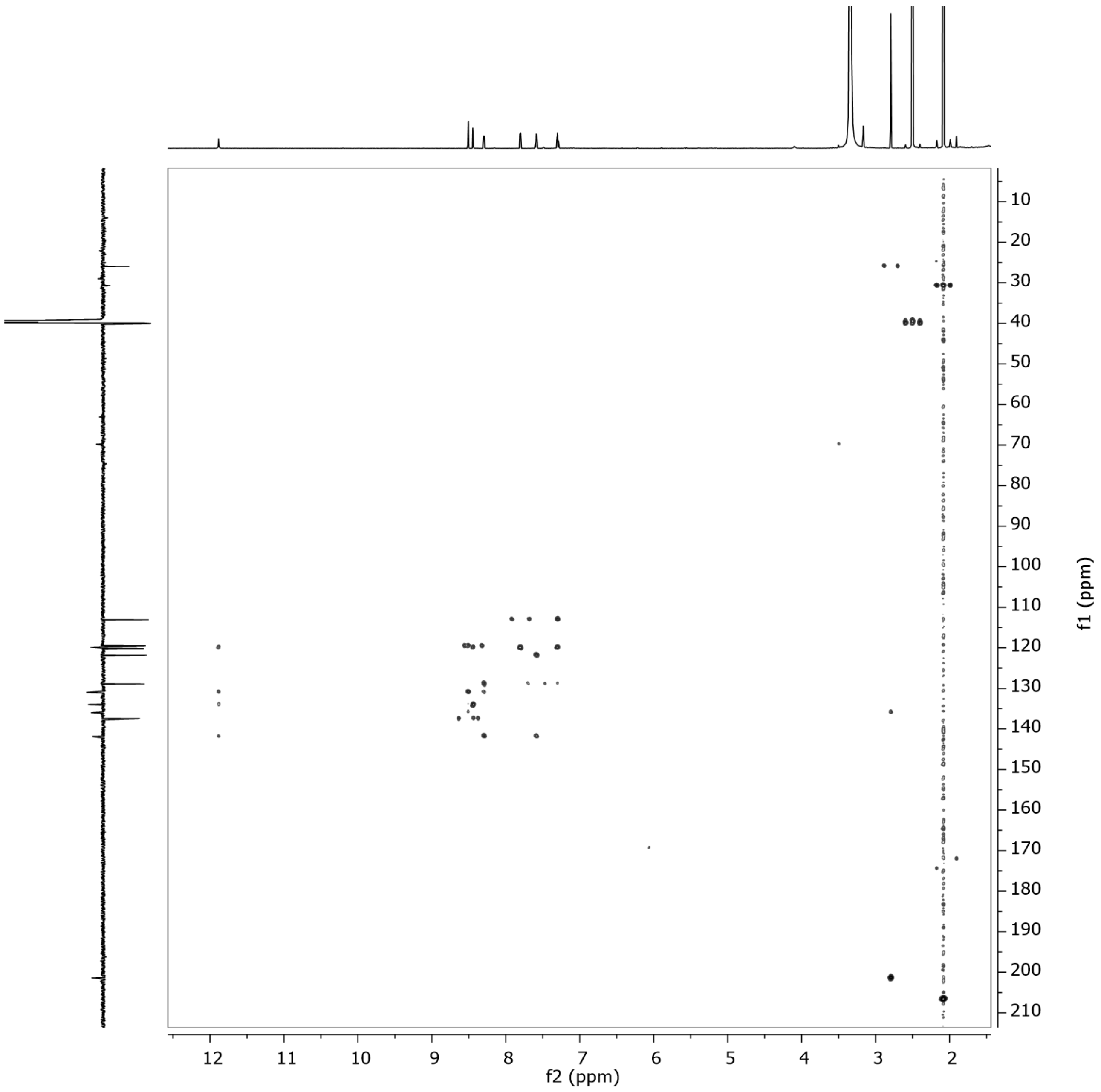


Figure S53. HMBC spectrum of **11** in DMSO-$d_6$ at 500 MHz.

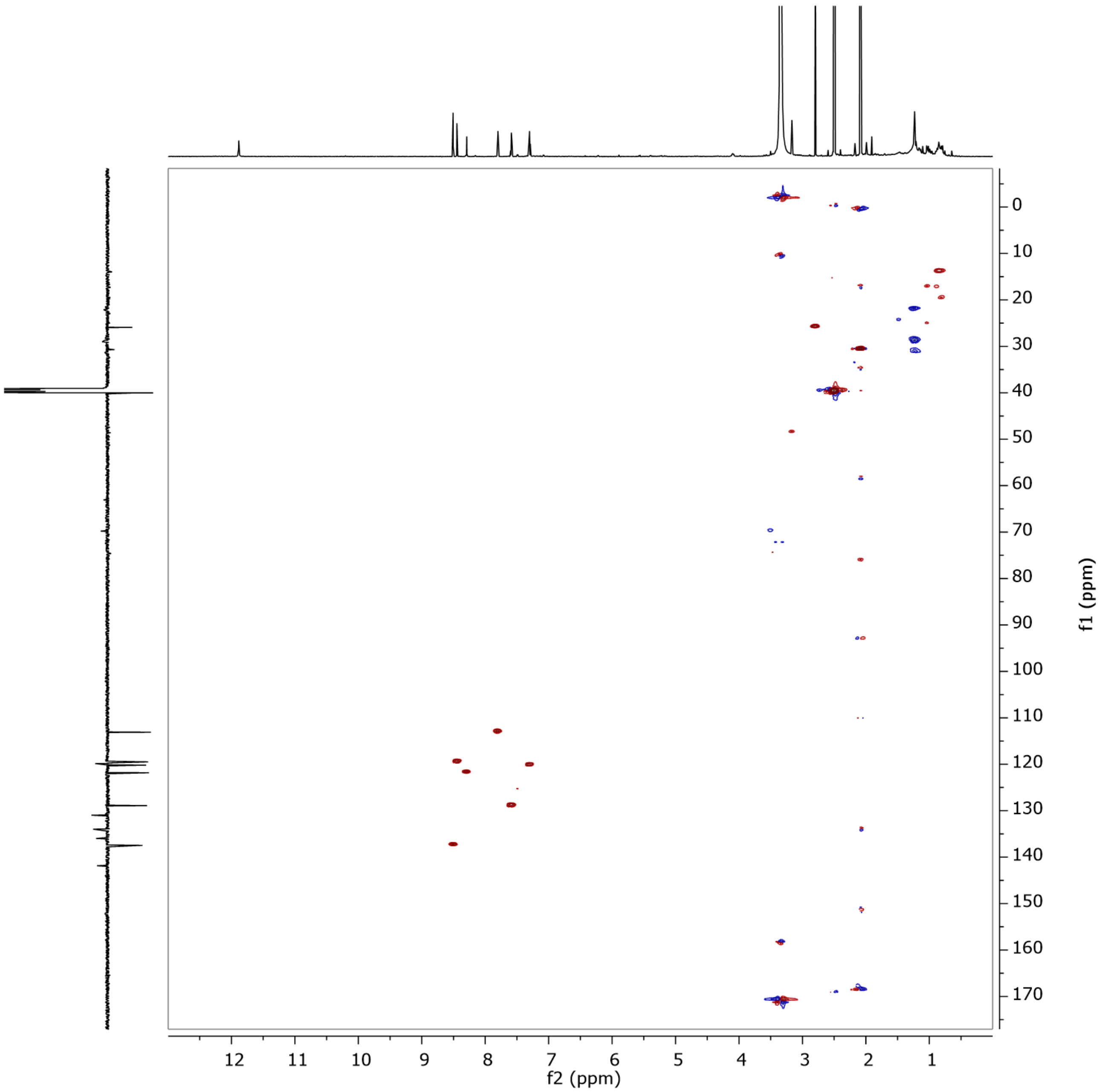


Figure S54. HSQC spectrum of **11** in DMSO-$d_6$ at 500 MHz.

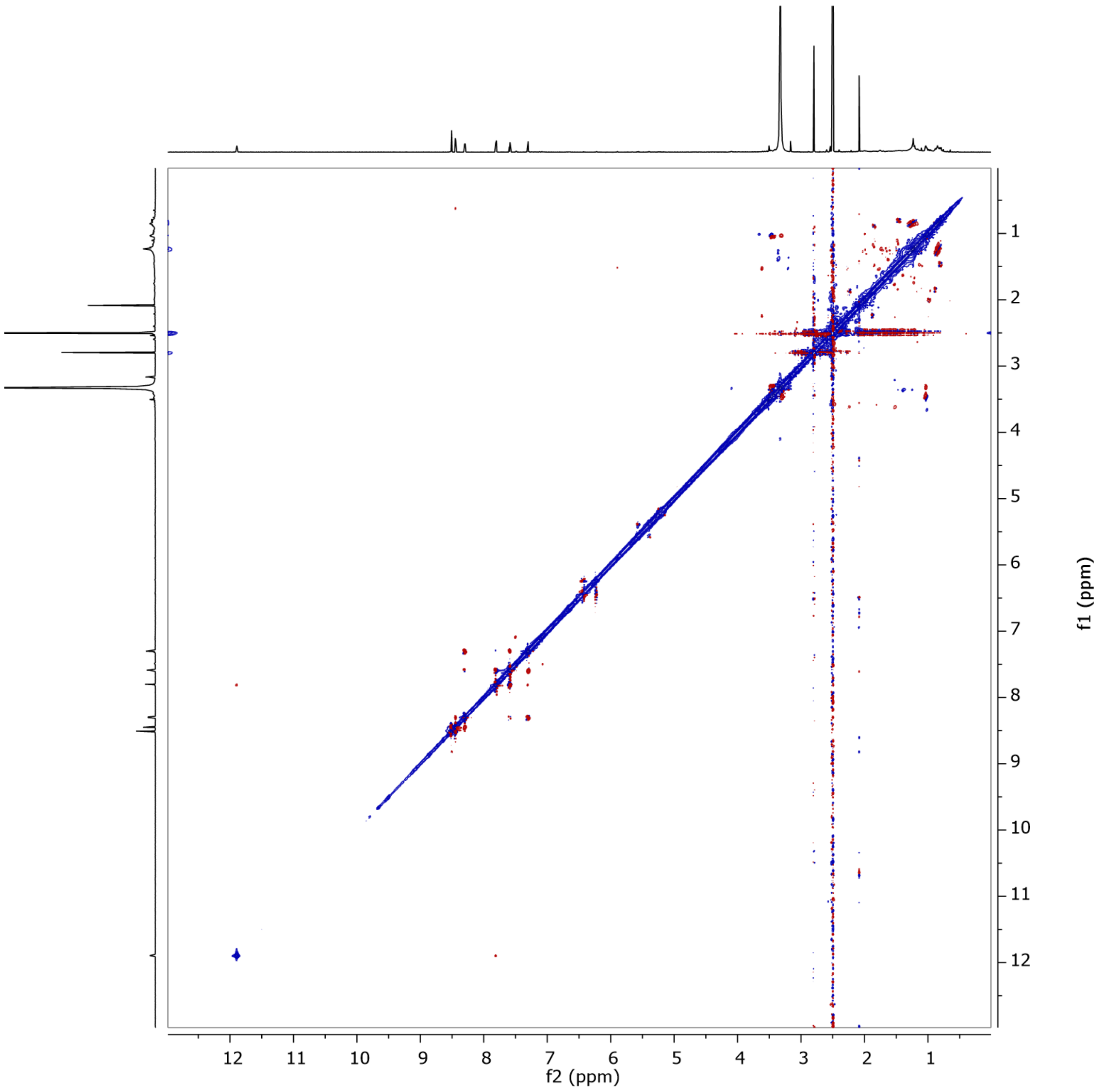


Figure S55. ROESY spectrum of **11** in DMSO-$d_6$ at 500 MHz.

### Display Report

**Analysis Info**

| | | | |
|---|---|---|---|
| Analysis Name | S:\DATA\MaXis\rto20_Rita Toshe\Hit extracts\MY04954-YM-Cordyceps blackwelliae\MY04954-YM-S-F17_23_01_12024.d | Acquisition Date | 28.06.2023 00:03:29 |
| Method | pos_säure_10000_screening_ms_100_2500_line.m | Operator | ate06 |
| Sample Name | MY04954-YM-S-F17 | Instrument | maXis |
| Comment | Screening01<br>Waters Acquity UPLC BEH C_18 1,7um 2.1x50mm | | |

**Acquisition Parameter**

Ion Polarity Positive

SPS Target Mass

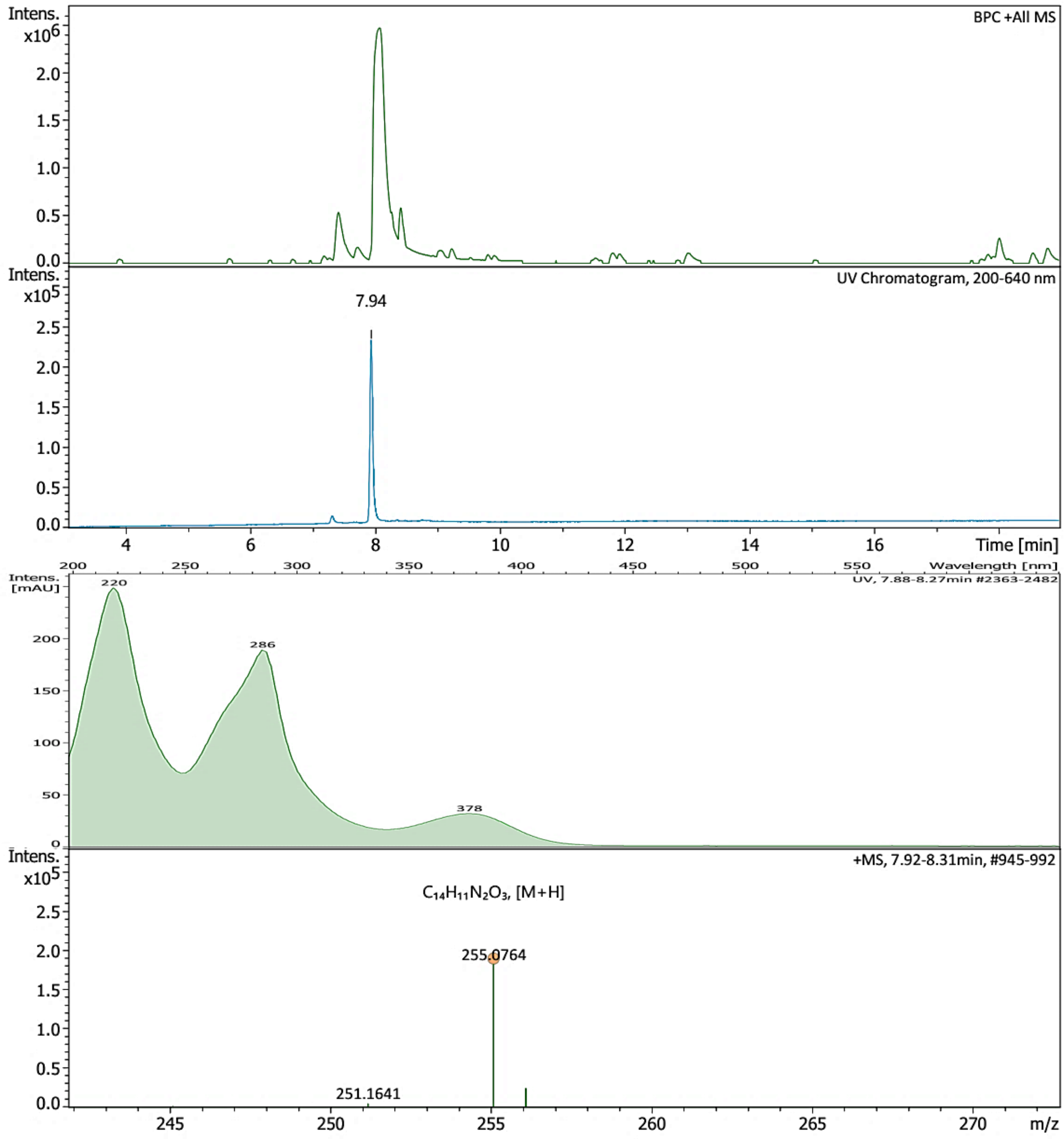


Bruker Compass DataAnalysis 6.1 printed: 15.12.2023 16:43:26 by: sel22 Page 1 of 1

Figure S56. HR-ESI-MS spectrum of **12**.

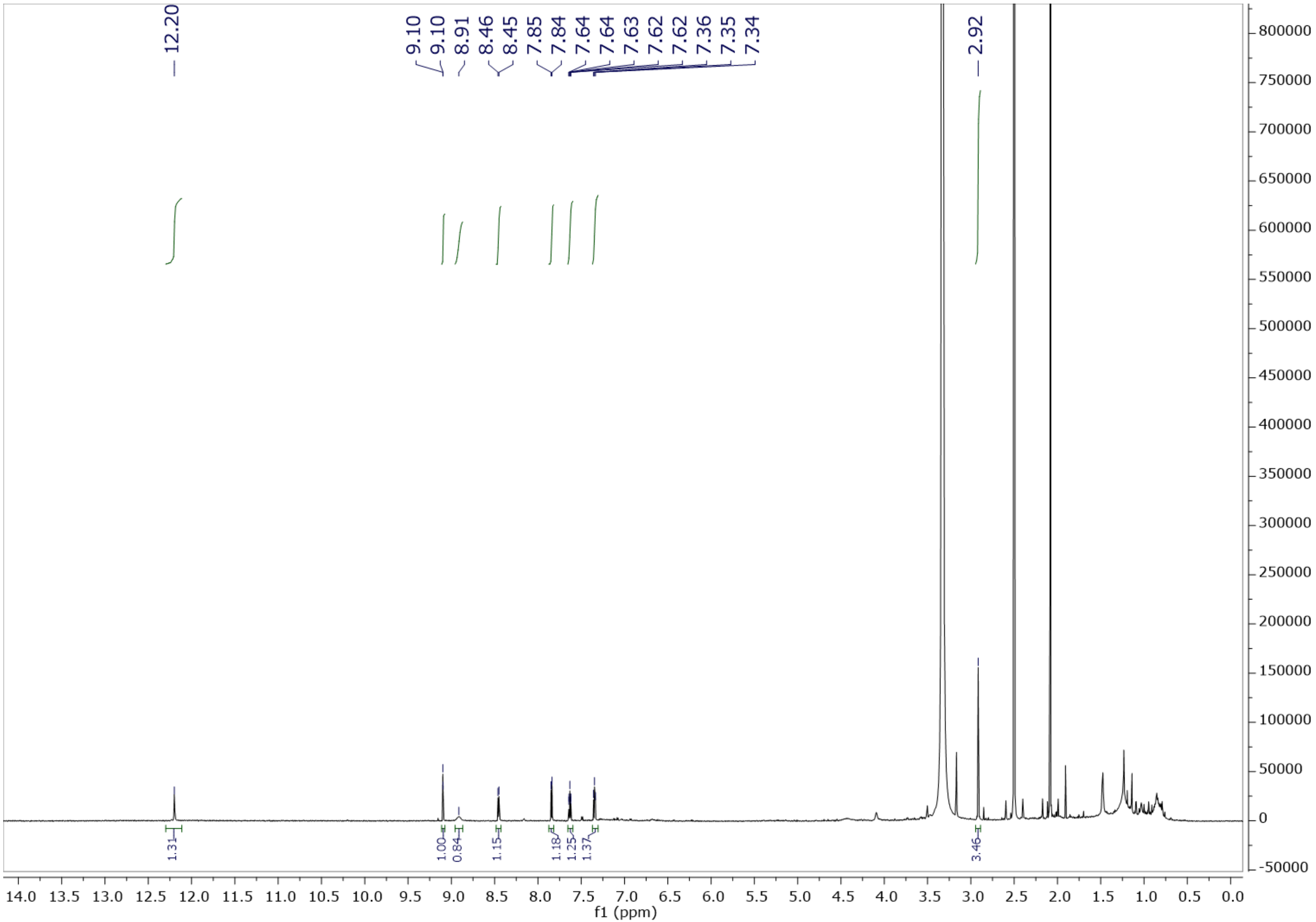


Figure S57. $^{1}H$ NMR spectrum of **12** in DMSO-$d_6$ at 500 MHz.

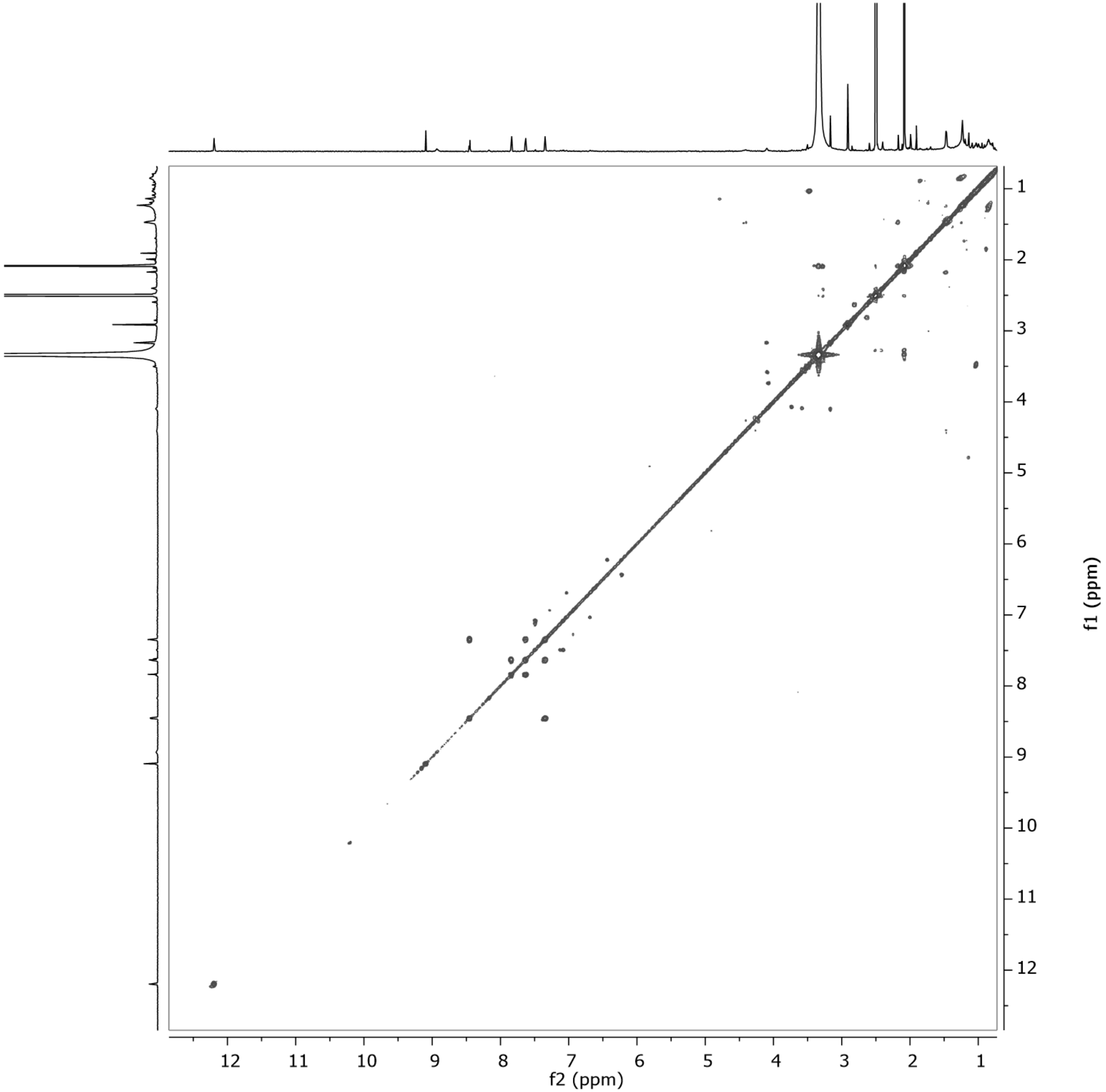

Figure S58. $^{1}H$–$^{1}H$ COSY spectrum of **12** in DMSO-$d_6$ at 500 MHz.

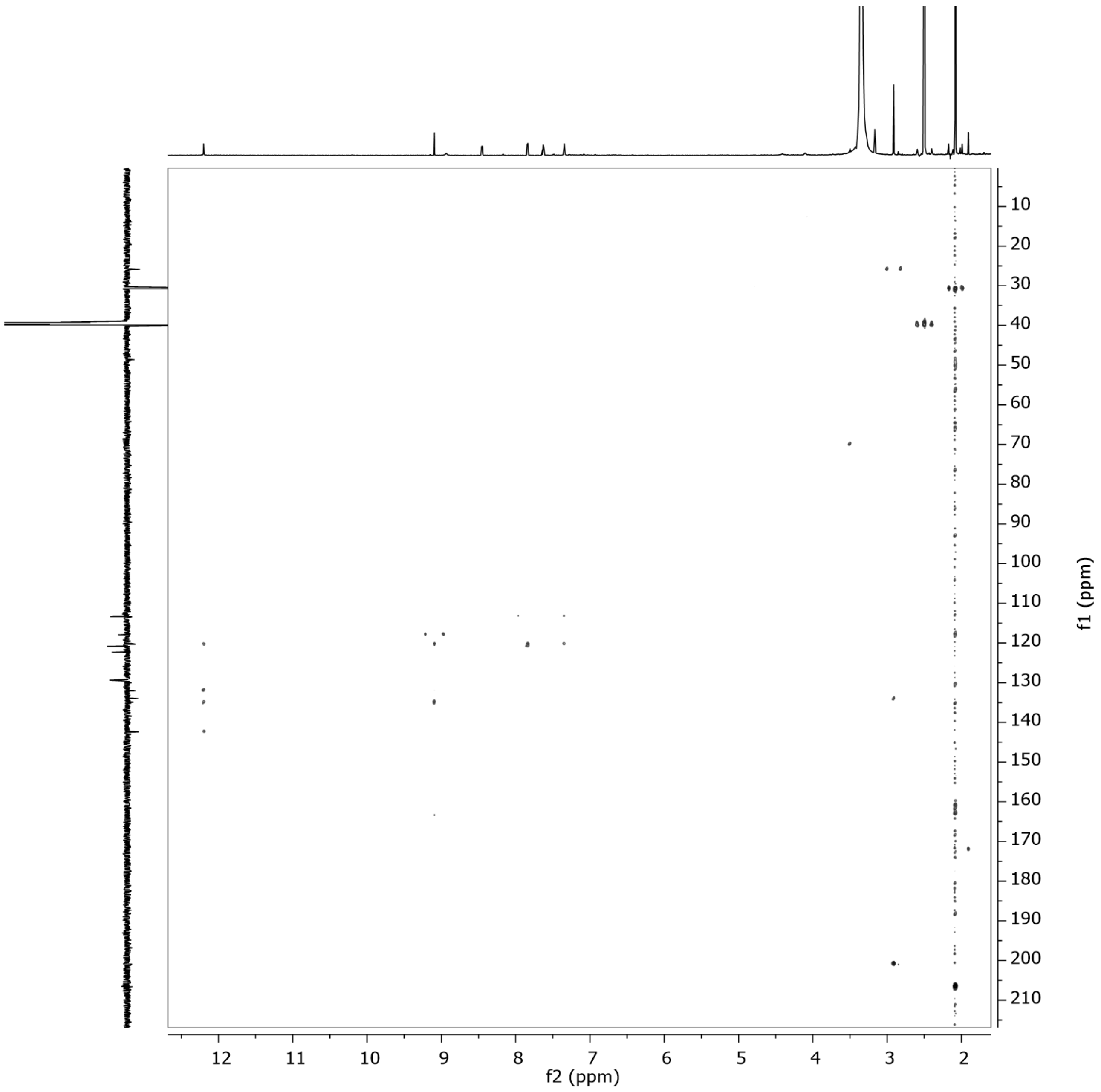


Figure S59. HMBC spectrum of **12** in DMSO-$d_6$ at 500 MHz.

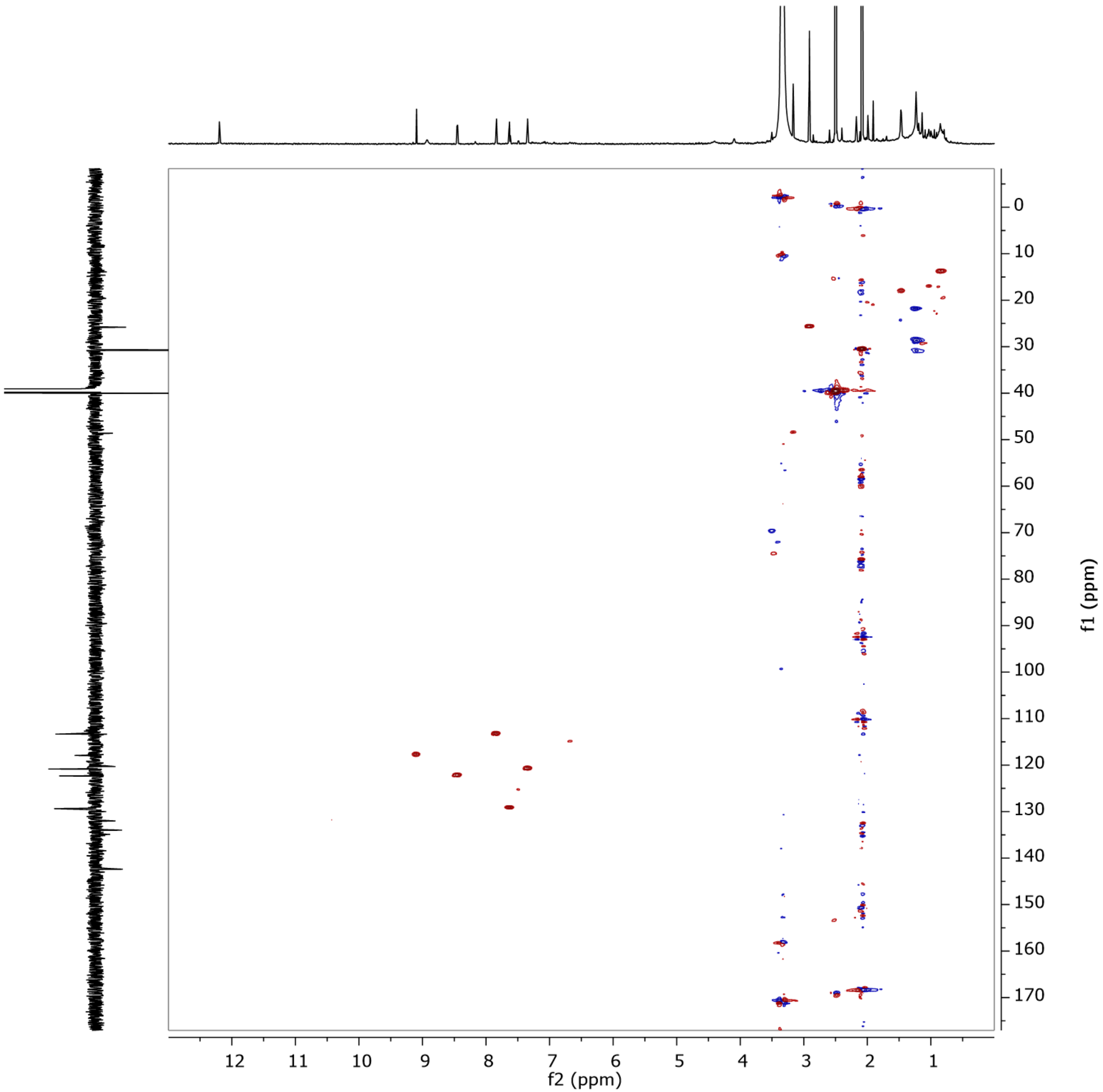


Figure S60. HSQC spectrum of **12** in DMSO-$d_6$ at 500 MHz.

Table S3. Antimicrobial properties of the isolated metabolites.

| Test Microorganism | MIC (µg/mL) | | | | | | | | | | | | Positive Control (µg/mL) |
|---|---|---|---|---|---|---|---|---|---|---|---|---|---|
| | 1 | 2 | 3 | 4 | 5 | 6 | 7 | 8 | 9 | 10 | 11 | 12 | |
| *Staphylococcus aureus* (DSM 346) | n.i. | n.i. | 66.6 | 66.6 | n.i. | n.i. | *n.i.* | n.i. | n.i. | n.i. | n.i. | n.i. | 0.21[G] |
| *Escherichia coli* (DSM 1116) | n.i. | n.i. | n.i. | n.i. | n.i. | n.i. | n.i. | n.i. | n.i. | n.i. | n.i. | n.i. | 0.42[G] |
| *Bacillus subtilis* (DSM 10) | n.i. | n.i. | n.i. | 66.6 | n.i. | n.i. | n.i. | n.i. | n.i. | n.i. | n.i. | n.i. | 16.6[O] |
| *Pseudomonas aeruginosa* (PA14) | n.i. | n.i. | n.i. | n.i. | n.i. | n.i. | n.i. | n.i. | n.i. | n.i. | n.i. | n.i. | 0.21[G] |
| *Wickerhamomyces anomalus* (DSM 6766) | n.i. | n.i. | n.d. | n.d. | n.d. | n.d. | n.i. | n.i. | n.i. | n.d. | n.d. | n.i. | 16.6[N] |
| *Candida albicans* (DSM 1665) | n.i. | n.i. | n.i. | n.i. | n.i. | n.i. | n.i. | n.i. | n.i. | n.i. | n.i. | n.i. | 8.3[N] |
| *Acinetobacter baumannii* (DSM 30008) | n.i. | n.i. | n.d. | n.d. | n.d. | n.d. | n.i. | n.i. | n.i. | n.d. | n.d. | n.i. | 0.52[C] |
| *Chromobacterium violaceum* (DSM 30191) | n.i. | n.i. | n.d. | n.d. | n.d. | n.d. | n.i. | n.i. | n.i. | n.d. | n.d. | n.i. | 1.70[G] |
| *Schizosaccharomyces pombe* (DSM 70572) | n.i. | n.i. | n.d. | n.d. | n.d. | n.d. | n.i. | n.i. | n.i. | n.d. | n.d. | n.i. | 8.30[N] |
| *Mucor hiemalis* (DSM 2656) | n.i. | n.i. | n.i. | n.i. | 66.6 | n.i. | n.i. | n.i. | n.i. | n.i. | n.i. | n.i. | 8.30[N] |
| *Rhodotorula glutinis* (DSM 10134) | n.i. | n.i. | n.d. | n.d. | n.d. | n.d. | n.i. | n.i. | n.i. | n.d. | n.d. | n.i. | 4.20[N] |
| *Mycobacterium smegmatis* (ATCC 700084) | n.i. | n.i. | n.i. | n.i. | n.i. | n.i. | n.i. | n.i. | n.i. | n.i. | n.i. | n.i. | 1.70[K] |

Table S4. Cytotoxic properties of the isolated metabolites.

| Test Cell Line | IC$_{50}$ (µM) | | | | | | | | | | | | Positive Control |
|---|---|---|---|---|---|---|---|---|---|---|---|---|---|
| | **1** | **2** | **3** | **4** | **5** | **6** | **7** | **8** | **9** | **10** | **11** | **12** | **Epothilone B (nM)** |
| Mouse fibroblast (L929) | n.a. | n.a. | n.a. | n.a. | n.a. | n.a. | n.a. | n.a. | n.a. | n.a. | n.a. | n.a. | 0.65 |
| Human endocervival adenocarcinoma (KB3.1) | n.a. | n.a. | n.a. | 13.3 | n.a. | n.a. | 36.5 | n.a. | n.a. | n.a. | n.a. | n.a. | 0.17 |
| Human prostate carcinoma (PC-3) | n.d. | n.d. | n.d. | 3.5 | n.d. | n.d. | n.d. | n.d. | n.d. | n.d. | n.d. | n.d. | 0.09 |
| Human breast adenocarcinoma (MCF-7) | n.d. | n.d. | n.d. | n.a. | n.d. | n.d. | n.d. | n.d. | n.d. | n.d. | n.d. | n.d. | 0.07 |
| Human ovarian cancer (SKOV-3) | n.d. | n.d. | n.d. | n.a. | n.d. | n.d. | n.d. | n.d. | n.d. | n.d. | n.d. | n.d. | 0.09 |
| Human epidermoid carcinoma (A431) | n.d. | n.d. | n.d. | 1.2 | n.d. | n.d. | n.d. | n.d. | n.d. | n.d. | n.d. | n.d. | 0.06 |
| Human lung carcinoma (A549) | n.d. | n.d. | n.d. | 0.9 | n.d. | n.d. | n.d. | n.d. | n.d. | n.d. | n.d. | n.d. | 0.05 |
